# Deciphering the genetic bases of canine longevity

**DOI:** 10.64898/2026.11.15.766459

**Authors:** Dimple Adiwal, Stéphanie Mottier, Armel Houel, Richard Guyon, Caroline Dufaure de Citres, Frédérique Sauvaget, Jessie Brun, Lucie Chevallier, Ambre Courtin, Grégoire Leroy, Julia Labadie, Jérôme Abadie, Marie Abitbol, Catherine André, Thomas Derrien, Benoit Hédan

**Author notes:** Food and Agriculture Organization of the United Nations (FAO), Rome, Italy. French Cani-DNA Biological Resource Centre (BRC).

## Abstract

Across dog breeds, the inverse correlation between body size and lifespan is well documented, but its genetic basis remains poorly understood, in part because size and lifespan are confounded at the breed level. Here, we assembled the Cani-DNA discovery cohort of 540 dogs from 27 breeds, with 20 dogs per breed, genotyped by low-pass whole-genome sequencing and imputation. We performed genome-wide association studies (GWAS) of breed-average body weight, breed-median lifespan and individual age at death, and tested the age-at-death loci for replication in the GRLS replication cohort of 3,024 Golden Retrievers. The GWAS of breed-average body weight recovered established body-size loci, confirming that the cohort could detect known associations. Of the 69 breed-weight loci, 30 were shared with breed-median lifespan, and at all of them the allele increasing body weight was associated with shorter lifespan. At the individual level, age at death, with breed-average body weight as a covariate, identified loci distinct from all breed-level loci. Three of them, the *CDH4*, *TSHZ3* and *TSC22D2* loci, replicated in the GRLS replication cohort. In humans, *CDH4* and *TSHZ3* have been implicated in longevity, and the region orthologous to the *TSC22D2* locus has been associated with markers of biological ageing. Together, these results identify genetic variation associated with the lifespan of individual dogs beyond breed body size, and support the dog as a model in which genetic leads on ageing can be identified.

## Introduction

Artificial selection has produced dog breeds in which the largest weigh almost a hundred times more than the smallest, and the longest-lived live at least twice as long as the shortest-lived (1,2). Across breeds, heavier dogs live shorter lives than lighter ones (1,3). In a study of more than 580,000 dogs in the United Kingdom, median life expectancy ranged from 5.4 years in the Caucasian Shepherd Dog to 15.4 years in the Lancashire Heeler, and large breeds had a shorter median lifespan than small breeds (2). This inverse relationship runs counter to the trend observed at the species level, where larger mammal species tend to live longer than smaller ones (1). Part of the shorter lifespan of heavier breeds may reflect cancer, since these breeds have higher cancer mortality (4,5). The two traits also share part of their genetic basis. A single *IGF1* haplotype is a major determinant of small size in dogs (6), and the *IGF1* region has also been associated with breed lifespan in dogs (7). This is consistent with the role of insulin/IGF-1 signalling in lifespan across species, since reducing this signalling extends lifespan in worms (8) and mice (9), and variants reducing IGF-1 receptor activity are enriched in human centenarians (10). Across breeds, lifespan is therefore closely tied to body size, both phenotypically and genetically.

Dogs are also a valuable model for studying the genetics of ageing. Companion dogs live in the same environment as their owners, and causes of death such as cancer and metabolic disease follow similar age patterns in dogs and in humans (11). Their breed structure also favours genetic mapping. Each breed is a closed population, in which linkage disequilibrium extends up to a hundred times further than in humans, so association studies in dogs require far fewer markers (12). In addition, variants of large effect, such as *IGFI* haplotype that determines small size, differ markedly in frequency between breeds (6,13–15). As a result, loci of large effect can be mapped in dog cohorts far smaller than those required in human studies (16).

This approach has been applied to lifespan, but mostly at the level of breeds. Genome-wide studies of dog longevity have tested associations with breed-average or breed-median lifespan (4,7,16–18). Because each breed contributes a single lifespan value, such analyses detect differences between breeds (16), and because breed lifespan is tied to body size, their loci are difficult to separate from those affecting size. Other studies have analysed the lifespan of individual dogs, but within a single breed, comparing long- and short-lived Golden Retrievers (19) or Cavalier King Charles Spaniels (20). However, lifespan varies widely between dogs of the same breed and between breeds of similar size (2). To our knowledge, no study has yet mapped the variation in lifespan between individual dogs across many breeds independently of breed body size.

Identifying variants associated with individual lifespan independently of body size has both veterinary and translational value. Genetic tests and genomic selection are already used to improve the health of pedigree breeds while preserving the characteristics that define them (21), and variants associated with lifespan could extend this approach to longevity. They could also help to address questions that are difficult to study in humans. Long-lived individuals carry as many disease-risk alleles as younger people (22), so their longevity may depend on protective alleles that buffer the effects of disease (23). Such alleles are difficult to identify, because cohorts of exceptionally long-lived people are small and their findings are difficult to replicate (24). A similar pattern may exist in dogs. In Golden Retrievers, a breed with a cancer-related mortality of up to 65% (25), a variant in *ERBB4* was associated with longer lifespan in two separate populations (19). Whether such protective alleles exist across breeds, and whether they act on specific diseases, remains to be understood. Identifying them could reveal protective mechanisms relevant to human ageing.

Here, we conducted genome-wide association studies (GWAS) of body size and longevity in the Cani-DNA discovery cohort, comprising 540 dogs from 27 breeds genotyped by low-pass whole-genome sequencing and imputation. We first ran a GWAS of breed-average body weight as a positive control for the statistical power of our cohort. We then performed GWAS of breed-median lifespan, with and without breed-average body weight as a covariate, to distinguish loci shared with body weight from loci specific to lifespan. Next, we analysed individual longevity, as age at death and as old/young status, both with breed-average body weight as a covariate. Finally, we tested the age-at-death loci for replication in the GRLS replication cohort, an independent cohort of 3,024 Golden Retrievers. Together, these analyses identify three loci associated with individual age at death independently of breed-average body weight and replicated in an independent cohort, all of which harbour genes previously linked to human longevity or ageing.

## Materials and Methods

### Ethics and cohort design

This study was approved by the Comité Rennais d’Éthique en matière d’Expérimentation Animale (CREEA, registered as CEEA no. 007; approval no. CE07-2020-11-CA). Blood or buccal swab samples were collected from privately owned dogs in France during routine veterinary care, with the owners’ informed consent, through the veterinary network of the Cani-DNA Biological Resource Centre (https://igdr.univ-rennes.fr/crb-cani-dna). Dates of birth were taken from the French studbook (Livre des Origines Français), and dates of death from the Cani-DNA records, updated with the French national register of identified dogs (I-CAD, https://www.i-cad.fr).

Twenty-seven breeds were selected among those most represented in the Cani-DNA biobank with a recorded date of death, so as to span a wide range of breed-average body weight (3 to 66 kg) and breed-median lifespan (5 to 14 years). They represent 14 of the 23 phylogenetic clades of the domestic dog defined by Parker et al. (14) and 9 of the 10 breed groups of the Fédération Cynologique Internationale (Table S1).

Dogs were selected on their age at death, in a case–control design matched by breed. In each breed, ten old dogs were chosen among the oldest available, targeting the 90th percentile of the breed’s age at death, and ten young dogs among those that died at or around the breed-median lifespan. Breed percentiles were computed from Société Centrale Canine (SCC) mortality records. Because the number of available dogs differed between breeds, the old dogs ranged from below the 80th to above the 99th percentile, with 53% at or above the 90th (Table S1). In every breed, the youngest old dog was older than the oldest young dog, by at least 1.7 years. After quality control, the cohort comprised 540 dogs, 270 old and 270 young, hereafter the Cani-DNA discovery cohort.

### Phenotypes

Longevity was analysed at two levels, that of the individual dog and that of the breed (Table 1). At the individual level, age at death, in years, was computed from the dates of birth and death, and old/young status was the case–control status defined above. At the breed level, breed-average body weight was the midpoint of the adult weight range in the American Kennel Club breed standard, and breed-median lifespan was the median age at death of the breed in the SCC mortality records. The relationship between breed-average body weight and breed-median lifespan across the 27 breeds was assessed by Pearson correlation and linear regression.

**Table 1.** Phenotypes and covariates analysed in the Cani-DNA discovery cohort (540 dogs, 27 breeds).

| Variable | Level | Definition | Source |
|---|---|---|---|
| Breed | Breed | Breed of the dog | Cani-DNA records |
| Sex | Individual | Male or female | Cani-DNA records |
| Age at death | Individual | Age at death, in years, from the dates of birth and death | French studbook (LOF); Cani-DNA records updated with I-CAD |
| Old/young status | Individual | Case (old): among the oldest dogs of its breed, targeting the 90th percentile of breed age at death. Control (young): died at or around the breed-median lifespan | Derived from age at death and SCC mortality records |
| Breed-average body weight | Breed | Midpoint of the adult weight range in the breed standard, in kg | American Kennel Club breed standards |
| Breed-median lifespan | Breed | Median age at death of the breed, in years | SCC mortality records |
| Weight-corrected age at death | Individual | Age at death − (12.99 − 0.08842 × breed-average body weight), in years; positive values indicate a dog that lived longer than expected for its size | Calculated from da Silva and Cross (26) |

To compare dogs of breeds with different expected lifespans, age at death was also expressed relative to the lifespan expected from breed-average body weight, using the phylogenetic regression of breed-median lifespan on adult body weight of da Silva and Cross (26). This weight-corrected age at death was computed as age at death − (12.99 − 0.08842 × breed-average body weight), so that positive values indicate a dog that lived longer than expected for its size. Data for each of the 540 dogs are provided in Table S1.

### Sequencing, imputation and quality control

Genomic DNA was extracted with the NucleoSpin 96 Tissue kit (Macherey-Nagel). Low-pass whole-genome sequencing and genotype imputation were performed by Gencove company (New York, USA), following established methodology for low-pass sequencing in dogs (27), with a mean coverage of 0.84× (range 0.17–2.05×). Reads were aligned to the CanFam3.1 reference assembly (28), and genotypes were imputed against Gencove’s reference panel of 676 dogs from 91 breeds (27). Imputed genotypes with a maximum genotype probability below 0.7 were set to missing.

Genetic sex was checked from sequencing depth on the X chromosome relative to the autosomes, and duplicate samples were removed. Within each breed, first-degree relatives were identified from identity-by-state distances, and one dog of each related pair was removed, chosen by balancing sequencing coverage, age at death and sex so as to keep ten old and ten young dogs per breed.

Variants were filtered in two stages with PLINK 2 (29). Across the sequenced and imputed dogs, biallelic variants were retained if their minor allele frequency was at least 2%, their missing genotype rate at most 20% and their Hardy–Weinberg exact test p-value at least 10⁻Z⁰, following Morrill et al. (30). This lenient Hardy–Weinberg threshold allows for the departures from equilibrium expected when inbred breeds are pooled. In the 540 dogs of the Cani-DNA discovery cohort, variants were then retained if their missing genotype rate was at most 2%, a combination with the genotype-probability threshold shown to retain most variants with few imputation errors in low-pass dog data (27), and if their minor allele frequency was at least 2% (30), leaving 8,890,296 variants.

Relatedness between dogs was modelled through the genetic relatedness matrix used in the association analyses, computed with GEMMA (31). Genetic relationships across the cohort were summarised by hierarchical clustering of pairwise identity-by-state distances computed with SNPRelate (32).

### Association testing

Associations were tested with univariate linear mixed models in GEMMA (31), with p-values from the likelihood-ratio test. Relatedness was modelled with centred genetic relatedness matrices computed from autosomal variants, using the leave-one-chromosome-out approach (LOCO) (33); variants on the X chromosome were tested with the matrix computed from all autosomes.

Breed-average body weight and breed-median lifespan were analysed as breed-level phenotypes, as in previous across-breed studies (13), and breed-median lifespan both unadjusted and adjusted for breed-average body weight. Age at death and old/young status were analysed with breed-average body weight as a covariate. Genomic inflation was assessed with the genomic inflation factor (λ) and quantile–quantile plots (Fig. S1).

Associations were declared genome-wide significant at p < 4 × 10⁻⁷, a standard threshold proposed for dog GWAS, derived by permutation across more than the 320 breeds of the Dog10K dataset (34). A suggestive threshold ten times less stringent (p < 4 × 10⁻⁶) was used to report loci for replication.

Associated variants were grouped into loci by linkage-disequilibrium clumping in PLINK 2, with the parameters of Dutrow et al. (35). Each variant with p < 4 × 10⁻⁶ could start a clump. Variants within 1 Mb of it, with p < 4 × 10⁻⁵ and rZ ≥ 0.5, joined that clump. Each clump formed a locus. Its lead variant was its most significant variant, and its region extended from the first to the last variant of the clump. A locus was genome-wide significant or suggestive according to the p-value of its lead variant.

When two loci of the same model lay within 1 Mb of each other, the weaker lead variant was tested again with the stronger lead variant as a covariate, following the stepwise approach of Hayward et al. (36). If the weaker lead variant remained below its threshold, the two loci were kept separate. Otherwise, they were merged into one locus, represented by the stronger lead variant. A breed-weight locus and a breed-lifespan locus were considered shared when their regions overlapped, and the direction of their effects was compared at the lead variant of the breed-weight locus, for the same allele in both GWAS.

### Annotation of loci

The resulting loci were then compared with those reported in previous GWAS of body size and lifespan in dogs, using CanFam3.1 coordinates. A locus was considered known if its region overlapped a published locus and novel otherwise.

Genes in each locus region were identified from the NCBI RefSeq annotation release 105 of CanFam3.1 (37), together with human RefSeq genes aligned to CanFam3.1 in the UCSC Genome Browser (38). The predicted consequence of each lead variant was obtained with the Ensembl Variant Effect Predictor (39). Each locus was annotated with the nearest named protein-coding gene to its lead variant within 1 Mb, or with its chromosomal position otherwise.

### Replication in the GRLS cohort

Loci associated with age at death were tested for replication in the Golden Retriever Lifetime Study, a prospective cohort of Golden Retrievers enrolled in the United States between 2012 and 2015, at six months to two years of age, and followed with annual veterinary examinations (40). The 3,024 dogs with survival, sex and body weight data were included. They had been genotyped with the Axiom Canine Genotyping Array Sets A and B (Thermo Fisher Scientific) (41), on CanFam3.1 coordinates. Dogs alive at the end of follow-up were censored at their last recorded visit, and body weight was averaged across visits.

The lead variants were absent from the array, so each locus was tested through a proxy variant. The proxy was the array variant within 1 Mb with the highest linkage disequilibrium with the lead variant in the 540 discovery dogs, with rZ of at least 0.7. A locus was tested when its proxy had a minor allele frequency of at least 5% in the GRLS replication cohort. The effect allele was the proxy allele in linkage disequilibrium with the discovery allele associated with longer life.

Survival was analysed with a mixed Cox proportional-hazards model in coxme (42), with the relatedness matrix of the GRLS dogs as a random effect, and sex and mean body weight as covariates. The matrix was computed with GEMMA from autosomal array variants, as in discovery. Effects were expressed as hazard ratios per copy of the effect allele. A locus was replicated when its hazard ratio was below 1 and its p-value below 0.05/K, K being the number of loci tested for the phenotype. Survival by genotype was described with Kaplan–Meier curves and log-rank tests, using the survival package (43). In the discovery cohort, the association between proxy genotype and weight-corrected age at death was tested by one-way analysis of variance (ANOVA).

### Gene-set enrichment and ageing databases

For each GWAS, the dog and human genes annotated at its loci were pooled and tested for over-representation with GSEApy (44), against seven Enrichr libraries (45): GO Biological Process 2026, KEGG 2026, Aging Perturbations from GEO, MGI Mammalian Phenotype Level 4 2024, Human Phenotype Ontology, WikiPathways 2024 Human and Reactome Pathways 2024. Terms with a Benjamini–Hochberg adjusted p-value below 0.05 were considered significant. Genes at individual-longevity loci were cross-referenced with the GenAge and LongevityMap databases of the Human Ageing Genomic Resources (46). The regions of loci replicated in the GRLS replication cohort were transferred to their orthologous regions in the human genome (GRCh38) with liftOver (38), to compare them with human longevity studies.

### Software

Genotype data were processed with PLINK 2 (v2.0.0-a.7.8), and association tests were run in GEMMA (v0.98.5). Variant consequences were predicted with the Ensembl Variant Effect Predictor (v113.0). Enrichment analyses were run in Python (v3.11.18) with GSEApy (v1.3.1). Identity-by-state distances and survival models were computed in R (v4.5.3), with SNPRelate (v1.44.0) for the distances, coxme (v2.2-22) and survival (v3.8-12) for the survival models, and data.table (v1.18.6.1) for data handling. Figures were drawn with ggplot2 (v4.0.3) (47), ggrepel (v0.9.8) (48) and survminer (v0.5.2) (49). The dendrogram was visualised with circlize (50) and dendextend (51).

## Results

### Cohort design, weight–lifespan relationship and population structure

To capture the diversity of longevity and body size across domestic dogs, we assembled the Cani-DNA discovery cohort, comprising 540 dogs from 27 breeds that represent 14 of the 23 phylogenetic clades of the domestic dog defined by Parker et al. (14) (Table S1). Their breed-average body weights range from 3 to 66 kg and their breed-median lifespans from 5.4 to 13.7 years. Across these breeds, heavier breeds lived shorter lives (Pearson r = −0.71, p = 4.0 × 10⁻⁵), losing on average 0.094 years, about a month, of median lifespan per kilogram of body weight (Fig 1A). The giant breeds lay at the short end of this range, with the Great Dane (66 kg) and the Dogue de Bordeaux (50 kg) both reaching a median of 5.4 years, while the medium-sized Whippet and Border Collie lived longest, at 13.7 and 13.1 years. The two smallest breeds, the Yorkshire Terrier and the Poodle, had median lifespans of about 10.4 years, shorter than those of the Whippet and Border Collie but nearly twice those of the Great Dane and Dogue de Bordeaux. Fig 1B shows the age at death of each dog, with old and young dogs separated within every breed.

**Fig 1.**
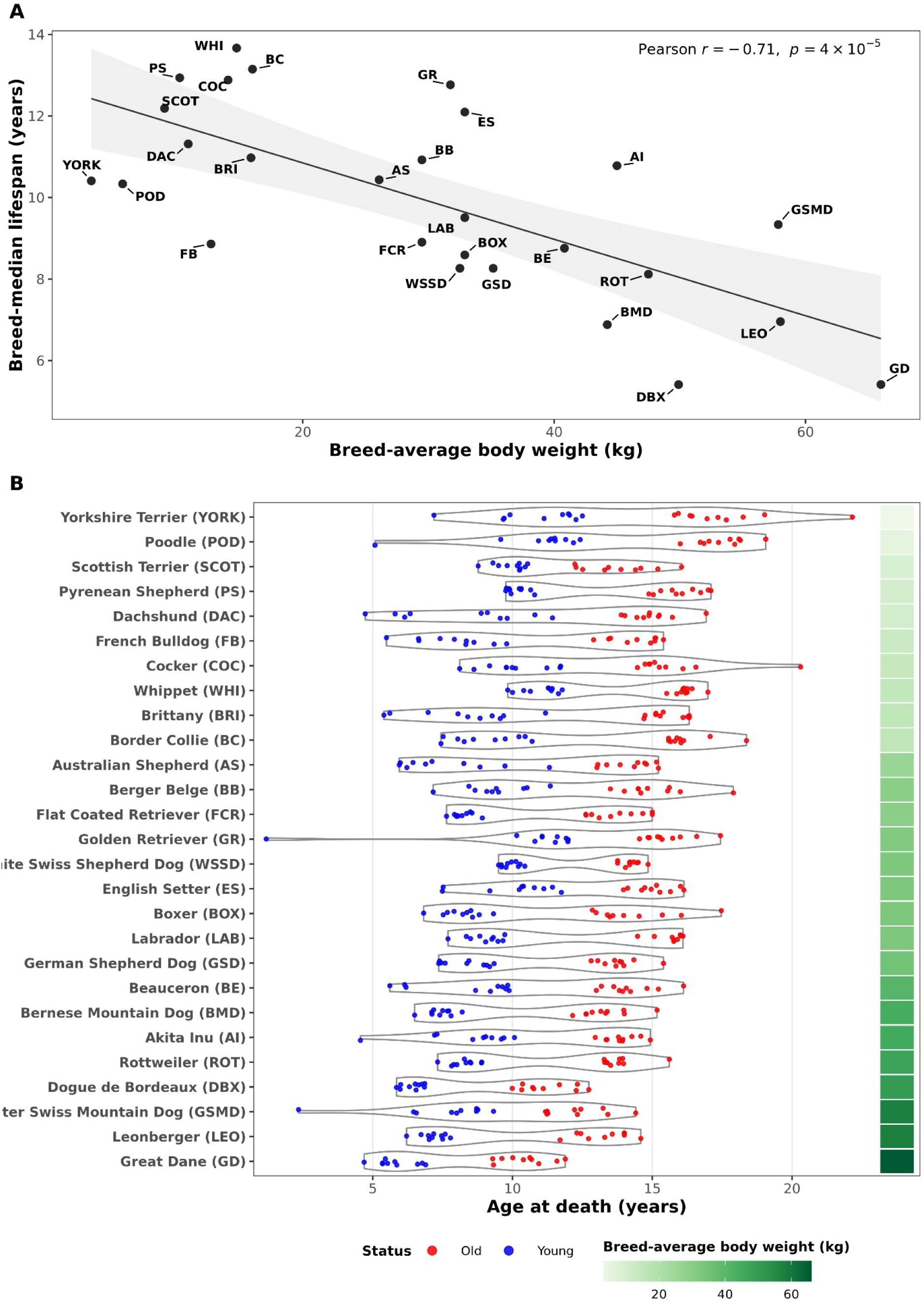
Breed-average body weight, breed-median lifespan and age at death in the Cani-DNA discovery cohort. (A) Breed-median lifespan against breed-average body weight for the 27 breeds. Each point is one breed, labelled with its code (Table S1). The line shows the linear regression, with its 95% confidence interval in grey, and the Pearson correlation coefficient (r) and its p-value are given in the panel. (B) Age at death of each of the 540 dogs, by breed. Breeds are named with their codes and ordered by breed-average body weight, from the lightest (top) to the heaviest (bottom). Each point is one dog, coloured by old/young status (red, old; blue, young), and violins show the distribution of age at death within each breed. The column on the right shows breed-average body weight on a green colour scale.

Since all association analyses rely on the genotype data, we first examined whether these data reflected the breed structure of the cohort. Unsupervised hierarchical clustering of pairwise identity-by-state distances, based on the proportion of alleles shared by each pair of dogs, recapitulated the breed structure from genotype data, with every dog clustering with its own breed (Fig 2). The genotypes of each dog therefore reflected its declared breed.

**Fig 2.**
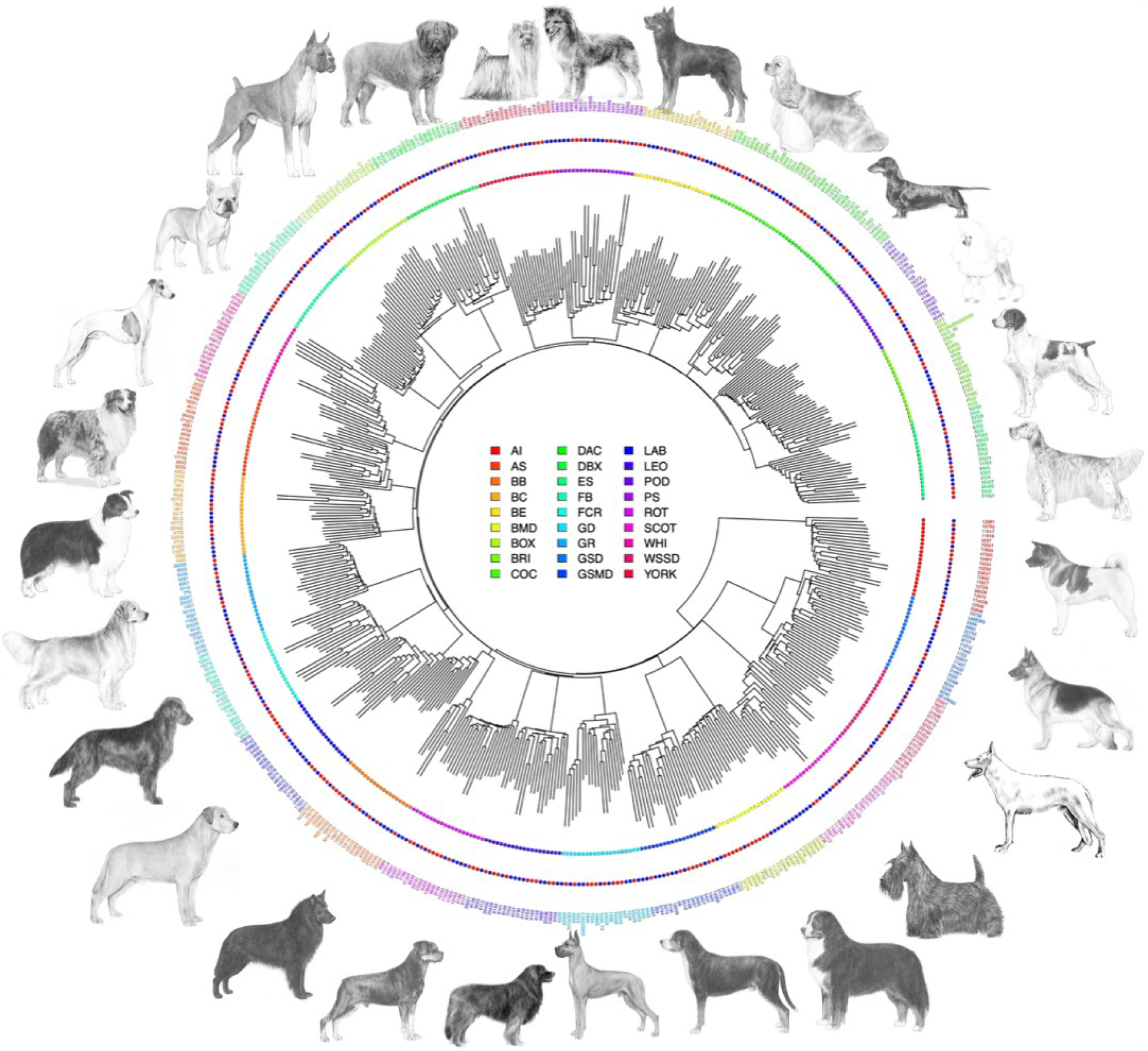
Genetic relationships between the 540 dogs of the Cani-DNA discovery cohort. Circular dendrogram from unsupervised hierarchical clustering of pairwise identity-by-state distances computed from genome-wide genotypes. Branch tips are coloured by breed, and silhouette of each breed (American Kennel Club) is shown around the periphery. Leaf marker shape indicates sex (circle, female; square, male), and the outer ring indicates old/young status (red, old; blue, young).

### Recovery of established body-size and lifespan loci

Breed body size is the best-characterised complex trait in dogs, and much of its variation across breeds is explained by a small number of loci of large effect (7,36). We therefore first performed a genome-wide association study (GWAS) of breed-average body weight, as a positive control for the ability of the cohort and the analysis pipeline to detect established loci. This GWAS identified 69 loci, 39 of them genome-wide significant (Fig 3A, top).

**Fig 3.**
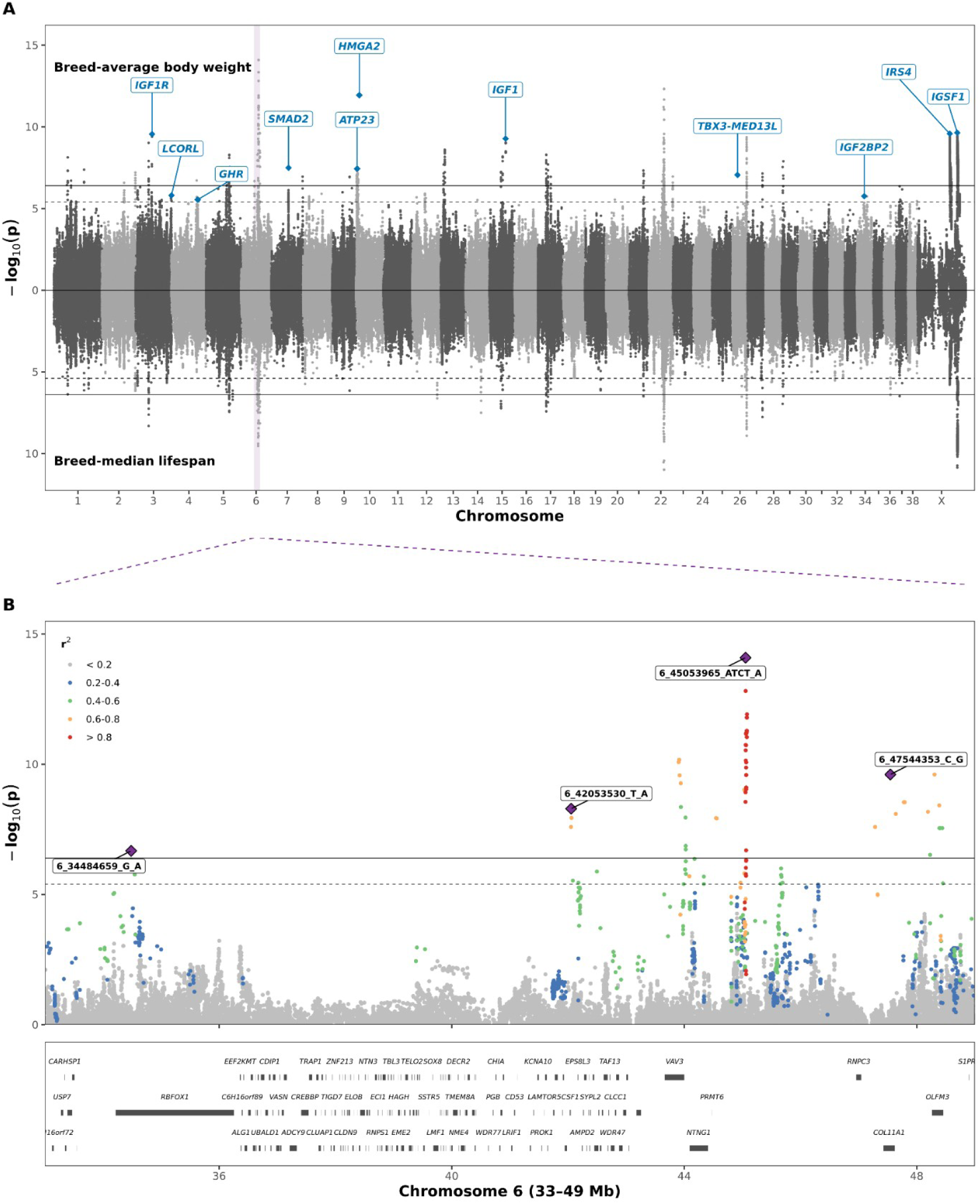
Genome-wide association with breed-average body weight and breed-median lifespan. (A) Miami plot of the GWAS of breed-average body weight (top) and breed-median lifespan (bottom) in the 540 dogs of the Cani-DNA discovery cohort. Each point is a variant, plotted by chromosomal position against −log₁₀(p) of the likelihood-ratio test. Solid lines mark the genome-wide significance threshold (p = 4 × 10⁻⁷) and dashed lines the suggestive threshold (p = 4 × 10⁻⁶). Loci overlapping previously reported loci (Table 2) are labelled with their reported genes. (B) Regional association with breed-average body weight across chromosome 6 (33–49 Mb), showing four loci, with their lead variants. Variants are coloured by their linkage disequilibrium (rZ) with the strongest lead variant, 6_45053965_ATCT_A. Protein-coding genes from the NCBI RefSeq annotation of CanFam3.1 are shown below.

**Table 2.** Previously known body-size and lifespan loci replicated in this study.

| Reported gene(s) | Lead variant | Effect allele | β (kg) | p | Breed-median lifespan | References |
|---|---|---|---|---|---|---|
| *HMGA2* | 10_8195605_T_C | T | 5.91 | 1.1 × 10<sup>−12</sup> | no | (7,36) |
| *IGSF1, ARHGAP36* | X_101730294_C_A | C | 2.69 | 2.3 × 10<sup>−10</sup> | yes | (7,36,53) |
| *IRS4, ACSL4* | X_82353310_G_A | A | 2.35 | 2.6 × 10<sup>−10</sup> | yes | (7,53) |
| *IGF1R* | 3_42142240_C_T | C | 4.45 | 2.8 × 10<sup>−10</sup> | no | (7,36) |
| *IGF1* | 15_40975163_C_T | C | 3.82 | 5.3 × 10<sup>−10</sup> | no | (4,7,36) |
| *SMAD2* | 7_43855694_T_C | T | 2.50 | 3.2 × 10<sup>−8</sup> | no | (4,7,36) |
| *ATP23* <sup>†</sup> | 10_2453907_C_T | C | 3.20 | 3.7 × 10<sup>−8</sup> | no | (52) |
| *TBX3–MED13L* | 26_13203919_G_A | G | 2.37 | 8.9 × 10<sup>−8</sup> | no | (7,36) |
| *LCORL* \* | 3_91772427_C_T | T | 3.39 | 1.6 × 10<sup>−6</sup> | no | (7,36) |
| *IGF2BP2* \* | 34_18537529_G_A | G | 2.50 | 1.7 × 10<sup>−6</sup> | no | (4,7,36) |
| *GHR* \* | 4_66936545_C_T | C | 1.75 | 2.8 × 10<sup>−6</sup> | no | (4,7,36) |

Effect allele, the allele associated with higher breed-average body weight. β, effect of each copy of the effect allele on breed-average body weight. Breed-median lifespan, locus also associated with breed-median lifespan in this study.
\* Suggestive locus (4 × 10<sup>−7</sup> ≤ p < 4 × 10<sup>−6</sup>). All other loci are genome-wide significant (p < 4 × 10<sup>−7</sup>).
† Locus reported for breed life expectancy. All other loci were reported for body size.

Eleven of these loci overlapped previously reported loci (Table 2). Ten were established body-size loci, including those near *IGF1*, *IGF1R*, *HMGA2* and *LCORL*, and the eleventh overlapped *ATP23*, a locus reported for breed life expectancy (52), a trait strongly correlated with breed weight. The remaining 58 loci were novel, 31 of them genome-wide significant (Table S2). By recovering established body-size loci, the breed-average body weight GWAS showed that the cohort and the analysis pipeline detect known associations.

In parallel, we performed a GWAS of breed-median lifespan, which identified 53 loci, 21 of them genome-wide significant (Fig 3A, bottom). Two of these loci overlapped loci previously reported for lifespan, both on the X chromosome, in the regions containing *IGSF1* (7,18) and *IRS4* (7), and both also overlapped breed-weight loci (Table 2). At the *IGSF1* locus, the breed-average body weight and breed-median lifespan GWAS shared the same lead variant, X_101730294_C_A. Its C allele was associated with higher breed-average body weight (2.69 kg per copy, p = 2.3 × 10⁻^10^) and shorter breed-median lifespan (0.36 years per copy, p = 1.4 × 10⁻^11^), consistent with the inverse relationship between breed-average body weight and breed-median lifespan across breeds (Fig 1A).

### Loci shared between breed-average body weight and breed-median lifespan follow the size–lifespan trade-off

Across dog breeds, heavier breeds have shorter lifespans than lighter ones (1,3), so a locus influencing breed size may also appear as a lifespan locus. We therefore examined which breed-weight loci were shared with breed-median lifespan. Of the 69 breed-weight loci, 30 overlapped a breed-lifespan locus, comprising the two known X-chromosome loci described above and 28 novel loci (Table S3, Fig S2). At all 30 loci, the allele associated with higher breed-average body weight was associated with shorter breed-median lifespan (Fig S2).

The strongest association of the breed-weight GWAS lay on chromosome 6 (p = 8.1 × 10⁻^15^), in a region containing *VAV3*, *NTNG1* and *PRMT6*. In humans, a locus near *PRMT6* has been associated with whole-body lean mass in women (54), and in pigs, fine-mapping of birth weight pointed to *VAV3* and *NTNG1* (55). This locus was shared with breed-median lifespan, as were two other breed-weight loci on the same chromosome (Fig 3B). One lay near *CSF1*, and the other within *COL11A1*, a gene associated with bone size in humans (56) and required for normal limb-bone growth in mice (57). A fourth locus on chromosome 6, within *RBFOX1*, was specific to breed-average body weight.

### Lifespan loci independent of breed-average body weight

Beyond the loci shared with body weight, 21 of the 53 breed-lifespan loci overlapped no breed-weight locus (Table S4, Fig S2). All were novel, and 5 reached genome-wide significance. The strongest (p = 3.2 × 10⁻⁸) lay on chromosome 14, in a region containing *NFE2L3*, *HNRNPA2B1* and *CBX3*. Expression of *HNRNPA2B1* has been associated with lifespan across mouse strains and with parental longevity in humans (58).

Because breed-average body weight and breed-median lifespan are correlated across dog breeds (1,3), any breed-lifespan locus may partly reflect this correlation, including loci that overlap no breed-weight locus. We therefore repeated the breed-median lifespan GWAS with breed-average body weight as a covariate. This analysis identified 29 loci, 4 of them genome-wide significant (Table S5, Fig S4). All were novel, and only one of them overlapped a breed-weight locus.

Five of the 29 loci overlapped one of the 21 lifespan-specific loci identified without adjustment. They included the loci within *PIP5K1B* and in the region containing *HNRNPA2B1*, both genome-wide significant after adjustment. They also included a region containing *PIK3IP1*, which encodes a negative regulator of PI3K signalling (59), a pathway whose partial inactivation extends lifespan in male mice (60). Twenty-two loci, among them the genome-wide significant loci within *CACNB4* and *CRTAC1*, were detected only after adjustment. Adjusting for breed-average body weight thus retained the lifespan-specific loci and revealed further loci associated with breed-median lifespan.

### Loci associated with individual longevity

The breed-level analyses compare breeds, so they cannot explain why one dog outlives another dog of the same breed. We therefore turned to individual longevity, analysed in two ways, with breed-average body weight as a covariate. Old/young status compared the 270 old dogs, among the oldest of their breed, with the 270 young dogs, which died around their breed-median lifespan, and identified no locus (Fig S5). Age at death used the exact age at which each dog died and identified 11 loci, 2 of them genome-wide significant (Fig 4).

**Fig 4.**
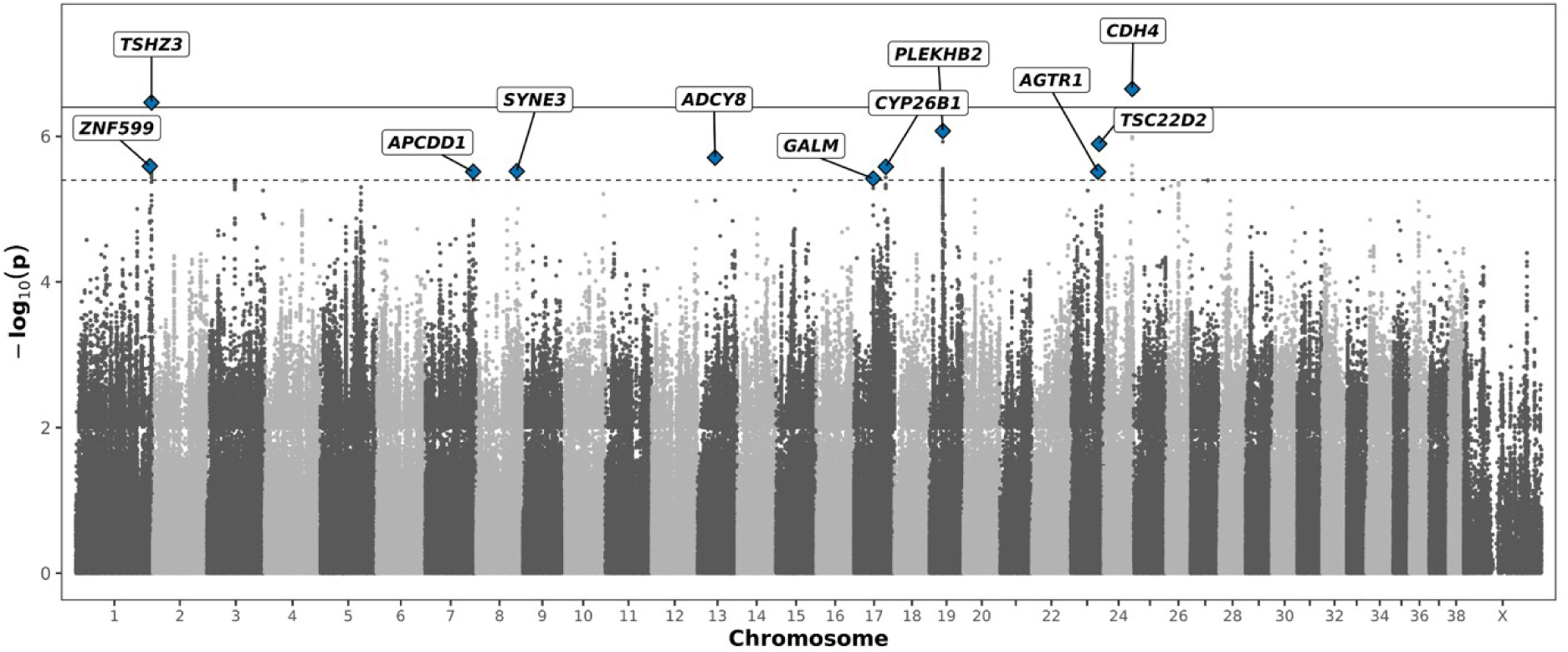
GWAS of age at death in the Cani-DNA discovery cohort. Manhattan plot of age at death, with breed-average body weight as a covariate, in the 540 dogs of the Cani-DNA discovery cohort. Each point is one variant, plotted by chromosomal position against its −log₁₀(p). Solid and dashed horizontal lines mark the genome-wide significance (p = 4 × 10⁻^7^) and suggestive (p = 4 × 10⁻^6^) thresholds. Blue diamonds mark the lead variants of the 11 loci, each labelled with its nearest gene.

At these 11 loci, each copy of the effect allele, defined as the allele associated with longer life, increased age at death by 0.97 to 2.98 years and none of the loci overlapped a breed-level locus.

Several of these loci lay near genes previously linked to longevity or ageing. The two genome-wide significant loci were the *CDH4* locus (p = 2.2 × 10⁻⁷) and the *TSHZ3* locus (p = 3.4 × 10⁻⁷). A common variant in *CDH4* was among 27 variants associated with lifespan in the Framingham Heart Study (61), and Alu deletions in *CDH4* are components of polygenic predictors of human longevity (62). *TSHZ3* was the gene most enriched for rare protein-altering variants in whole-genome sequences of supercentenarians, although this enrichment was not significant in a second cohort of long-lived individuals (24). Two of the suggestive loci were the *TSC22D2* and *AGTR1* loci. In humans, the *TSC22D2*–*LINC01214* region carries variants associated with epigenetic age acceleration (63,64) and with mosaic loss of chromosome Y in men (65). *AGTR1* encodes the angiotensin II type 1 receptor, whose disruption prolongs lifespan in mice (66).

We then tested whether the genes at the 11 loci were over-represented in known pathways and functional categories. Genes at three loci were over-represented in the WikiPathways pathway “Envelope proteins and potential roles in Emery–Dreifuss muscular dystrophy (EDMD) physiopathology” (3 of 46 genes, adjusted p = 0.0025; Table S7). *SYNE3* and *ADCY8* lie on chromosomes 8 and 13, and *BANF1* lies within the *GALM* locus on chromosome 17 (Table S6). In humans, mutations in *BANF1* cause a progeroid syndrome, a disorder with features of premature ageing (67).

### Replication of age-at-death loci in Golden Retrievers

The Cani-DNA discovery cohort relied on imputed genotypes from low-coverage sequencing and combined 27 breeds. We therefore tested the age-at-death loci in the GRLS replication cohort, from the Golden Retriever Lifetime Study, an independent prospective cohort of 3,024 Golden Retrievers followed from early life and genotyped on an array (68). Lead variants were absent from the GRLS array, so each locus was tested through a proxy, an array variant in strong linkage disequilibrium with the lead variant of the locus (rZ ≥ 0.7 in the 540 dogs of the Cani-DNA discovery cohort). Such a proxy existed for four loci (Table 3). For the other seven loci, no variant on the GRLS array met this criterion, so these loci could not be tested in the GRLS replication cohort.

**Table 3.**
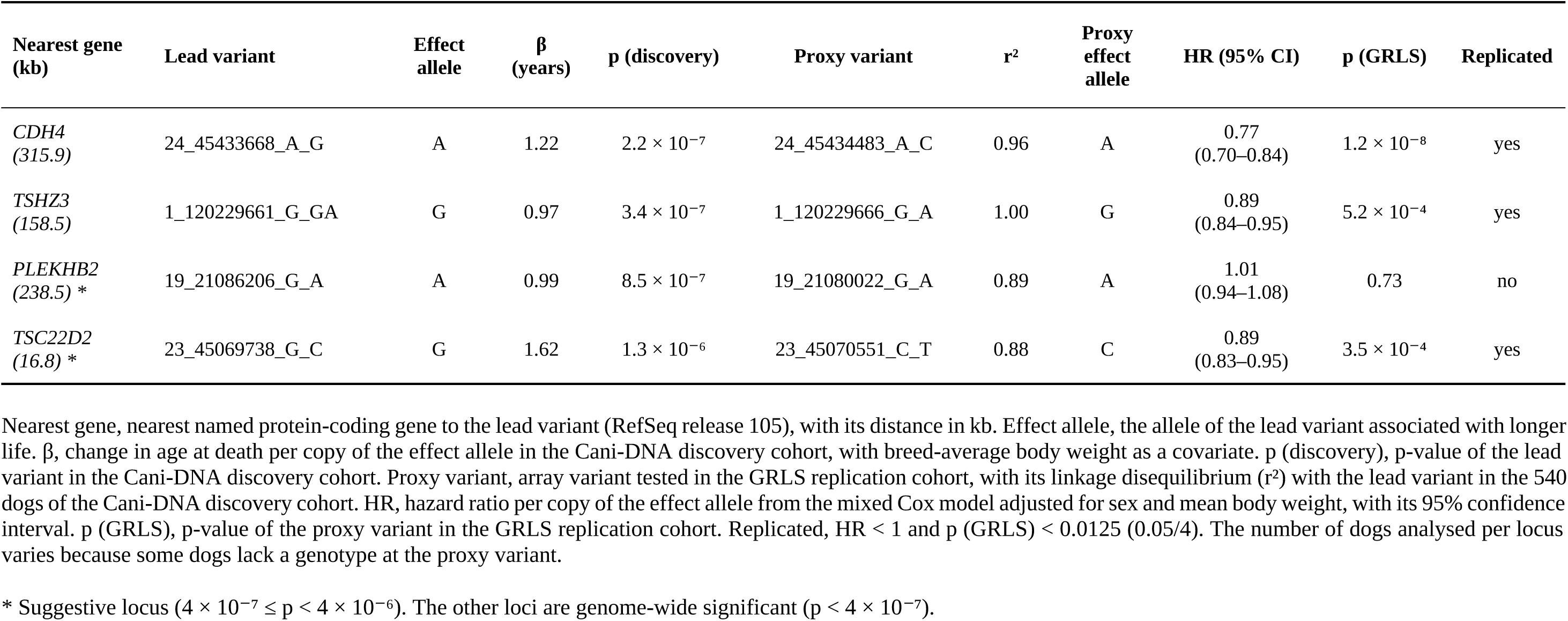
Replication of age-at-death loci in the GRLS replication cohort.

| **Nearest gene (kb)** | **Lead variant** | **Effect allele** | **β (years)** | **p (discovery)** | **Proxy variant** | **r<sup>2</sup>** | **Proxy effect allele** | **HR (95% CI)** | **p (GRLS)** | **Replicated** |
|---|---|---|---|---|---|---|---|---|---|---|
| *CDH4*<br>*(315.9)* | 24_45433668_A_G | A | 1.22 | 2.2 × 10<sup>−7</sup> | 24_45434483_A_C | 0.96 | A | 0.77<br>(0.70–0.84) | 1.2 × 10<sup>−8</sup> | yes |
| *TSHZ3*<br>*(158.5)* | 1_120229661_G_GA | G | 0.97 | 3.4 × 10<sup>−7</sup> | 1_120229666_G_A | 1.00 | G | 0.89<br>(0.84–0.95) | 5.2 × 10<sup>−4</sup> | yes |
| *PLEKHB2*<br>*(238.5)* \* | 19_21086206_G_A | A | 0.99 | 8.5 × 10<sup>−7</sup> | 19_21080022_G_A | 0.89 | A | 1.01<br>(0.94–1.08) | 0.73 | no |
| *TSC22D2*<br>*(16.8)* \* | 23_45069738_G_C | G | 1.62 | 1.3 × 10<sup>−6</sup> | 23_45070551_C_T | 0.88 | C | 0.89<br>(0.83–0.95) | 3.5 × 10<sup>−4</sup> | yes |

Nearest gene, nearest named protein-coding gene to the lead variant (RefSeq release 105), with its distance in kb. Effect allele, the allele of the lead variant associated with longer life. β, change in age at death per copy of the effect allele in the Cani-DNA discovery cohort, with breed-average body weight as a covariate. p (discovery), p-value of the lead variant in the Cani-DNA discovery cohort. Proxy variant, array variant tested in the GRLS replication cohort, with its linkage disequilibrium (r<sup>2</sup>) with the lead variant in the 540 dogs of the Cani-DNA discovery cohort. HR, hazard ratio per copy of the effect allele from the mixed Cox model adjusted for sex and mean body weight, with its 95% confidence interval. p (GRLS), p-value of the proxy variant in the GRLS replication cohort. Replicated, HR < 1 and p (GRLS) < 0.0125 (0.05/4). The number of dogs analysed per locus varies because some dogs lack a genotype at the proxy variant.
\* Suggestive locus (4 × 10<sup>−7</sup> ≤ p < 4 × 10<sup>−6</sup>). The other loci are genome-wide significant (p < 4 × 10<sup>−7</sup>).

At each proxy, the effect allele was the allele associated with longer life in the Cani-DNA discovery cohort. In this cohort, weight-corrected age at death, the age at which a dog died relative to the lifespan expected from its breed-average body weight, increased with each copy of the effect allele at all four proxies (ANOVA p = 3.5 × 10⁻⁷ to 2.3 × 10⁻⁴; Fig 5A, Fig S6A).

**Fig 5.**
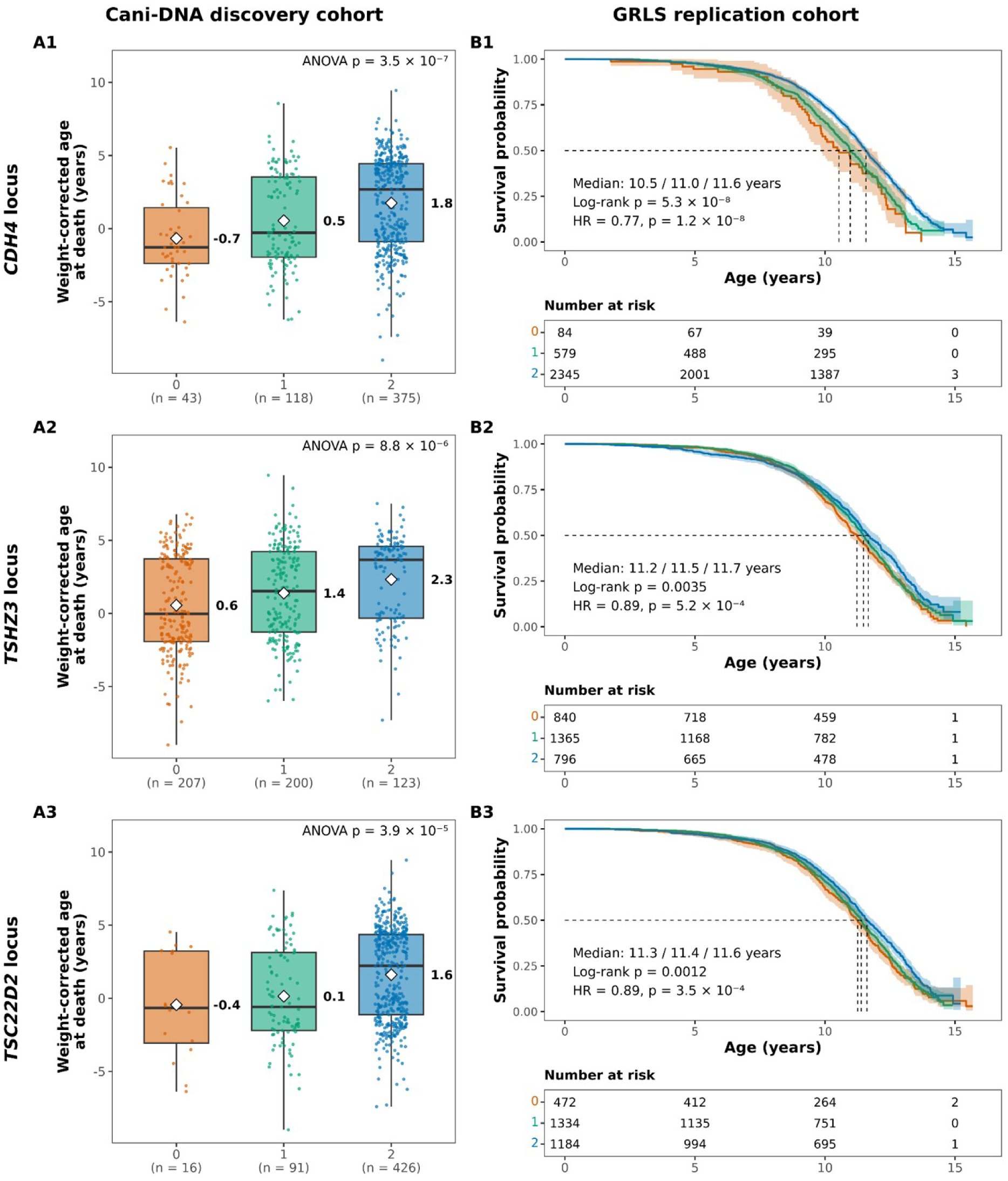
Replication of three age-at-death loci in the GRLS replication cohort. Rows show the *CDH4* (1), *TSHZ3* (2) and *TSC22D2* (3) loci, each tested through its proxy variant (Table 3). Genotypes are given as the number of copies of the effect allele (0, orange; 1, green; 2, blue). (A) Weight-corrected age at death in the Cani-DNA discovery cohort by genotype at the proxy variant. Boxes show the median, white diamonds and the labels beside them give the genotype means, and the number of dogs is shown below each genotype. The p-value is from a one-way analysis of variance. (B) Kaplan–Meier survival curves in the GRLS replication cohort by genotype at the same proxy variant, with 95% confidence intervals. Dashed lines mark median survival. The hazard ratio (HR) per copy of the effect allele, with its 95% confidence interval and p-value, is from the mixed Cox model adjusted for sex and mean body weight. The log-rank p-value compares the three survival curves.

In the GRLS replication cohort, dogs carrying two copies of the effect allele lived longer than dogs carrying zero copies at three of the four loci. At the *CDH4* locus, dogs carrying two copies of the effect allele had a median survival of 11.6 years, against 10.5 years for dogs carrying zero copies (log-rank p = 5.3 × 10⁻⁸; Fig 5B). At the *TSHZ3* locus, dogs carrying two copies of the effect allele had a median survival of 11.7 years, against 11.2 years for dogs carrying zero copies (log-rank p = 0.0035). At the *TSC22D2* locus, dogs carrying two copies of the effect allele had a median survival of 11.6 years, against 11.3 years for dogs carrying zero copies (log-rank p = 0.0012). Survival did not differ by genotype at the *PLEKHB2* locus (log-rank p = 0.93; Fig S6B).

Because survival curves do not account for relatedness between dogs or for differences in sex and body weight, we tested each locus with a mixed Cox model including them. Three loci replicated in the GRLS replication cohort, with a hazard ratio below 1 and p < 0.0125 (Table 3). At the *CDH4* locus, each copy of the effect allele lowered the risk of death at any given age by 23% (HR = 0.77, 95% CI 0.70–0.84, p = 1.2 × 10⁻⁸), so dogs carrying two copies had about 41% lower risk than dogs carrying zero copies. At the *TSHZ3* locus, each copy of the effect allele lowered the risk of death at any given age by 11% (HR = 0.89, 95% CI 0.84–0.95, p = 5.2 × 10⁻⁴), so dogs carrying two copies had about 21% lower risk than dogs carrying zero copies. At the *TSC22D2* locus, each copy of the effect allele also lowered the risk of death at any given age by 11% (HR = 0.89, 95% CI 0.83–0.95, p = 3.5 × 10⁻⁴), so dogs carrying two copies had about 21% lower risk than dogs carrying zero copies. The *PLEKHB2* locus did not replicate (HR = 1.01, 95% CI 0.94–1.08, p = 0.73). Replication in an independent single-breed cohort thus supports the *CDH4*, *TSHZ3* and *TSC22D2* loci as associated with individual longevity, independently of body weight.

## Discussion

This study aimed to identify loci associated with longevity in dogs beyond the effect of breed body size. Genome-wide studies of dog longevity have so far used breed-level phenotypes (4,7,16,18,52) or compared long- and short-lived dogs within a single breed (19,20). Neither design captures the variation in longevity between individual dogs across breeds. We analysed this variation directly, as the age at death of individual dogs from 27 breeds with breed-average body weight as a covariate, and identified 11 loci. To our knowledge, this is the first multi-breed GWAS of individual age at death in dogs. Three of the four loci that could be tested replicated in an independent cohort of Golden Retrievers.

The *CDH4*, *TSHZ3* and *TSC22D2* loci, which replicated in the GRLS replication cohort, harbour genes with links to human ageing. *CDH4* encodes R-cadherin, a member of the cadherin family of cell adhesion molecules. Its promoter is frequently methylated and silenced in colorectal and gastric cancers, which suggests a role as a tumour suppressor (69). Variation in *CDH4* has also been associated with human longevity. In the Framingham Heart Study, the frequency of the minor allele of an intronic variant in *CDH4* increased with age, and this variant was among 27 selected for an effect on lifespan by six statistical methods (61). In ethnic Tatars from the Volga-Ural region of Russia, genotype combinations including the deletion allele of an Alu polymorphism in *CDH4* were more frequent among people aged 90 years or older (62). Yashin and colleagues proposed that cell adhesion genes such as *CDH4* could favour longevity through protection against cancer (61). This hypothesis could be tested in dogs, in which cancer is a leading cause of death in older dogs (70).

*TSHZ3* encodes a zinc-finger transcription factor required for the development and function of cortical projection neurons, and its heterozygous deletion causes a syndrome that includes autism spectrum disorder (71). Its link with longevity comes from the whole-genome sequencing of 17 supercentenarians, in which *TSHZ3* was the gene most enriched for rare protein-altering variants (24). This enrichment did not reach significance in 99 further long-lived individuals. The association of the *TSHZ3* locus with age at death in the Cani-DNA discovery cohort, and its replication in Golden Retrievers, point again to this gene as a candidate for longevity.

*TSC22D2* encodes a member of the TSC22 domain family of transcriptional regulators. It was first described as a cancer-associated gene in a family affected by multiple cancers, and its overexpression inhibits the growth of colorectal cancer cells (72). In humans, the region orthologous to the *TSC22D2* locus lies about 105 kb from a variant associated with epigenetic age acceleration (73) and about 91 kb from a variant associated with mosaic loss of chromosome Y in men (65). Both traits are markers of biological ageing that predict mortality (74,75). In each of the three cases, evidence from human studies of ageing finds independent support in dogs, which therefore offer a setting in which such leads can be tested (11).

Replication in the GRLS replication cohort is informative because this cohort avoids the two main weaknesses of the Cani-DNA discovery cohort. Its dogs were genotyped on an array, which guards against artefacts of the low-coverage imputation used in discovery. They also belong to a single breed. In a cohort of many breeds, an allele can appear associated with lifespan simply because it is more common in breeds that live longer for other reasons. Within a single breed, this explanation does not apply, so the replicated loci are associated with how long dogs live within a breed, and not only with differences between breeds. Confounding within the breed remains possible, which is why the mixed Cox model accounted for relatedness, sex and body weight. Body weight was also measured differently in the two cohorts, as a breed average in the Cani-DNA discovery cohort and for each dog in the GRLS replication cohort, and three loci replicated despite this difference.

The single-breed design also limits what could be tested. Only alleles that segregate in Golden Retrievers can be tested in this cohort, and linkage disequilibrium between a lead variant and its proxy, estimated across 27 breeds, may differ within one. All four tested effect alleles segregated in the GRLS replication cohort, at frequencies of 0.41 to 0.88, but seven of the 11 loci had no suitable proxy on the array and remain untested. At the *PLEKHB2* locus, the effect allele was common in Golden Retrievers (0.41), so the lack of replication does not reflect its absence. It may reflect an effect that depends on genetic background, or a false positive in the discovery cohort. Genetic background can modify, and even reverse, the effect of a variant or of an intervention. This has been shown for null alleles studied on 30 mouse strains (76) and for dietary restriction, which extends lifespan in some recombinant inbred mouse strains and shortens it in others (77).

Among the seven loci that could not be tested in the GRLS replication cohort, the *AGTR1* locus is of particular interest. *AGTR1* encodes the angiotensin II type 1 receptor, which mediates most of the effects of angiotensin II in the renin–angiotensin system (78), and disruption of this receptor in mice prolongs lifespan (79). The locus reached only suggestive significance, and the lack of a proxy on the GRLS array left it untested, so its replication is a priority for future cohorts. Beyond individual loci, the genes at three age-at-death loci, *SYNE3*, *ADCY8* and *BANF1*, were over-represented in a pathway of nuclear envelope proteins implicated in Emery–Dreifuss muscular dystrophy. Alterations of the nuclear envelope have been linked to human ageing (80), and mutations in *BANF1*, one of these three genes, cause a hereditary progeroid syndrome (67). *BANF1* was, however, assigned to the *GALM* locus through the human gene annotation and is absent from the dog annotation of this region. Testing the *AGTR1* locus and the loci of the nuclear-envelope pathway will require cohorts genotyped at their lead variants.

The breed-level analyses detect variants whose frequency differs between breeds. That such level loci could be detected in 540 dogs reflects the structure of dog breeds. Selective breeding has made breeds genetically distinct populations (14), between which variants of large effect on morphology differ markedly in frequency (13). This advantage applies mainly to such large-effect variants, and it allowed the GWAS of breed-average body weight to recover established body-size loci, such as those near *IGF1*, *IGF1R*, *HMGA2* and *LCORL* (7,36).

At all 30 loci shared between breed-average body weight and breed-median lifespan, the allele increasing weight was associated with shorter lifespan, as expected from the inverse relationship between size and lifespan across breeds (1,3). These shared loci cannot, however, separate a direct effect of a locus on both traits from the correlation between the two breed phenotypes, since any locus that changes breed size will also be associated with breed lifespan. Some genes at these loci may nonetheless act on both traits. One candidate is *PRKG1*, which lies at a shared locus and whose orthologue in *Caenorhabditis elegans*, *egl-4*, regulates both body size and lifespan through two separate pathways (81).

Adjusting breed-median lifespan for breed-average body weight addressed this limitation. After adjustment, the *IRS4* region was no longer associated with lifespan, and the association of the *IGSF1* region fell below the suggestive threshold (p = 2.5 × 10⁻^4^). Their association with breed lifespan therefore appears to act largely through body size. The association remaining at the *IGSF1* region agrees with a previous study that reported this region for lifespan adjusted for sex and body size (18). Five of the 21 lifespan-specific loci remained associated after adjustment, which makes them candidates for effects on lifespan independent of size. Among them, the locus whose region contains *HNRNPA2B1* remained genome-wide significant. *HNRNPA2B1* encodes an RNA-binding protein that regulates RNA splicing, stability and transport (82), and its expression has been associated with lifespan across mouse strains and with parental longevity in humans (58). The locus near *PRDM1* also remained associated. *PRDM1* encodes a transcriptional regulator of lymphocyte differentiation whose expression increases with age in human lymphocytes, which suggests a role in immunosenescence (83). The locus containing *PIK3IP1* points to PI3K signalling, since *PIK3IP1* negatively regulates PI3K (59), and partial inactivation of PI3K extends lifespan in male mice (60).

All breed-level loci share a limitation. Breeds differ in many traits besides size, so a locus selected for one trait can become associated with another. Two weight-specific loci on chromosome 13, which flank *RSPO2* and contain *OXR1* and *PKHD1L1*, illustrate this limitation. An insertion in the 3′ untranslated region of *RSPO2* causes furnishings, the moustache and eyebrows of wire-haired breeds (84), and long hair in Sapsaree dogs (85). The same study associated the surrounding region, including *OXR1* and *PKHD1L1*, with hair length, and found a signature of selection on this region in long-haired breeds (85). *PKHD1L1* was also associated with hair length in a second study (86). These two loci may therefore reflect selection on coat or an effect on body weight itself. Breeds differ in both coat and size, and breeders may have selected the two together, so alleles selected for coat could become associated with breed weight. Variants selected for coat can also affect other traits, as shown in dogs for variants at *RSPO2* and *FGF5*, which determine fur type and are also associated with blood metabolite levels (87). Alternatively, these loci may act on body weight directly. The *PKHD1L1* locus contains *TRHR*, which encodes the thyrotropin-releasing hormone receptor and has been associated with lean body mass in humans (88). In dogs, a GWAS of features extracted from whole-body images by deep learning also identified this chromosome 13 region, and proposed *TRHR* as a candidate gene for both body development and hair growth (89). Our data therefore cannot tell whether these two loci affect body weight directly or reflect selection on coat. Our data therefore cannot tell whether these two loci affect body weight directly or reflect selection on coat.

At the individual level, old/young status identified no locus, whereas age at death identified 11. Age at death, with breed-average body weight as a covariate, captures variation between dogs, and its 11 loci were distinct from all loci found at the breed level, which suggests that the two levels of analysis capture different parts of the genetic variation in longevity. With 270 old and 270 young dogs, the comparison was well powered for large effects, whereas complex traits such as longevity are expected to involve many variants of small effects (61). In dogs, large effects come mainly from variants that differ between breeds (13,16). Because old and young dogs were selected within each breed, both groups carried these variants at similar frequencies, so they could not contribute to the comparison. The determinants of extreme longevity may also differ between breeds, since causes of death themselves differ by breed (70). A variant favouring old age would then be carried by the old dogs of only a few breeds, and its signal would be diluted across the 27 breeds. The definition of old dogs added further heterogeneity. Old dogs were defined relative to their own breed’s distribution of age at death, and their ages ranged from below the 80th to at or above the 99th percentile across breeds, which weakened the contrast between groups. Age at death, analysed as a continuous trait, does not depend on this definition, which may explain why it detected loci where old/young status did not.

These limitations also show how this work can be extended. The loci identified here are candidate regions, and the causal variants at the replicated loci remain to be identified by fine-mapping and functional studies. Long-read sequencing would help this effort, since it resolves structural variants and transposable element insertions that short-read data miss (90). Larger cohorts recording the age at death and body weight of each dog would allow both traits to be analysed at the individual level, and genotyping at the lead variants would allow the seven untested loci to be tested for replication.

In summary, analysing the age at death of individual dogs across 27 breeds identified loci associated with longevity independently of breed body size, three of which replicated in an independent cohort of Golden Retrievers. Several of these loci harbour genes previously implicated in human longevity and ageing, which supports the dog as a model in which genetic leads on ageing can be identified and tested. With further validation, these findings could inform breeding strategies that promote longer and healthier lives in dogs.

## Acknowledgements

We thank the Cani-DNA Biological Resource Centre (UAR BIOSIT, Université de Rennes) for the canine samples, obtained from its main site in Rennes and from its four other sites at the French national veterinary schools of Nantes (Oniris), Alfort (EnvA), Toulouse (ENVT) and Lyon (VetAgro Sup). We thank the veterinarians of the Canine Genetics team (IGDR), in particular A.-S. Guillory, A. Garand, F. Allais, A. Guyon and A. Prouteau, and the Société Centrale Canine (SCC), which helped to collect samples from old dogs through targeted communication. We thank I-CAD, the French national register of identified dogs, for updating the dates of death of the dogs in our cohort. We also thank the French breed clubs, veterinarians, breeders and owners who volunteered to donate samples, health data and follow-up information for their dogs. We are grateful to our colleagues at IGDR for their advice and help, in particular Mathias Hebras for helpful discussions on the GWAS analyses, and to the GenOuest bioinformatics core facility (https://www.genouest.org) for data storage and computing infrastructure.

We would like to thank Morris Animal Foundation, its staff members and all participants in the Golden Retriever Lifetime Study, including the dog owners, their golden retrievers and the Study veterinarians who made this work possible.

## Disclaimer

The views expressed in this publication are those of the author(s) and do not necessarily reflect the views or policies of the Food and Agriculture Organization of the United Nations.

## Supplementary figures

**Fig S1.**
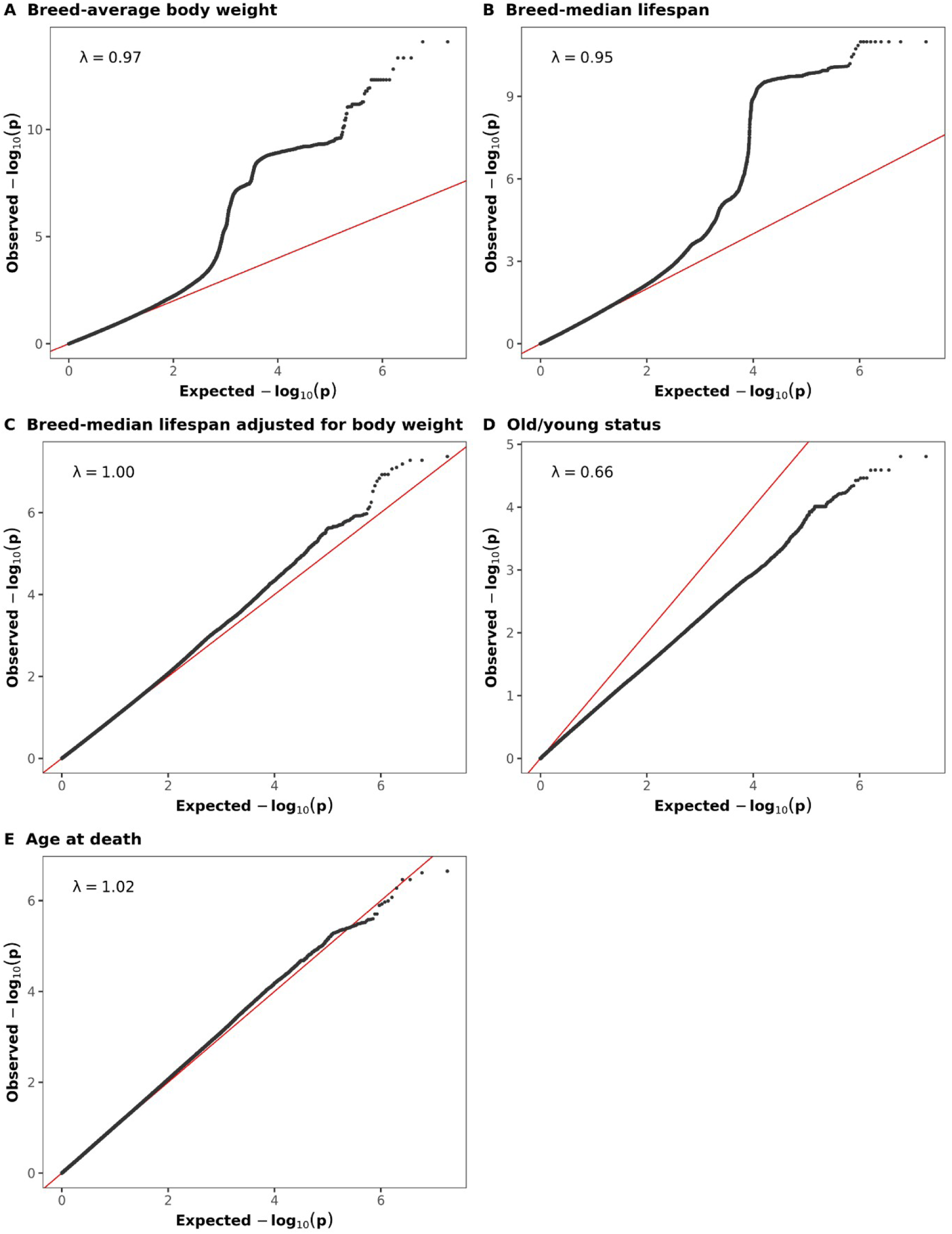
Quantile–quantile plots of the five genome-wide association analyses. Observed against expected −log₁₀(p) for the likelihood-ratio test at 8,890,296 variants, for breed-average body weight (A), breed-median lifespan (B), breed-median lifespan adjusted for breed-average body weight (C), old/young status (D) and age at death (E). The red line marks the expected distribution under the null hypothesis, and λ is the genomic inflation factor.

**Fig S2.**
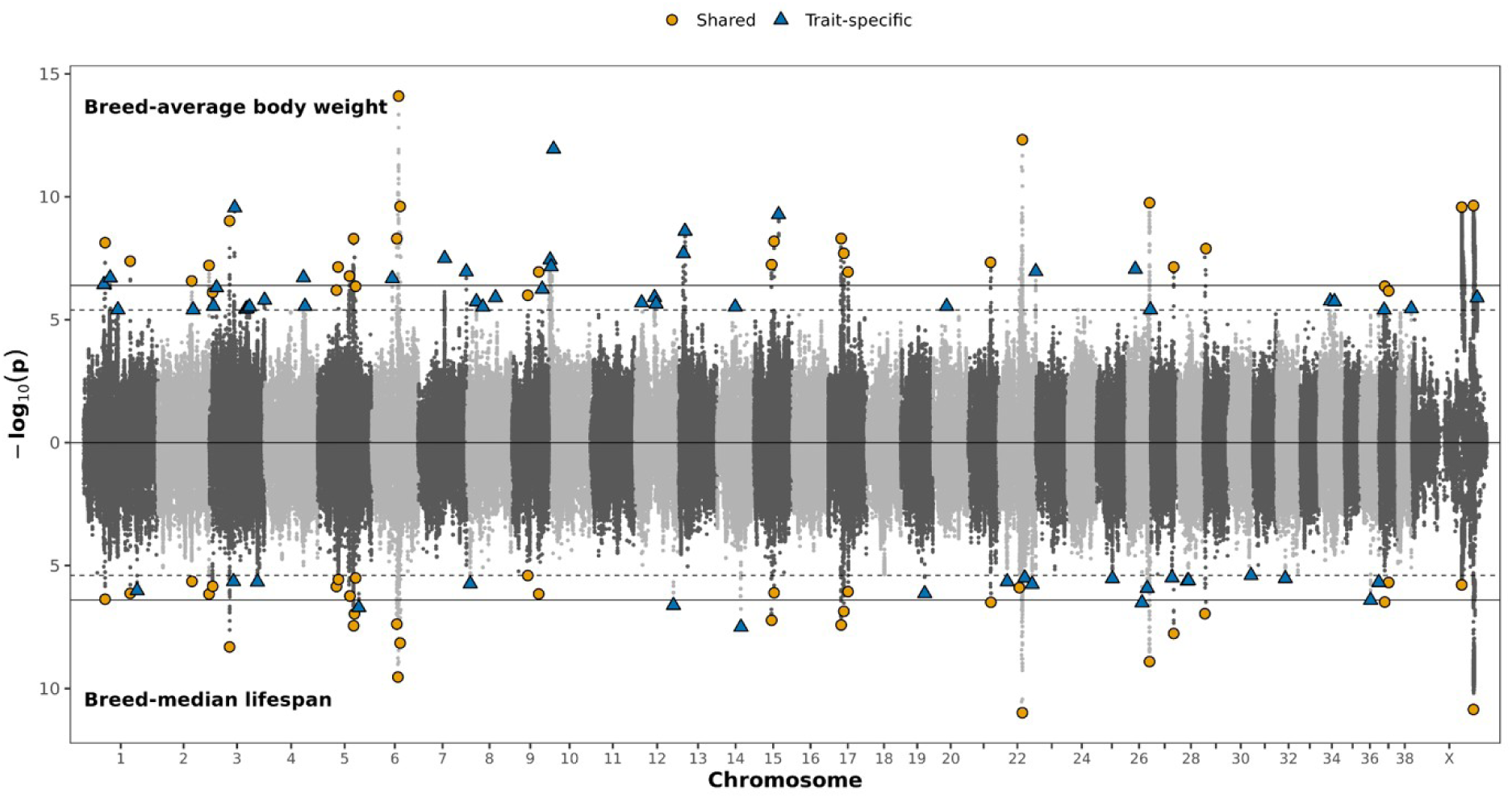
Loci shared between breed-average body weight and breed-median lifespan. Miami plot of the GWAS of breed-average body weight (top) and breed-median lifespan (bottom) in the 540 dogs of the Cani-DNA discovery cohort. Each point is one variant, plotted by chromosomal position against its −log₁₀(p). Solid and dashed horizontal lines mark the genome-wide significance (p = 4 × 10⁻⁷) and suggestive (p = 4 × 10⁻⁶) thresholds. The lead variant of each locus is marked by category. Orange triangles mark loci shared between the two GWAS: 30 breed-weight loci (top) and the 32 breed-lifespan loci overlapping them (bottom). Blue circles mark loci specific to one trait: 39 breed-weight loci (top) and 21 breed-lifespan loci (bottom).

**Fig S3.**
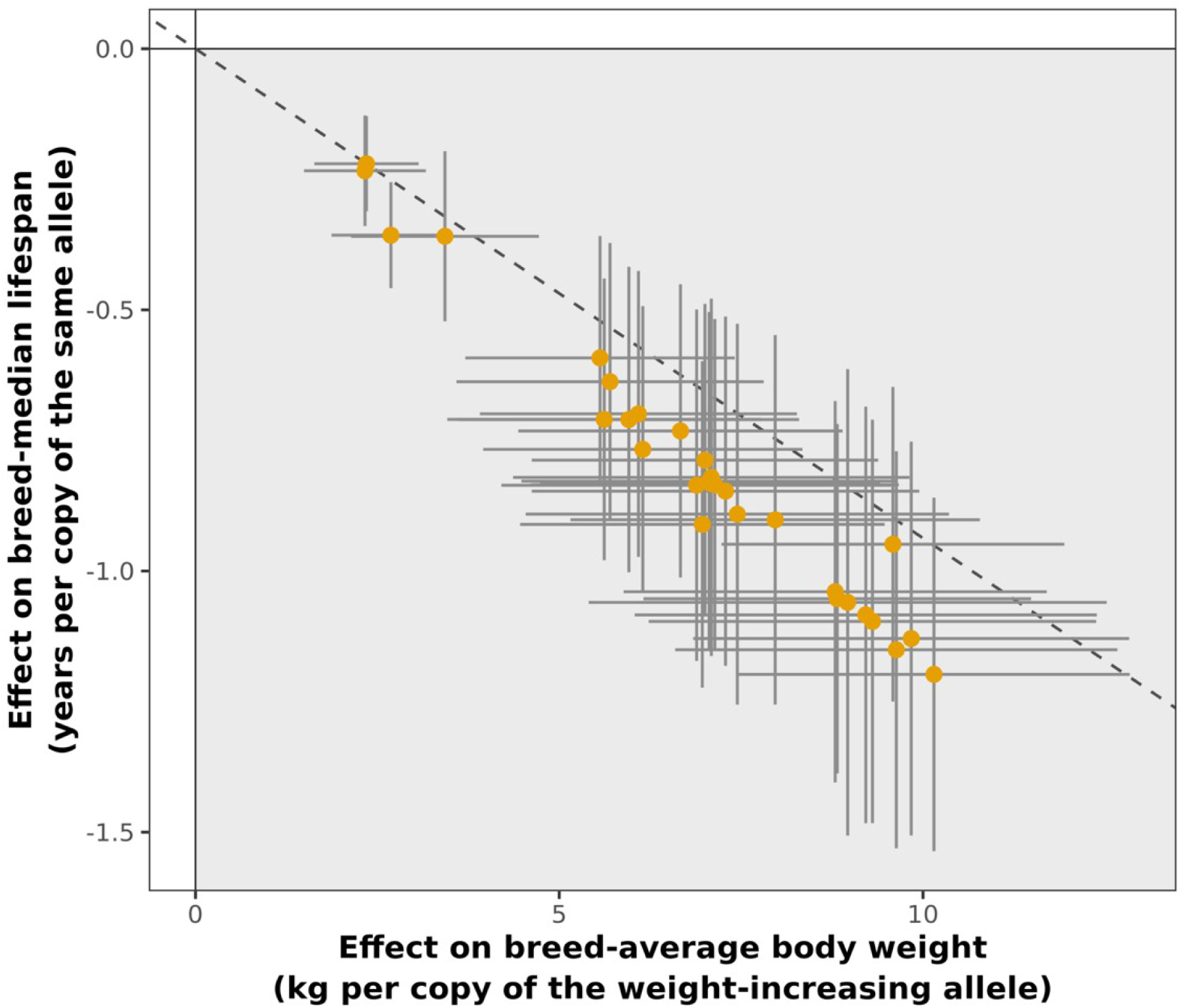
Effects of the 30 breed-average body weight loci shared with breed-median lifespan. Each point is one locus, measured at its breed-weight lead variant. The x-axis gives the effect of the weight-increasing allele on breed-average body weight (kg per copy), and the y-axis the effect of the same allele on breed-median lifespan (years per copy), with 95% confidence intervals. The shaded area marks effects in the direction of the size–lifespan trade-off. The dashed line shows the across-breed regression of breed-median lifespan on breed-average body weight (Fig 1A).

**Fig S4.**
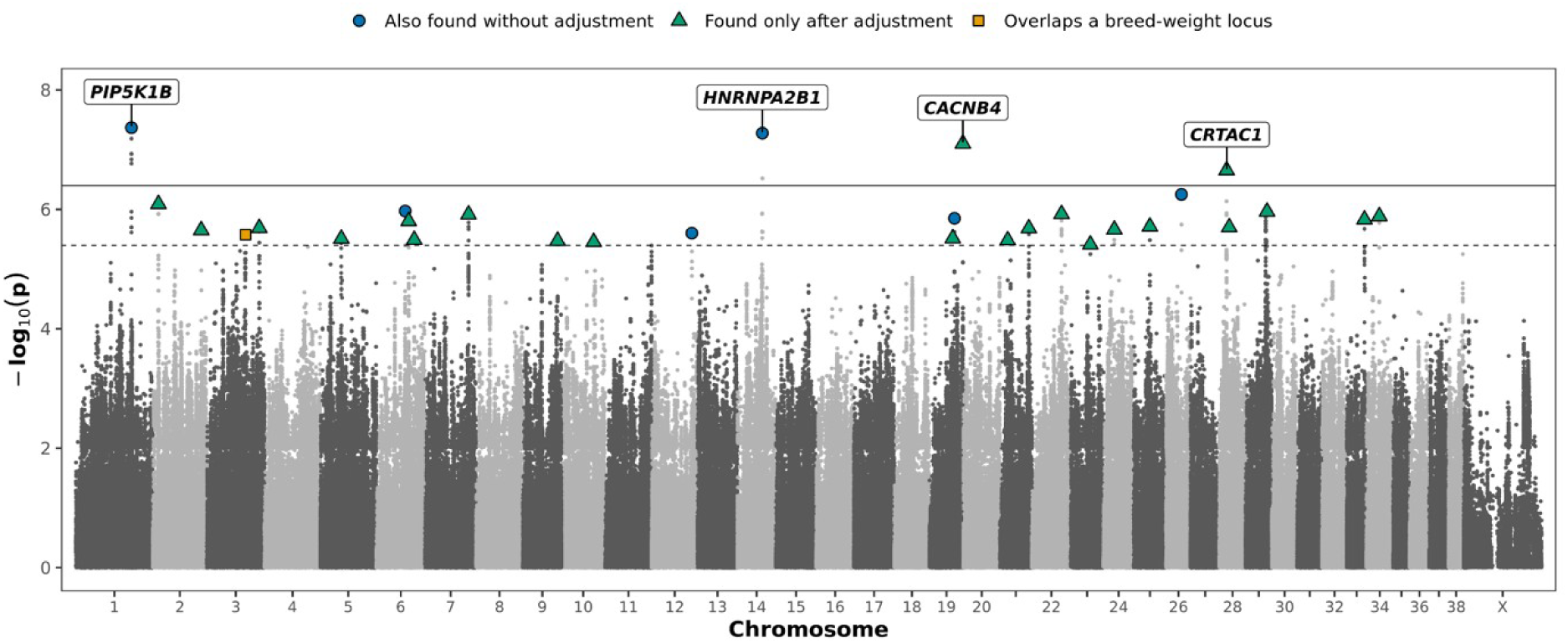
GWAS of breed-median lifespan adjusted for breed-average body weight. Manhattan plot of the 540 dogs of the Cani-DNA discovery cohort, with breed-average body weight as a covariate. Solid and dashed horizontal lines mark the genome-wide significance (p = 4 × 10⁻⁷) and suggestive (p = 4 × 10⁻⁶) thresholds. The lead variants of the 29 loci are marked by their relation to the unadjusted analyses. Loci overlapping a lifespan-specific locus are shown as blue circles. Loci overlapping a breed-weight locus or a breed-lifespan locus shared with weight are shown as orange triangles. Loci detected only after adjustment are shown as green squares. The four genome-wide significant loci are labelled with their nearest gene.

**Fig S5.**
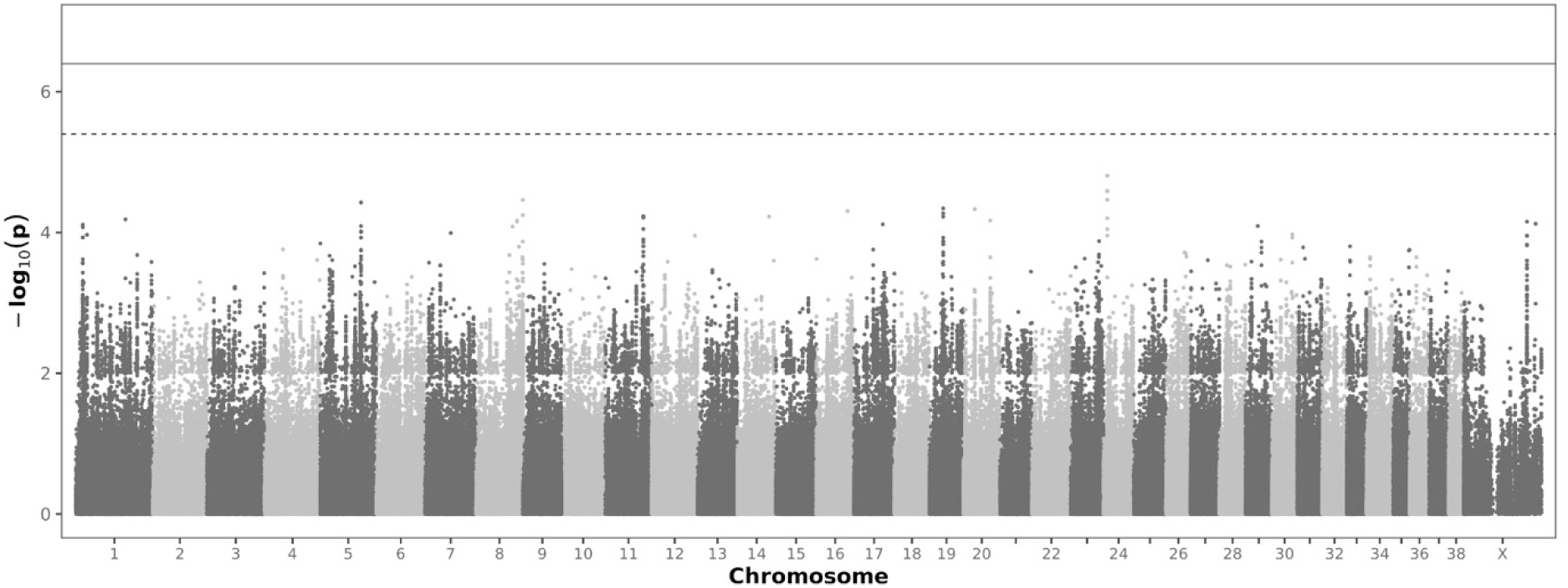
GWAS of old/young status in the Cani-DNA discovery cohort. Manhattan plot of old/young status, with breed-average body weight as a covariate, in the Cani-DNA discovery cohort. Solid and dashed horizontal lines mark the genome-wide significance (p = 4 × 10⁻⁷) and suggestive (p = 4 × 10⁻⁶) thresholds.

**Fig S6.**
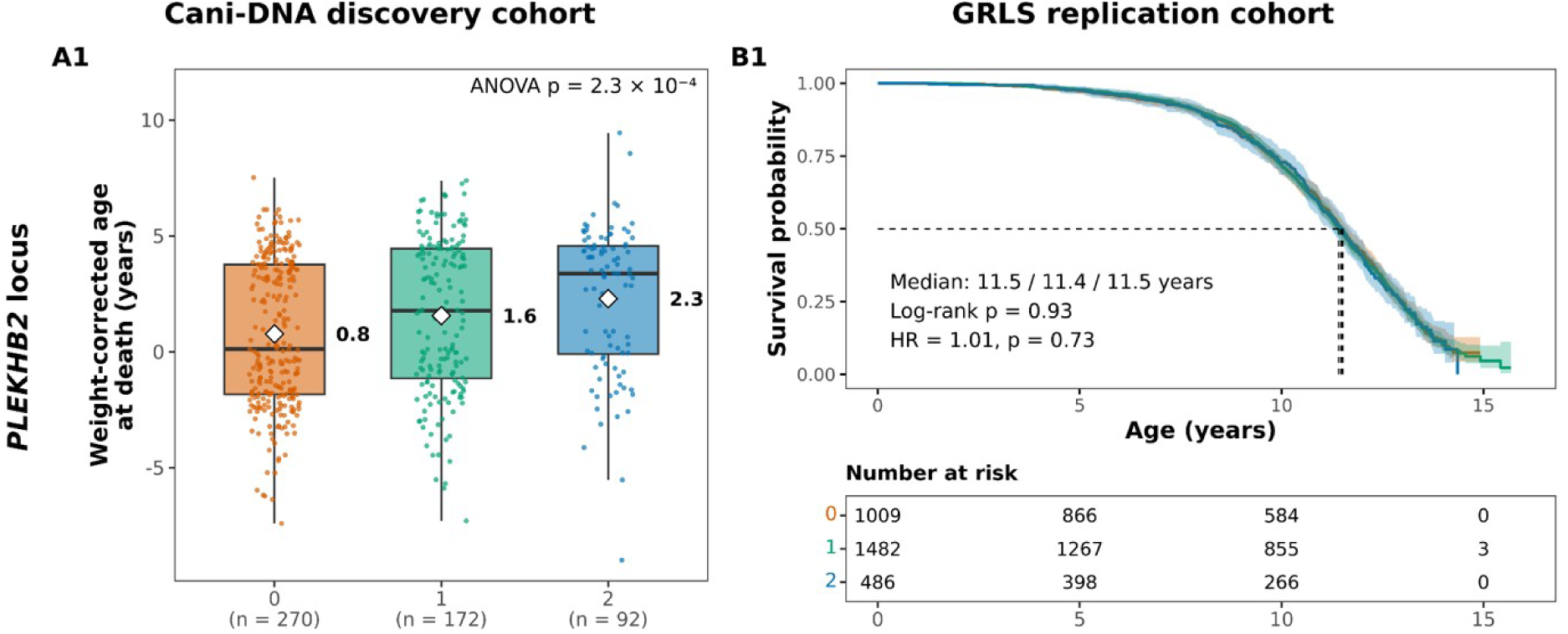
The *PLEKHB2* locus in the Cani-DNA discovery and GRLS replication cohorts. (A) Weight-corrected age at death in the Cani-DNA discovery cohort by number of copies of the effect allele at the proxy variant. (B) Kaplan–Meier survival curves in the GRLS replication cohort by genotype at the same proxy variant. Colours, boxes, diamonds, survival curves and statistics are as in Fig 5.

## Supplementary tables

**Table S1.**
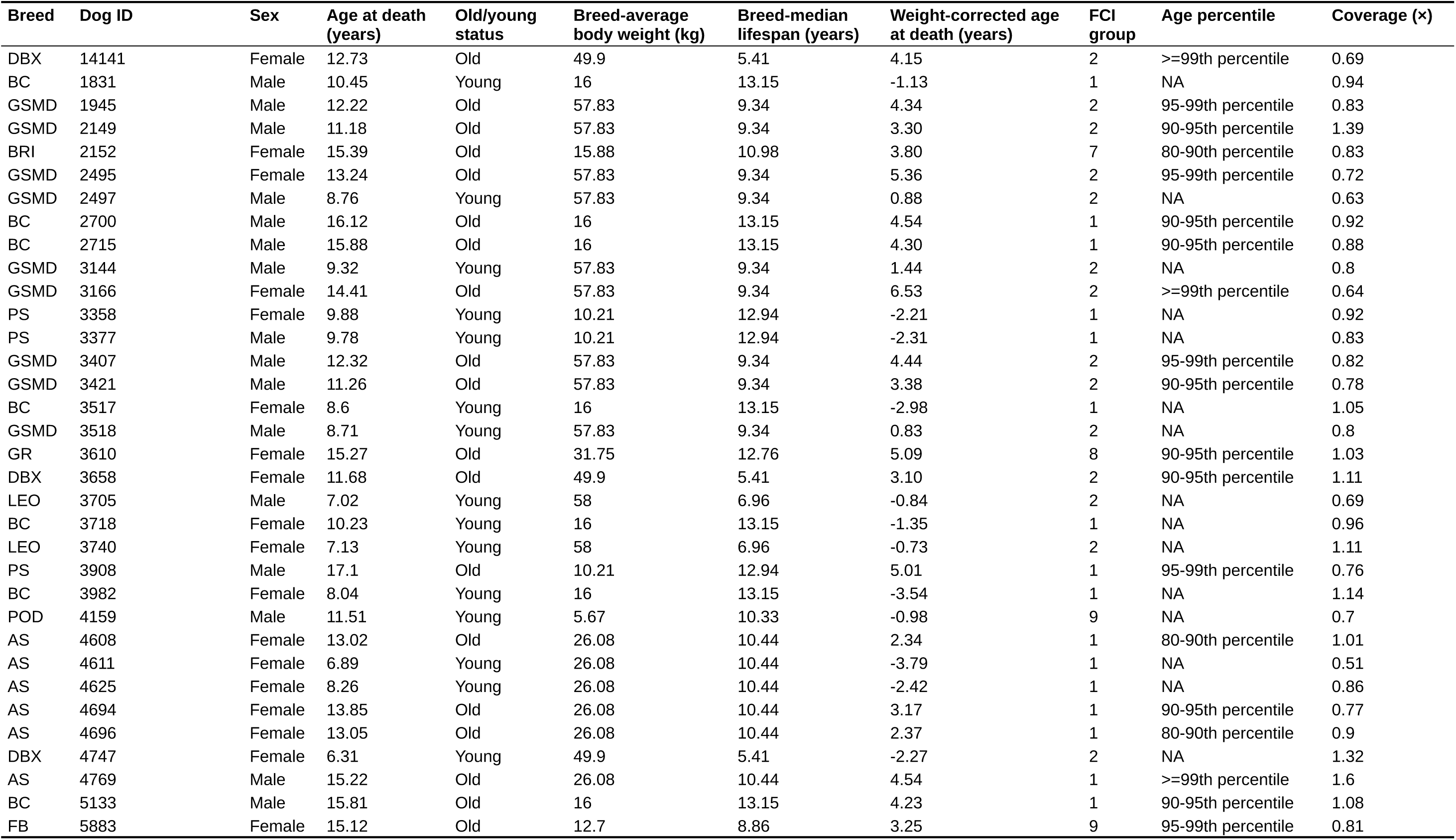
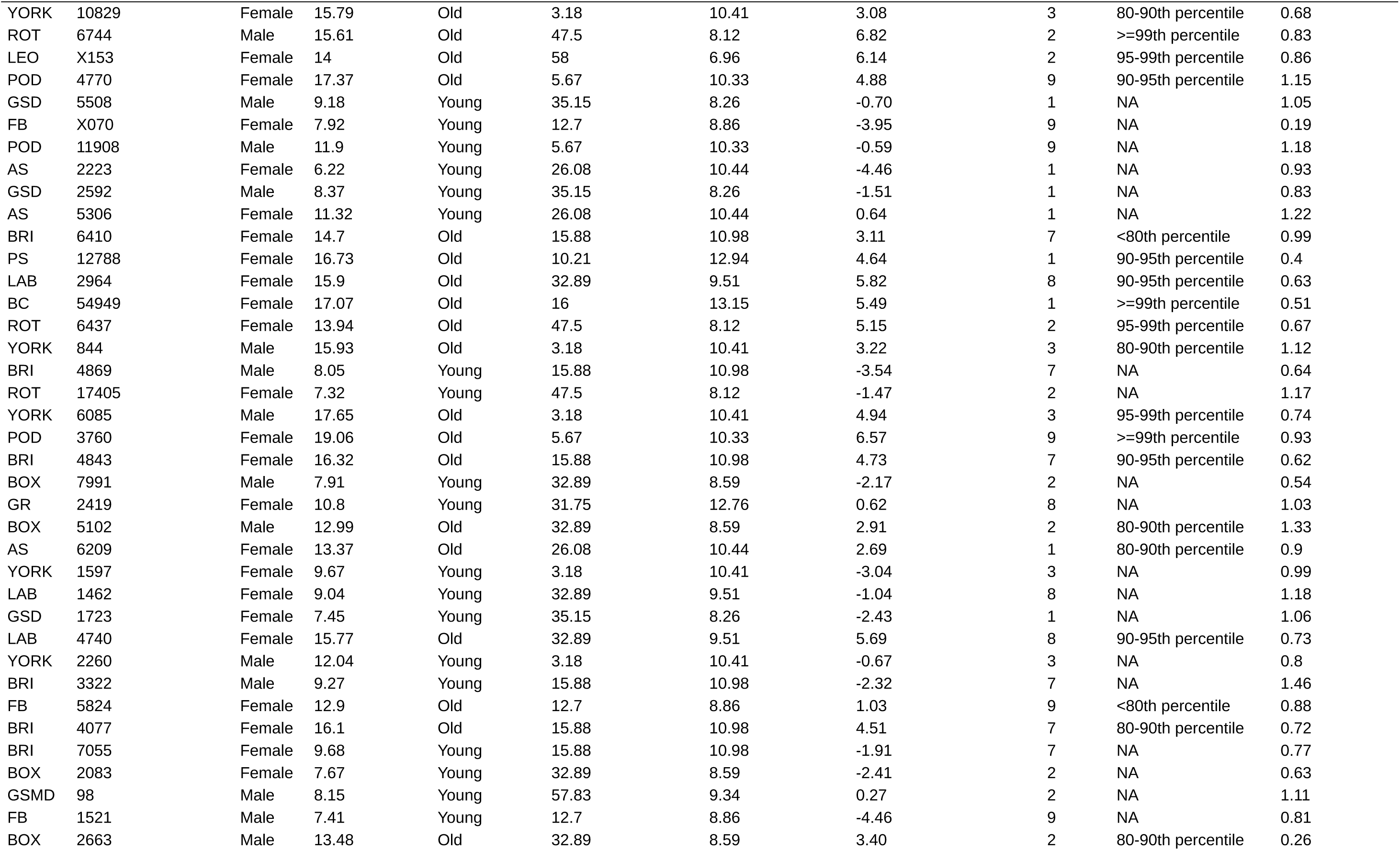
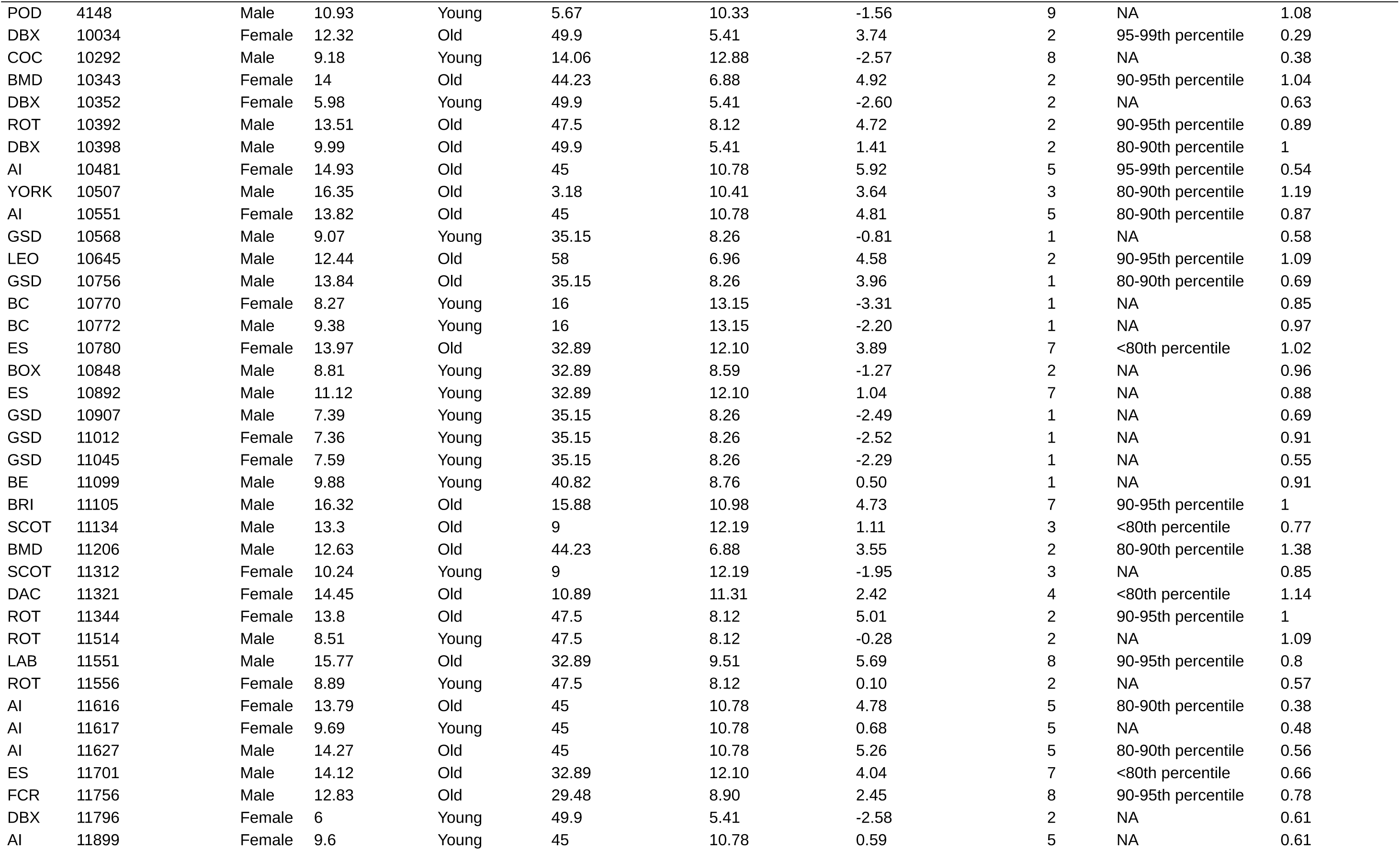
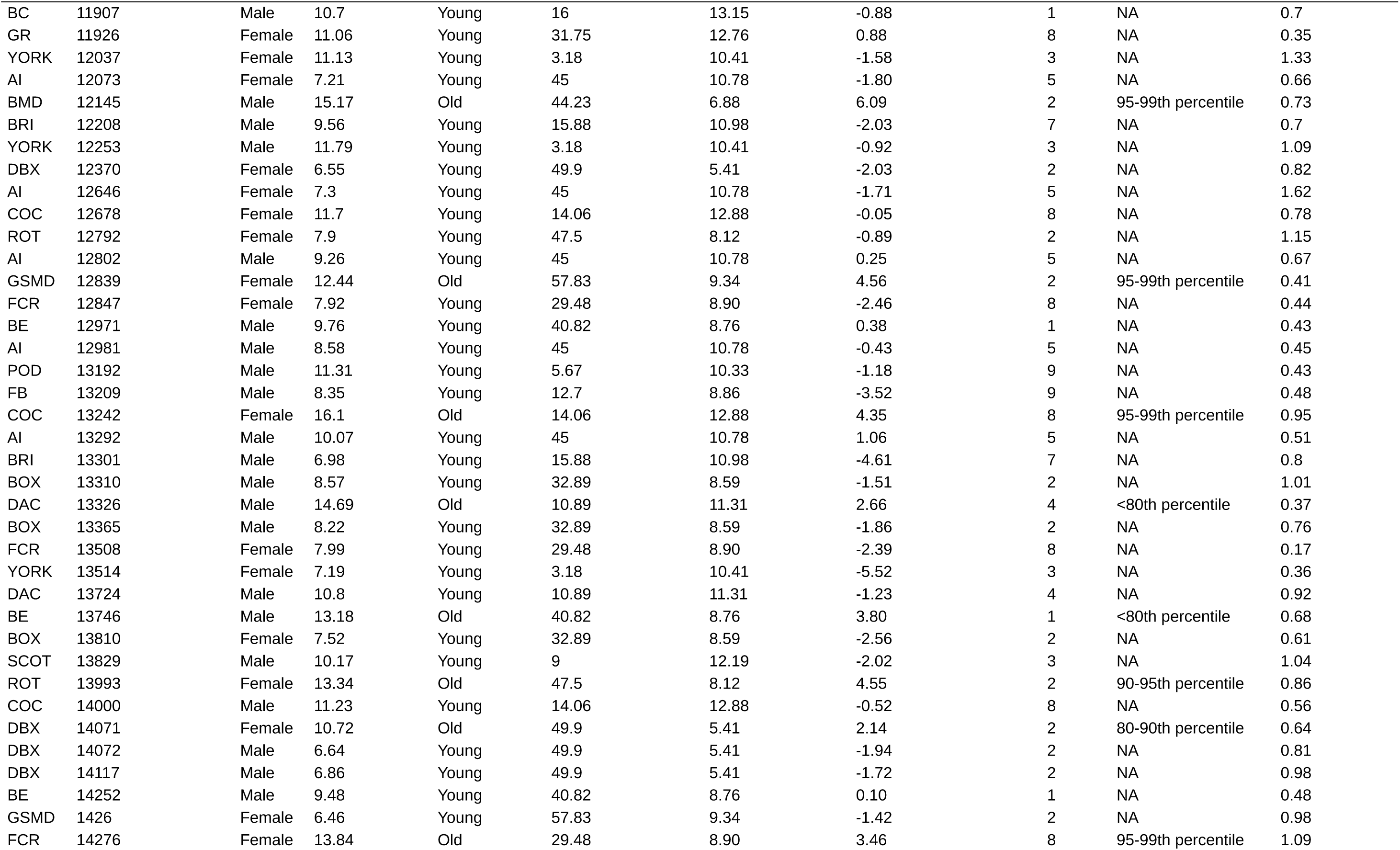
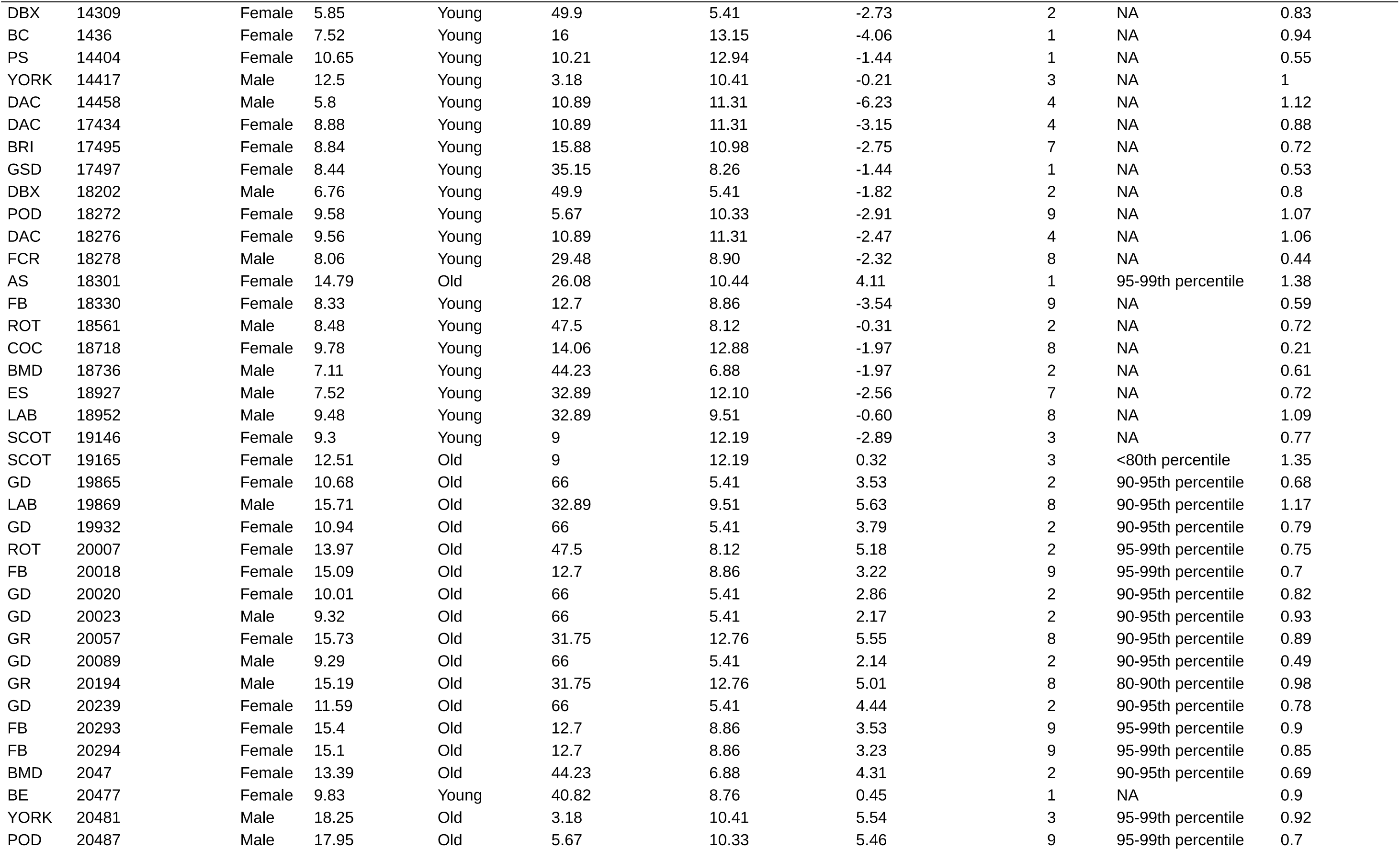
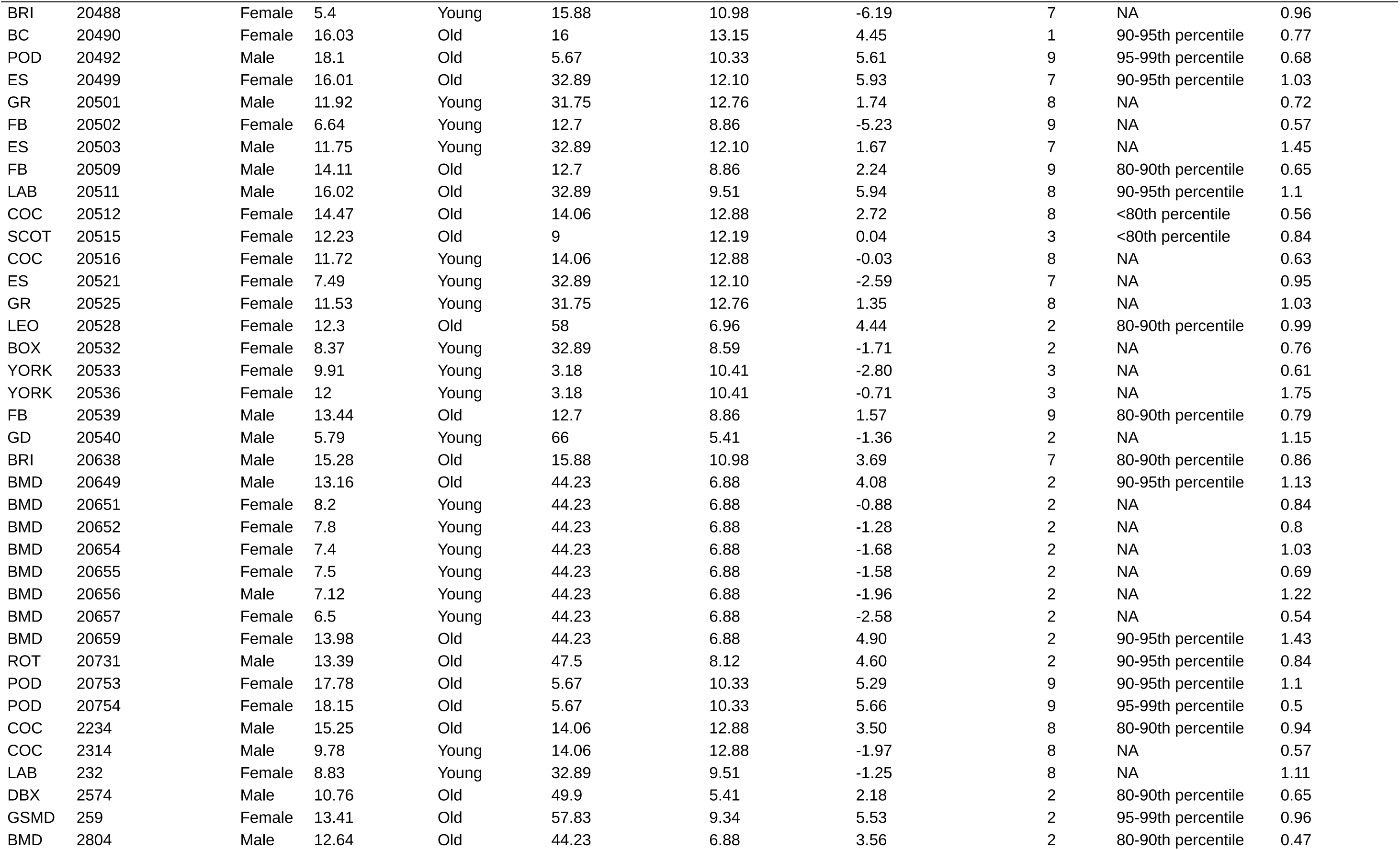
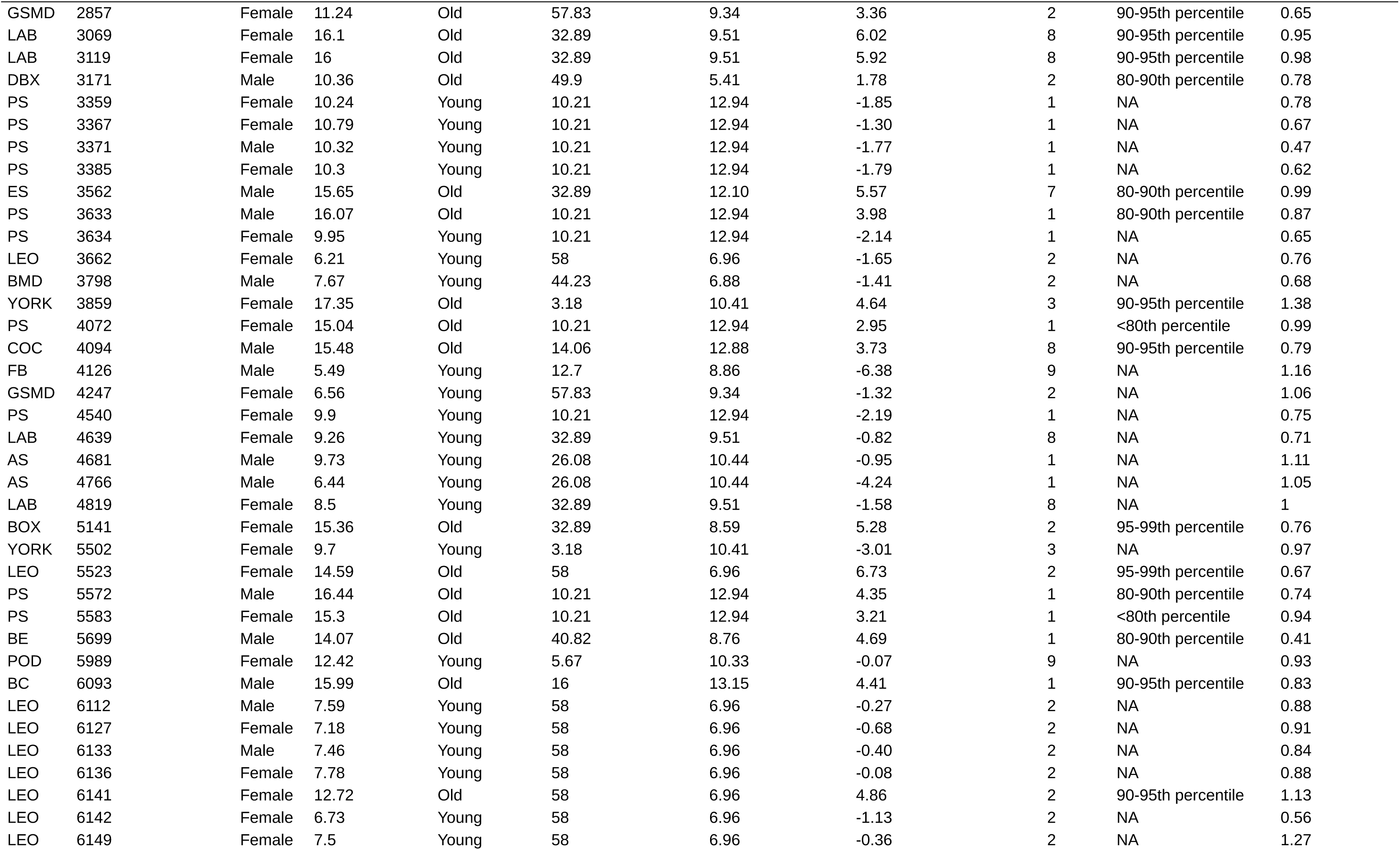
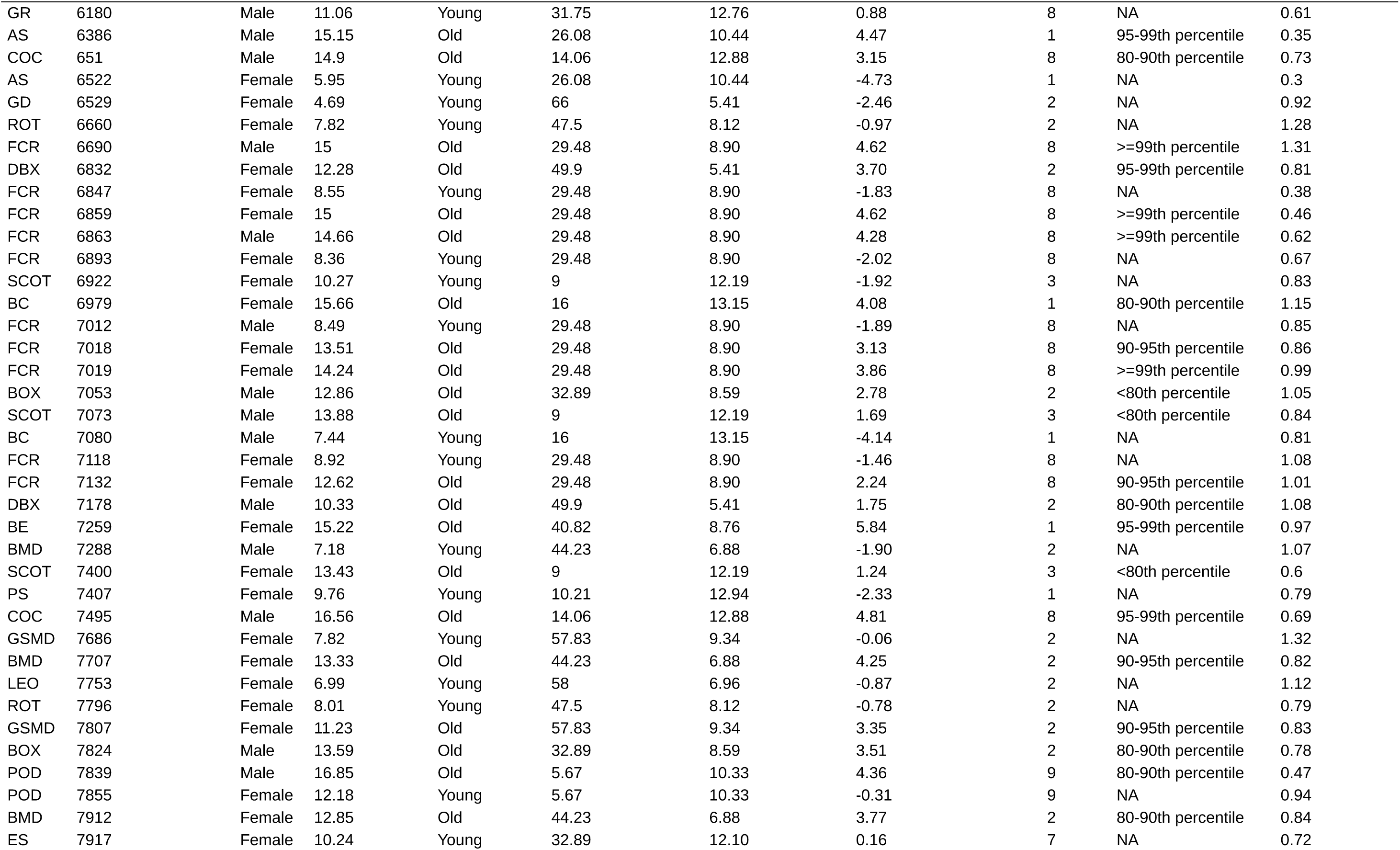
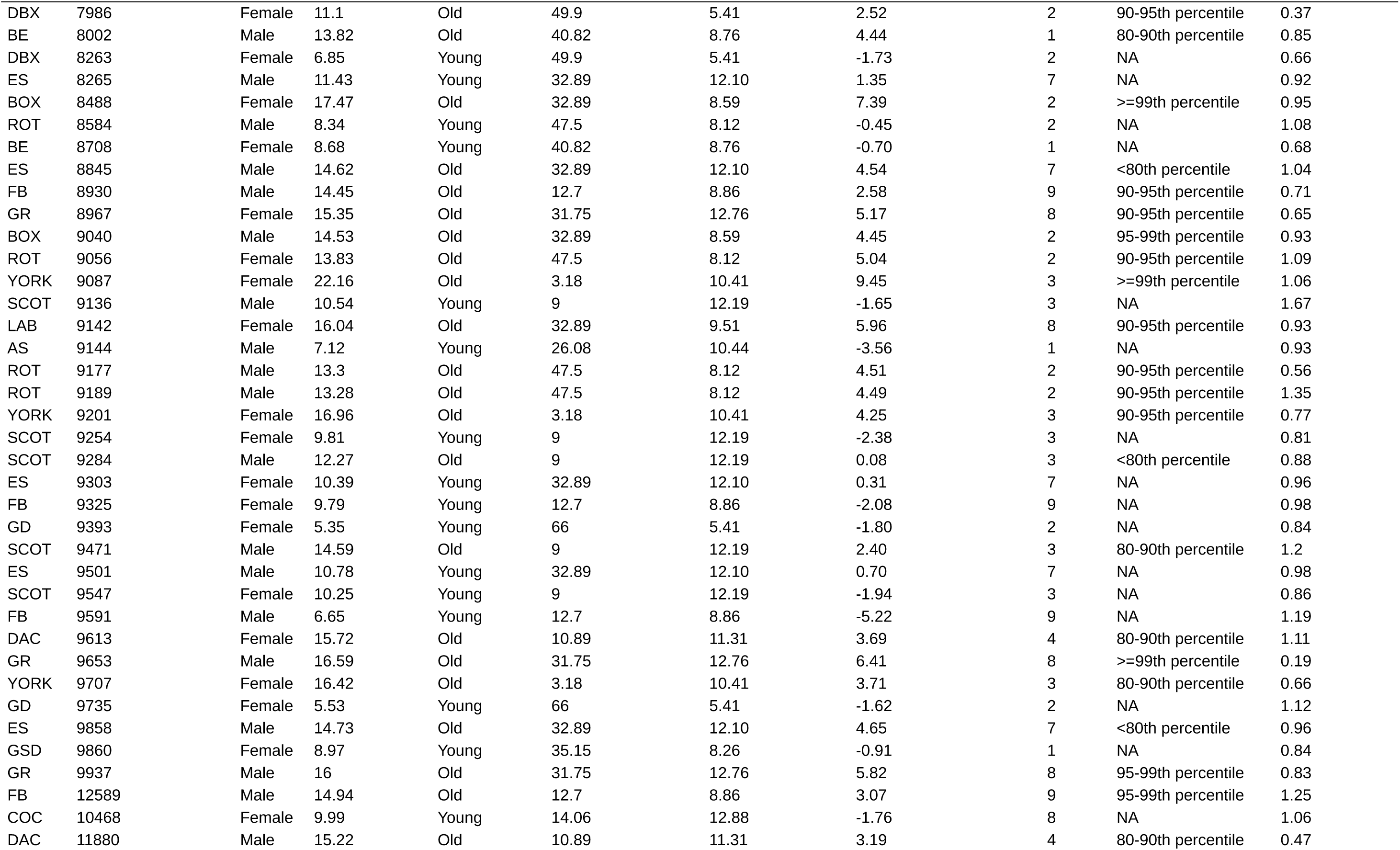
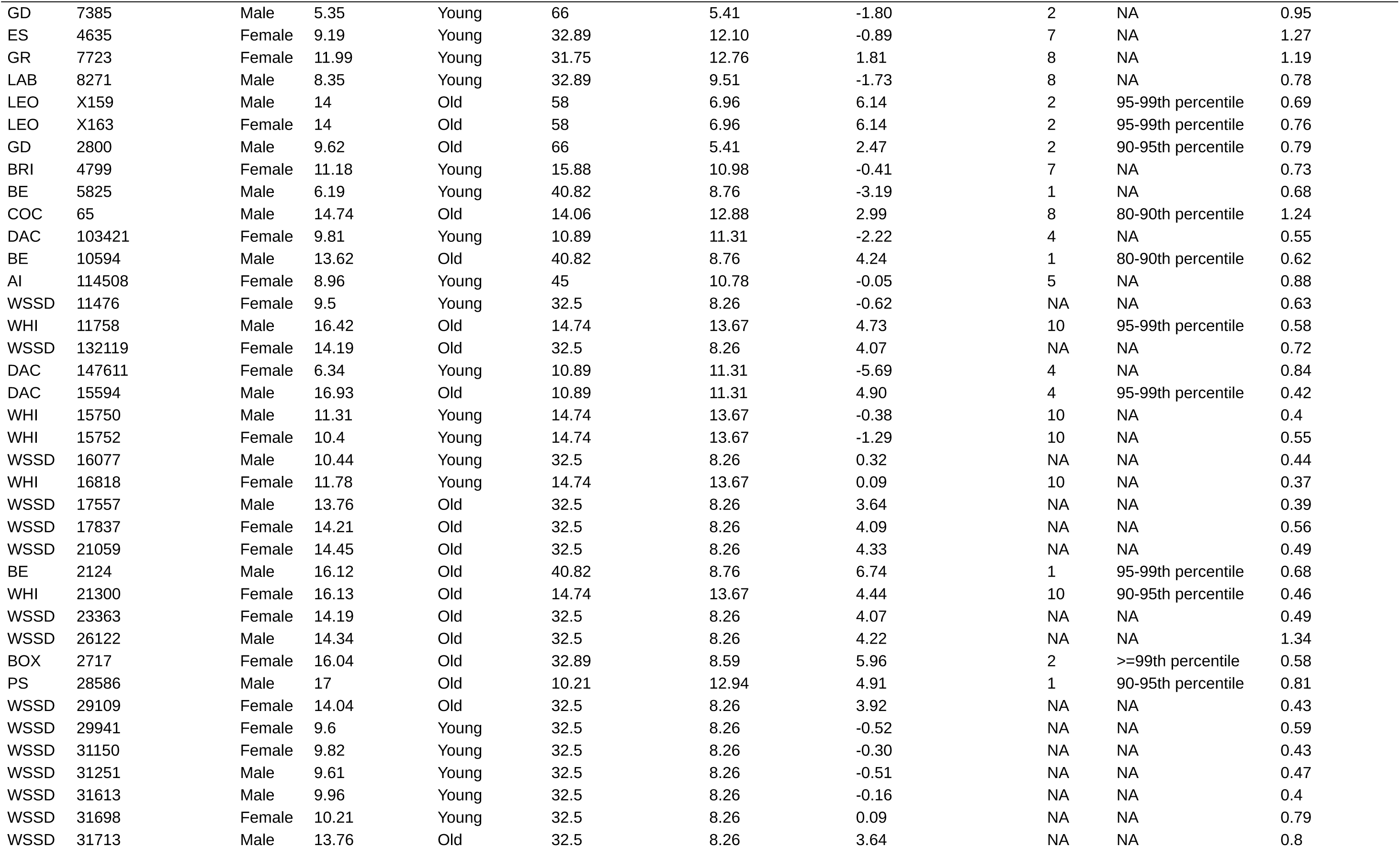
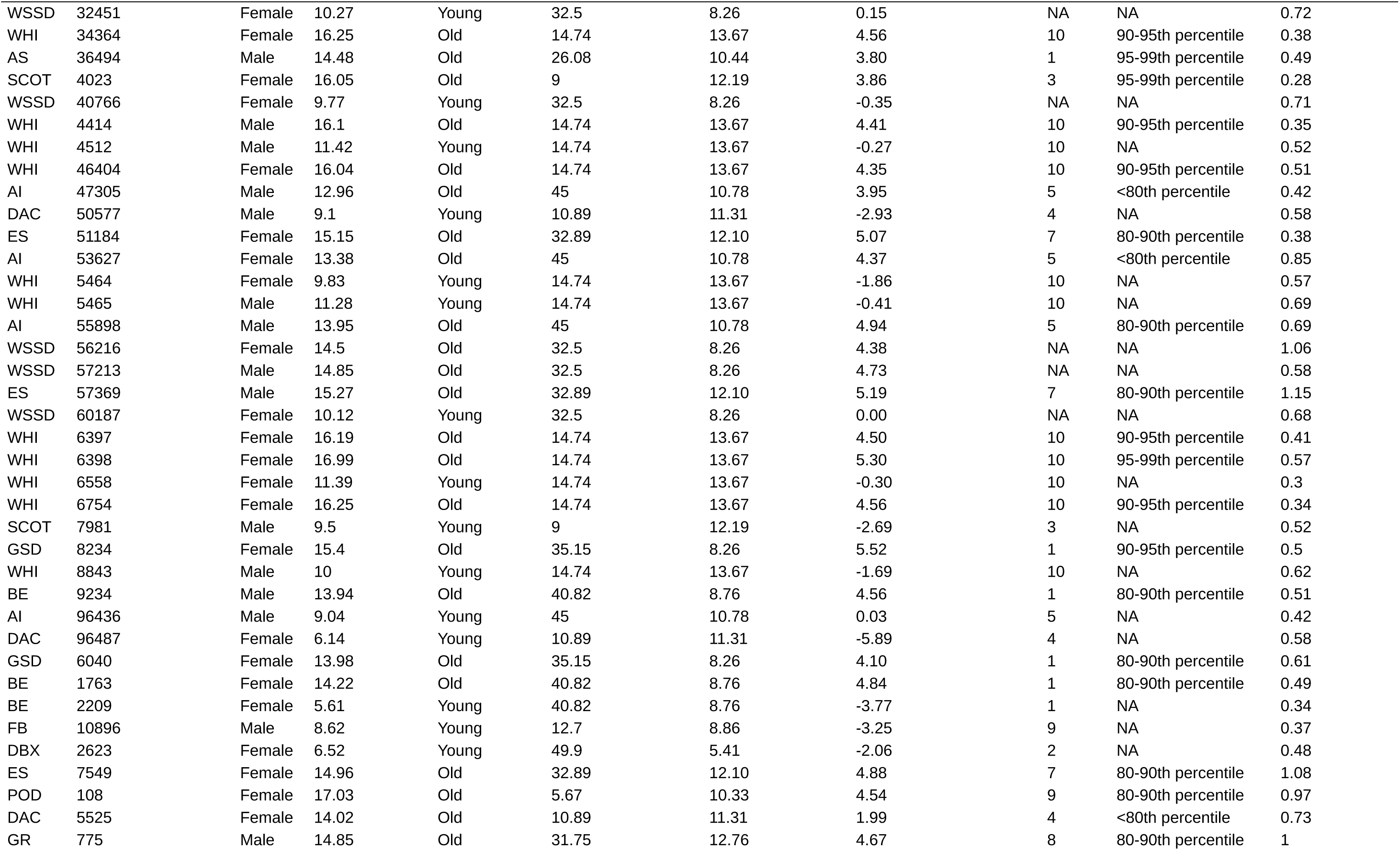
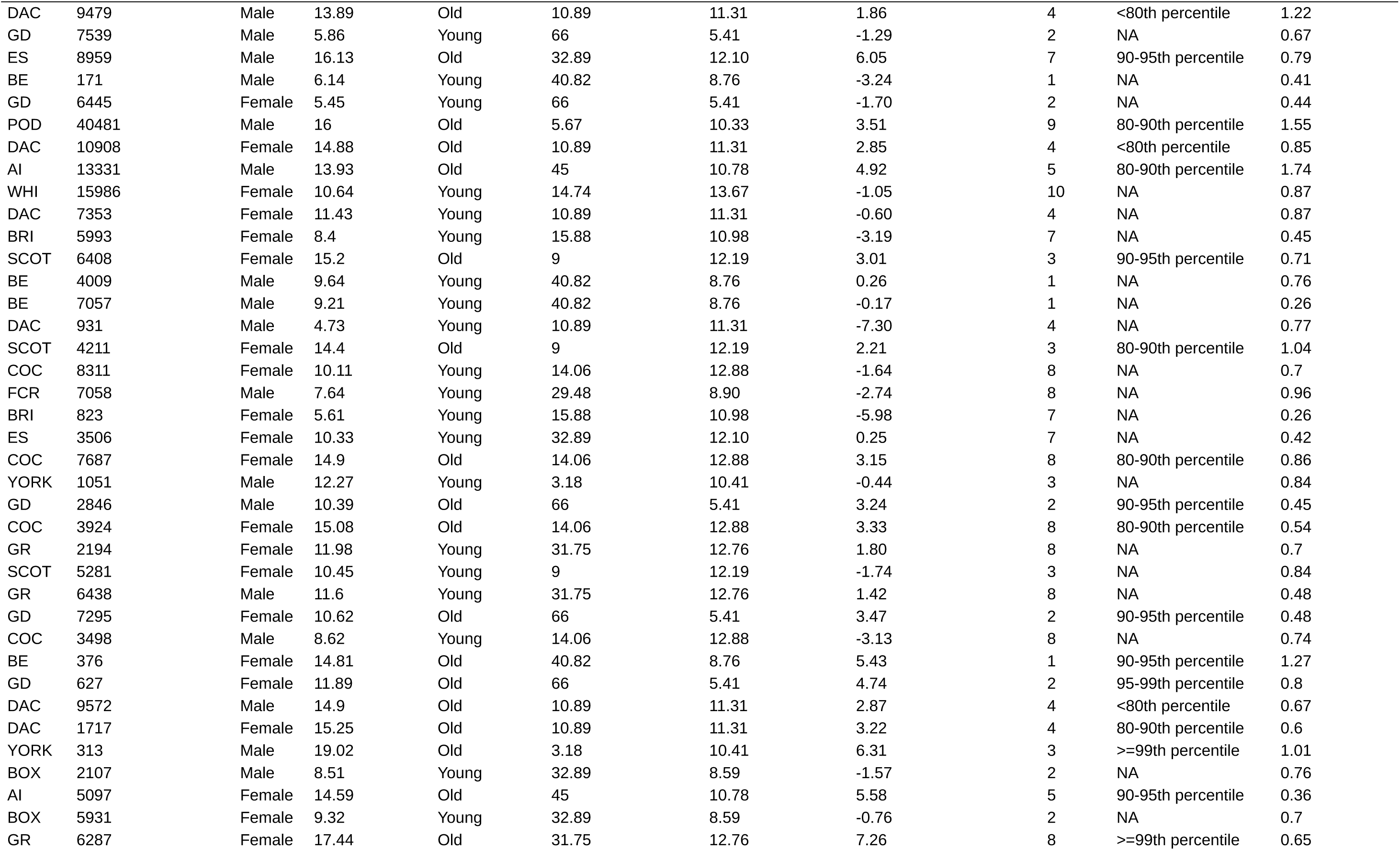
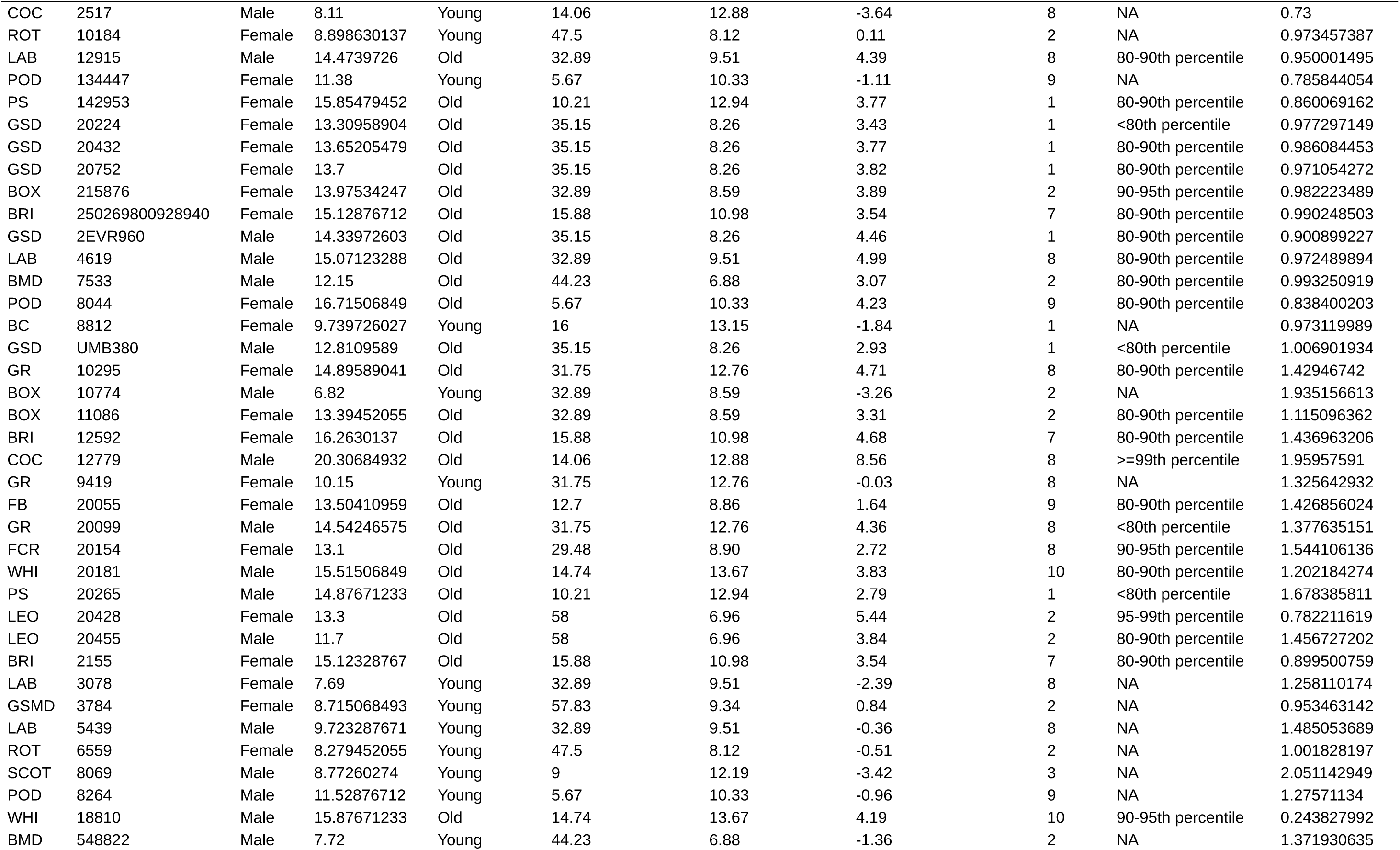
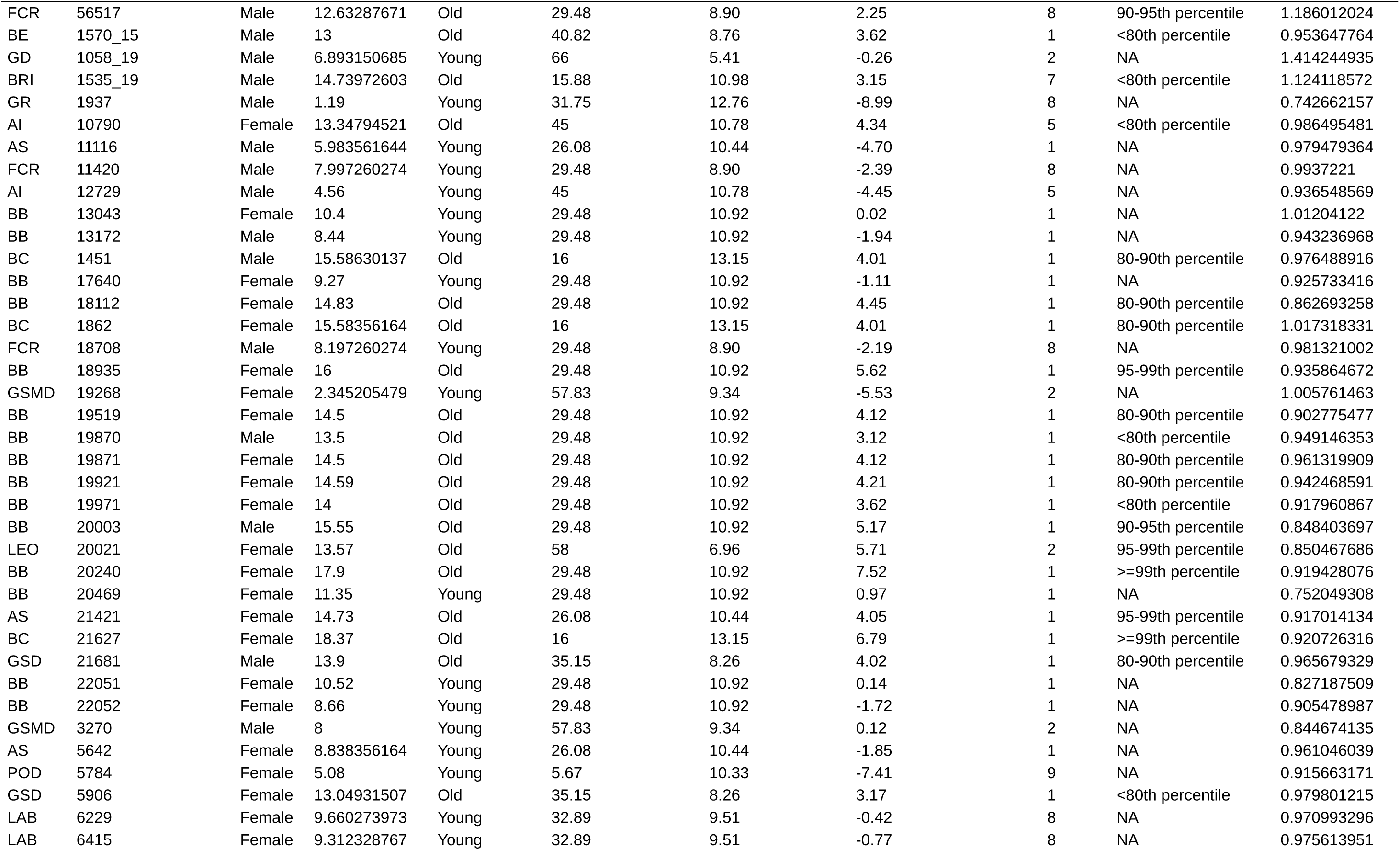
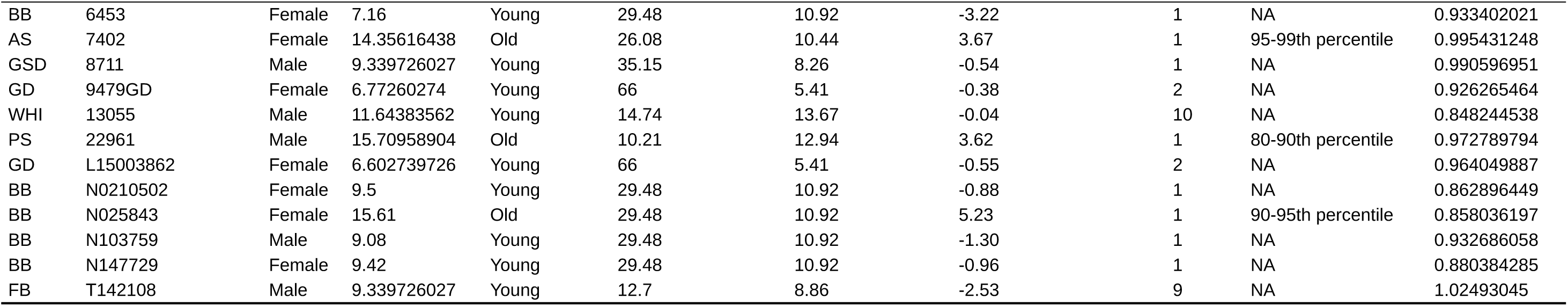
Phenotypic data of the 540 dogs of the Cani-DNA discovery cohort.

| Breed | Dog ID | Sex | Age at death (years) | Old/young status | Breed-average body weight (kg) | Breed-median lifespan (years) | Weight-corrected age at death (years) | FCI group | Age percentile | Coverage (×) |
|---|---|---|---|---|---|---|---|---|---|---|
| DBX | 14141 | Female | 12.73 | Old | 49.9 | 5.41 | 4.15 | 2 | >=99th percentile | 0.69 |
| BC | 1831 | Male | 10.45 | Young | 16 | 13.15 | -1.13 | 1 | NA | 0.94 |
| GSMD | 1945 | Male | 12.22 | Old | 57.83 | 9.34 | 4.34 | 2 | 95-99th percentile | 0.83 |
| GSMD | 2149 | Male | 11.18 | Old | 57.83 | 9.34 | 3.30 | 2 | 90-95th percentile | 1.39 |
| BRI | 2152 | Female | 15.39 | Old | 15.88 | 10.98 | 3.80 | 7 | 80-90th percentile | 0.83 |
| GSMD | 2495 | Female | 13.24 | Old | 57.83 | 9.34 | 5.36 | 2 | 95-99th percentile | 0.72 |
| GSMD | 2497 | Male | 8.76 | Young | 57.83 | 9.34 | 0.88 | 2 | NA | 0.63 |
| BC | 2700 | Male | 16.12 | Old | 16 | 13.15 | 4.54 | 1 | 90-95th percentile | 0.92 |
| BC | 2715 | Male | 15.88 | Old | 16 | 13.15 | 4.30 | 1 | 90-95th percentile | 0.88 |
| GSMD | 3144 | Male | 9.32 | Young | 57.83 | 9.34 | 1.44 | 2 | NA | 0.8 |
| GSMD | 3166 | Female | 14.41 | Old | 57.83 | 9.34 | 6.53 | 2 | >=99th percentile | 0.64 |
| PS | 3358 | Female | 9.88 | Young | 10.21 | 12.94 | -2.21 | 1 | NA | 0.92 |
| PS | 3377 | Male | 9.78 | Young | 10.21 | 12.94 | -2.31 | 1 | NA | 0.83 |
| GSMD | 3407 | Male | 12.32 | Old | 57.83 | 9.34 | 4.44 | 2 | 95-99th percentile | 0.82 |
| GSMD | 3421 | Male | 11.26 | Old | 57.83 | 9.34 | 3.38 | 2 | 90-95th percentile | 0.78 |
| BC | 3517 | Female | 8.6 | Young | 16 | 13.15 | -2.98 | 1 | NA | 1.05 |
| GSMD | 3518 | Male | 8.71 | Young | 57.83 | 9.34 | 0.83 | 2 | NA | 0.8 |
| GR | 3610 | Female | 15.27 | Old | 31.75 | 12.76 | 5.09 | 8 | 90-95th percentile | 1.03 |
| DBX | 3658 | Female | 11.68 | Old | 49.9 | 5.41 | 3.10 | 2 | 90-95th percentile | 1.11 |
| LEO | 3705 | Male | 7.02 | Young | 58 | 6.96 | -0.84 | 2 | NA | 0.69 |
| BC | 3718 | Female | 10.23 | Young | 16 | 13.15 | -1.35 | 1 | NA | 0.96 |
| LEO | 3740 | Female | 7.13 | Young | 58 | 6.96 | -0.73 | 2 | NA | 1.11 |
| PS | 3908 | Male | 17.1 | Old | 10.21 | 12.94 | 5.01 | 1 | 95-99th percentile | 0.76 |
| BC | 3982 | Female | 8.04 | Young | 16 | 13.15 | -3.54 | 1 | NA | 1.14 |
| POD | 4159 | Male | 11.51 | Young | 5.67 | 10.33 | -0.98 | 9 | NA | 0.7 |
| AS | 4608 | Female | 13.02 | Old | 26.08 | 10.44 | 2.34 | 1 | 80-90th percentile | 1.01 |
| AS | 4611 | Female | 6.89 | Young | 26.08 | 10.44 | -3.79 | 1 | NA | 0.51 |
| AS | 4625 | Female | 8.26 | Young | 26.08 | 10.44 | -2.42 | 1 | NA | 0.86 |
| AS | 4694 | Female | 13.85 | Old | 26.08 | 10.44 | 3.17 | 1 | 90-95th percentile | 0.77 |
| AS | 4696 | Female | 13.05 | Old | 26.08 | 10.44 | 2.37 | 1 | 80-90th percentile | 0.9 |
| DBX | 4747 | Female | 6.31 | Young | 49.9 | 5.41 | -2.27 | 2 | NA | 1.32 |
| AS | 4769 | Male | 15.22 | Old | 26.08 | 10.44 | 4.54 | 1 | >=99th percentile | 1.6 |
| BC | 5133 | Male | 15.81 | Old | 16 | 13.15 | 4.23 | 1 | 90-95th percentile | 1.08 |
| FB | 5883 | Female | 15.12 | Old | 12.7 | 8.86 | 3.25 | 9 | 95-99th percentile | 0.81 |
| YORK | 10829 | Female | 15.79 | Old | 3.18 | 10.41 | 3.08 | 3 | 80-90th percentile | 0.68 |
| ROT | 6744 | Male | 15.61 | Old | 47.5 | 8.12 | 6.82 | 2 | >=99th percentile | 0.83 |
| LEO | X153 | Female | 14 | Old | 58 | 6.96 | 6.14 | 2 | 95-99th percentile | 0.86 |
| POD | 4770 | Female | 17.37 | Old | 5.67 | 10.33 | 4.88 | 9 | 90-95th percentile | 1.15 |
| GSD | 5508 | Male | 9.18 | Young | 35.15 | 8.26 | -0.70 | 1 | NA | 1.05 |
| FB | X070 | Female | 7.92 | Young | 12.7 | 8.86 | -3.95 | 9 | NA | 0.19 |
| POD | 11908 | Male | 11.9 | Young | 5.67 | 10.33 | -0.59 | 9 | NA | 1.18 |
| AS | 2223 | Female | 6.22 | Young | 26.08 | 10.44 | -4.46 | 1 | NA | 0.93 |
| GSD | 2592 | Male | 8.37 | Young | 35.15 | 8.26 | -1.51 | 1 | NA | 0.83 |
| AS | 5306 | Female | 11.32 | Young | 26.08 | 10.44 | 0.64 | 1 | NA | 1.22 |
| BRI | 6410 | Female | 14.7 | Old | 15.88 | 10.98 | 3.11 | 7 | <80th percentile | 0.99 |
| PS | 12788 | Female | 16.73 | Old | 10.21 | 12.94 | 4.64 | 1 | 90-95th percentile | 0.4 |
| LAB | 2964 | Female | 15.9 | Old | 32.89 | 9.51 | 5.82 | 8 | 90-95th percentile | 0.63 |
| BC | 54949 | Female | 17.07 | Old | 16 | 13.15 | 5.49 | 1 | >=99th percentile | 0.51 |
| ROT | 6437 | Female | 13.94 | Old | 47.5 | 8.12 | 5.15 | 2 | 95-99th percentile | 0.67 |
| YORK | 844 | Male | 15.93 | Old | 3.18 | 10.41 | 3.22 | 3 | 80-90th percentile | 1.12 |
| BRI | 4869 | Male | 8.05 | Young | 15.88 | 10.98 | -3.54 | 7 | NA | 0.64 |
| ROT | 17405 | Female | 7.32 | Young | 47.5 | 8.12 | -1.47 | 2 | NA | 1.17 |
| YORK | 6085 | Male | 17.65 | Old | 3.18 | 10.41 | 4.94 | 3 | 95-99th percentile | 0.74 |
| POD | 3760 | Female | 19.06 | Old | 5.67 | 10.33 | 6.57 | 9 | >=99th percentile | 0.93 |
| BRI | 4843 | Female | 16.32 | Old | 15.88 | 10.98 | 4.73 | 7 | 90-95th percentile | 0.62 |
| BOX | 7991 | Male | 7.91 | Young | 32.89 | 8.59 | -2.17 | 2 | NA | 0.54 |
| GR | 2419 | Female | 10.8 | Young | 31.75 | 12.76 | 0.62 | 8 | NA | 1.03 |
| BOX | 5102 | Male | 12.99 | Old | 32.89 | 8.59 | 2.91 | 2 | 80-90th percentile | 1.33 |
| AS | 6209 | Female | 13.37 | Old | 26.08 | 10.44 | 2.69 | 1 | 80-90th percentile | 0.9 |
| YORK | 1597 | Female | 9.67 | Young | 3.18 | 10.41 | -3.04 | 3 | NA | 0.99 |
| LAB | 1462 | Female | 9.04 | Young | 32.89 | 9.51 | -1.04 | 8 | NA | 1.18 |
| GSD | 1723 | Female | 7.45 | Young | 35.15 | 8.26 | -2.43 | 1 | NA | 1.06 |
| LAB | 4740 | Female | 15.77 | Old | 32.89 | 9.51 | 5.69 | 8 | 90-95th percentile | 0.73 |
| YORK | 2260 | Male | 12.04 | Young | 3.18 | 10.41 | -0.67 | 3 | NA | 0.8 |
| BRI | 3322 | Male | 9.27 | Young | 15.88 | 10.98 | -2.32 | 7 | NA | 1.46 |
| FB | 5824 | Female | 12.9 | Old | 12.7 | 8.86 | 1.03 | 9 | <80th percentile | 0.88 |
| BRI | 4077 | Female | 16.1 | Old | 15.88 | 10.98 | 4.51 | 7 | 80-90th percentile | 0.72 |
| BRI | 7055 | Female | 9.68 | Young | 15.88 | 10.98 | -1.91 | 7 | NA | 0.77 |
| BOX | 2083 | Female | 7.67 | Young | 32.89 | 8.59 | -2.41 | 2 | NA | 0.63 |
| GSMD | 98 | Male | 8.15 | Young | 57.83 | 9.34 | 0.27 | 2 | NA | 1.11 |
| FB | 1521 | Male | 7.41 | Young | 12.7 | 8.86 | -4.46 | 9 | NA | 0.81 |
| BOX | 2663 | Male | 13.48 | Old | 32.89 | 8.59 | 3.40 | 2 | 80-90th percentile | 0.26 |
| POD | 4148 | Male | 10.93 | Young | 5.67 | 10.33 | -1.56 | 9 | NA | 1.08 |
| DBX | 10034 | Female | 12.32 | Old | 49.9 | 5.41 | 3.74 | 2 | 95-99th percentile | 0.29 |
| COC | 10292 | Male | 9.18 | Young | 14.06 | 12.88 | -2.57 | 8 | NA | 0.38 |
| BMD | 10343 | Female | 14 | Old | 44.23 | 6.88 | 4.92 | 2 | 90-95th percentile | 1.04 |
| DBX | 10352 | Female | 5.98 | Young | 49.9 | 5.41 | -2.60 | 2 | NA | 0.63 |
| ROT | 10392 | Male | 13.51 | Old | 47.5 | 8.12 | 4.72 | 2 | 90-95th percentile | 0.89 |
| DBX | 10398 | Male | 9.99 | Old | 49.9 | 5.41 | 1.41 | 2 | 80-90th percentile | 1 |
| AI | 10481 | Female | 14.93 | Old | 45 | 10.78 | 5.92 | 5 | 95-99th percentile | 0.54 |
| YORK | 10507 | Male | 16.35 | Old | 3.18 | 10.41 | 3.64 | 3 | 80-90th percentile | 1.19 |
| AI | 10551 | Female | 13.82 | Old | 45 | 10.78 | 4.81 | 5 | 80-90th percentile | 0.87 |
| GSD | 10568 | Male | 9.07 | Young | 35.15 | 8.26 | -0.81 | 1 | NA | 0.58 |
| LEO | 10645 | Male | 12.44 | Old | 58 | 6.96 | 4.58 | 2 | 90-95th percentile | 1.09 |
| GSD | 10756 | Male | 13.84 | Old | 35.15 | 8.26 | 3.96 | 1 | 80-90th percentile | 0.69 |
| BC | 10770 | Female | 8.27 | Young | 16 | 13.15 | -3.31 | 1 | NA | 0.85 |
| BC | 10772 | Male | 9.38 | Young | 16 | 13.15 | -2.20 | 1 | NA | 0.97 |
| ES | 10780 | Female | 13.97 | Old | 32.89 | 12.10 | 3.89 | 7 | <80th percentile | 1.02 |
| BOX | 10848 | Male | 8.81 | Young | 32.89 | 8.59 | -1.27 | 2 | NA | 0.96 |
| ES | 10892 | Male | 11.12 | Young | 32.89 | 12.10 | 1.04 | 7 | NA | 0.88 |
| GSD | 10907 | Male | 7.39 | Young | 35.15 | 8.26 | -2.49 | 1 | NA | 0.69 |
| GSD | 11012 | Female | 7.36 | Young | 35.15 | 8.26 | -2.52 | 1 | NA | 0.91 |
| GSD | 11045 | Female | 7.59 | Young | 35.15 | 8.26 | -2.29 | 1 | NA | 0.55 |
| BE | 11099 | Male | 9.88 | Young | 40.82 | 8.76 | 0.50 | 1 | NA | 0.91 |
| BRI | 11105 | Male | 16.32 | Old | 15.88 | 10.98 | 4.73 | 7 | 90-95th percentile | 1 |
| SCOT | 11134 | Male | 13.3 | Old | 9 | 12.19 | 1.11 | 3 | <80th percentile | 0.77 |
| BMD | 11206 | Male | 12.63 | Old | 44.23 | 6.88 | 3.55 | 2 | 80-90th percentile | 1.38 |
| SCOT | 11312 | Female | 10.24 | Young | 9 | 12.19 | -1.95 | 3 | NA | 0.85 |
| DAC | 11321 | Female | 14.45 | Old | 10.89 | 11.31 | 2.42 | 4 | <80th percentile | 1.14 |
| ROT | 11344 | Female | 13.8 | Old | 47.5 | 8.12 | 5.01 | 2 | 90-95th percentile | 1 |
| ROT | 11514 | Male | 8.51 | Young | 47.5 | 8.12 | -0.28 | 2 | NA | 1.09 |
| LAB | 11551 | Male | 15.77 | Old | 32.89 | 9.51 | 5.69 | 8 | 90-95th percentile | 0.8 |
| ROT | 11556 | Female | 8.89 | Young | 47.5 | 8.12 | 0.10 | 2 | NA | 0.57 |
| AI | 11616 | Female | 13.79 | Old | 45 | 10.78 | 4.78 | 5 | 80-90th percentile | 0.38 |
| AI | 11617 | Female | 9.69 | Young | 45 | 10.78 | 0.68 | 5 | NA | 0.48 |
| AI | 11627 | Male | 14.27 | Old | 45 | 10.78 | 5.26 | 5 | 80-90th percentile | 0.56 |
| ES | 11701 | Male | 14.12 | Old | 32.89 | 12.10 | 4.04 | 7 | <80th percentile | 0.66 |
| FCR | 11756 | Male | 12.83 | Old | 29.48 | 8.90 | 2.45 | 8 | 90-95th percentile | 0.78 |
| DBX | 11796 | Female | 6 | Young | 49.9 | 5.41 | -2.58 | 2 | NA | 0.61 |
| AI | 11899 | Female | 9.6 | Young | 45 | 10.78 | 0.59 | 5 | NA | 0.61 |
| BC | 11907 | Male | 10.7 | Young | 16 | 13.15 | -0.88 | 1 | NA | 0.7 |
| GR | 11926 | Female | 11.06 | Young | 31.75 | 12.76 | 0.88 | 8 | NA | 0.35 |
| YORK | 12037 | Female | 11.13 | Young | 3.18 | 10.41 | -1.58 | 3 | NA | 1.33 |
| AI | 12073 | Female | 7.21 | Young | 45 | 10.78 | -1.80 | 5 | NA | 0.66 |
| BMD | 12145 | Male | 15.17 | Old | 44.23 | 6.88 | 6.09 | 2 | 95-99th percentile | 0.73 |
| BRI | 12208 | Male | 9.56 | Young | 15.88 | 10.98 | -2.03 | 7 | NA | 0.7 |
| YORK | 12253 | Male | 11.79 | Young | 3.18 | 10.41 | -0.92 | 3 | NA | 1.09 |
| DBX | 12370 | Female | 6.55 | Young | 49.9 | 5.41 | -2.03 | 2 | NA | 0.82 |
| AI | 12646 | Female | 7.3 | Young | 45 | 10.78 | -1.71 | 5 | NA | 1.62 |
| COC | 12678 | Female | 11.7 | Young | 14.06 | 12.88 | -0.05 | 8 | NA | 0.78 |
| ROT | 12792 | Female | 7.9 | Young | 47.5 | 8.12 | -0.89 | 2 | NA | 1.15 |
| AI | 12802 | Male | 9.26 | Young | 45 | 10.78 | 0.25 | 5 | NA | 0.67 |
| GSMD | 12839 | Female | 12.44 | Old | 57.83 | 9.34 | 4.56 | 2 | 95-99th percentile | 0.41 |
| FCR | 12847 | Female | 7.92 | Young | 29.48 | 8.90 | -2.46 | 8 | NA | 0.44 |
| BE | 12971 | Male | 9.76 | Young | 40.82 | 8.76 | 0.38 | 1 | NA | 0.43 |
| AI | 12981 | Male | 8.58 | Young | 45 | 10.78 | -0.43 | 5 | NA | 0.45 |
| POD | 13192 | Male | 11.31 | Young | 5.67 | 10.33 | -1.18 | 9 | NA | 0.43 |
| FB | 13209 | Male | 8.35 | Young | 12.7 | 8.86 | -3.52 | 9 | NA | 0.48 |
| COC | 13242 | Female | 16.1 | Old | 14.06 | 12.88 | 4.35 | 8 | 95-99th percentile | 0.95 |
| AI | 13292 | Male | 10.07 | Young | 45 | 10.78 | 1.06 | 5 | NA | 0.51 |
| BRI | 13301 | Male | 6.98 | Young | 15.88 | 10.98 | -4.61 | 7 | NA | 0.8 |
| BOX | 13310 | Male | 8.57 | Young | 32.89 | 8.59 | -1.51 | 2 | NA | 1.01 |
| DAC | 13326 | Male | 14.69 | Old | 10.89 | 11.31 | 2.66 | 4 | <80th percentile | 0.37 |
| BOX | 13365 | Male | 8.22 | Young | 32.89 | 8.59 | -1.86 | 2 | NA | 0.76 |
| FCR | 13508 | Female | 7.99 | Young | 29.48 | 8.90 | -2.39 | 8 | NA | 0.17 |
| YORK | 13514 | Female | 7.19 | Young | 3.18 | 10.41 | -5.52 | 3 | NA | 0.36 |
| DAC | 13724 | Male | 10.8 | Young | 10.89 | 11.31 | -1.23 | 4 | NA | 0.92 |
| BE | 13746 | Male | 13.18 | Old | 40.82 | 8.76 | 3.80 | 1 | <80th percentile | 0.68 |
| BOX | 13810 | Female | 7.52 | Young | 32.89 | 8.59 | -2.56 | 2 | NA | 0.61 |
| SCOT | 13829 | Male | 10.17 | Young | 9 | 12.19 | -2.02 | 3 | NA | 1.04 |
| ROT | 13993 | Female | 13.34 | Old | 47.5 | 8.12 | 4.55 | 2 | 90-95th percentile | 0.86 |
| COC | 14000 | Male | 11.23 | Young | 14.06 | 12.88 | -0.52 | 8 | NA | 0.56 |
| DBX | 14071 | Female | 10.72 | Old | 49.9 | 5.41 | 2.14 | 2 | 80-90th percentile | 0.64 |
| DBX | 14072 | Male | 6.64 | Young | 49.9 | 5.41 | -1.94 | 2 | NA | 0.81 |
| DBX | 14117 | Male | 6.86 | Young | 49.9 | 5.41 | -1.72 | 2 | NA | 0.98 |
| BE | 14252 | Male | 9.48 | Young | 40.82 | 8.76 | 0.10 | 1 | NA | 0.48 |
| GSMD | 1426 | Female | 6.46 | Young | 57.83 | 9.34 | -1.42 | 2 | NA | 0.98 |
| FCR | 14276 | Female | 13.84 | Old | 29.48 | 8.90 | 3.46 | 8 | 95-99th percentile | 1.09 |
| DBX | 14309 | Female | 5.85 | Young | 49.9 | 5.41 | -2.73 | 2 | NA | 0.83 |
| BC | 1436 | Female | 7.52 | Young | 16 | 13.15 | -4.06 | 1 | NA | 0.94 |
| PS | 14404 | Female | 10.65 | Young | 10.21 | 12.94 | -1.44 | 1 | NA | 0.55 |
| YORK | 14417 | Male | 12.5 | Young | 3.18 | 10.41 | -0.21 | 3 | NA | 1 |
| DAC | 14458 | Male | 5.8 | Young | 10.89 | 11.31 | -6.23 | 4 | NA | 1.12 |
| DAC | 17434 | Female | 8.88 | Young | 10.89 | 11.31 | -3.15 | 4 | NA | 0.88 |
| BRI | 17495 | Female | 8.84 | Young | 15.88 | 10.98 | -2.75 | 7 | NA | 0.72 |
| GSD | 17497 | Female | 8.44 | Young | 35.15 | 8.26 | -1.44 | 1 | NA | 0.53 |
| DBX | 18202 | Male | 6.76 | Young | 49.9 | 5.41 | -1.82 | 2 | NA | 0.8 |
| POD | 18272 | Female | 9.58 | Young | 5.67 | 10.33 | -2.91 | 9 | NA | 1.07 |
| DAC | 18276 | Female | 9.56 | Young | 10.89 | 11.31 | -2.47 | 4 | NA | 1.06 |
| FCR | 18278 | Male | 8.06 | Young | 29.48 | 8.90 | -2.32 | 8 | NA | 0.44 |
| AS | 18301 | Female | 14.79 | Old | 26.08 | 10.44 | 4.11 | 1 | 95-99th percentile | 1.38 |
| FB | 18330 | Female | 8.33 | Young | 12.7 | 8.86 | -3.54 | 9 | NA | 0.59 |
| ROT | 18561 | Male | 8.48 | Young | 47.5 | 8.12 | -0.31 | 2 | NA | 0.72 |
| COC | 18718 | Female | 9.78 | Young | 14.06 | 12.88 | -1.97 | 8 | NA | 0.21 |
| BMD | 18736 | Male | 7.11 | Young | 44.23 | 6.88 | -1.97 | 2 | NA | 0.61 |
| ES | 18927 | Male | 7.52 | Young | 32.89 | 12.10 | -2.56 | 7 | NA | 0.72 |
| LAB | 18952 | Male | 9.48 | Young | 32.89 | 9.51 | -0.60 | 8 | NA | 1.09 |
| SCOT | 19146 | Female | 9.3 | Young | 9 | 12.19 | -2.89 | 3 | NA | 0.77 |
| SCOT | 19165 | Female | 12.51 | Old | 9 | 12.19 | 0.32 | 3 | <80th percentile | 1.35 |
| GD | 19865 | Female | 10.68 | Old | 66 | 5.41 | 3.53 | 2 | 90-95th percentile | 0.68 |
| LAB | 19869 | Male | 15.71 | Old | 32.89 | 9.51 | 5.63 | 8 | 90-95th percentile | 1.17 |
| GD | 19932 | Female | 10.94 | Old | 66 | 5.41 | 3.79 | 2 | 90-95th percentile | 0.79 |
| ROT | 20007 | Female | 13.97 | Old | 47.5 | 8.12 | 5.18 | 2 | 95-99th percentile | 0.75 |
| FB | 20018 | Female | 15.09 | Old | 12.7 | 8.86 | 3.22 | 9 | 95-99th percentile | 0.7 |
| GD | 20020 | Female | 10.01 | Old | 66 | 5.41 | 2.86 | 2 | 90-95th percentile | 0.82 |
| GD | 20023 | Male | 9.32 | Old | 66 | 5.41 | 2.17 | 2 | 90-95th percentile | 0.93 |
| GR | 20057 | Female | 15.73 | Old | 31.75 | 12.76 | 5.55 | 8 | 90-95th percentile | 0.89 |
| GD | 20089 | Male | 9.29 | Old | 66 | 5.41 | 2.14 | 2 | 90-95th percentile | 0.49 |
| GR | 20194 | Male | 15.19 | Old | 31.75 | 12.76 | 5.01 | 8 | 80-90th percentile | 0.98 |
| GD | 20239 | Female | 11.59 | Old | 66 | 5.41 | 4.44 | 2 | 90-95th percentile | 0.78 |
| FB | 20293 | Female | 15.4 | Old | 12.7 | 8.86 | 3.53 | 9 | 95-99th percentile | 0.9 |
| FB | 20294 | Female | 15.1 | Old | 12.7 | 8.86 | 3.23 | 9 | 95-99th percentile | 0.85 |
| BMD | 2047 | Female | 13.39 | Old | 44.23 | 6.88 | 4.31 | 2 | 90-95th percentile | 0.69 |
| BE | 20477 | Female | 9.83 | Young | 40.82 | 8.76 | 0.45 | 1 | NA | 0.9 |
| YORK | 20481 | Male | 18.25 | Old | 3.18 | 10.41 | 5.54 | 3 | 95-99th percentile | 0.92 |
| POD | 20487 | Male | 17.95 | Old | 5.67 | 10.33 | 5.46 | 9 | 95-99th percentile | 0.7 |
| BRI | 20488 | Female | 5.4 | Young | 15.88 | 10.98 | -6.19 | 7 | NA | 0.96 |
| BC | 20490 | Female | 16.03 | Old | 16 | 13.15 | 4.45 | 1 | 90-95th percentile | 0.77 |
| POD | 20492 | Male | 18.1 | Old | 5.67 | 10.33 | 5.61 | 9 | 95-99th percentile | 0.68 |
| ES | 20499 | Female | 16.01 | Old | 32.89 | 12.10 | 5.93 | 7 | 90-95th percentile | 1.03 |
| GR | 20501 | Male | 11.92 | Young | 31.75 | 12.76 | 1.74 | 8 | NA | 0.72 |
| FB | 20502 | Female | 6.64 | Young | 12.7 | 8.86 | -5.23 | 9 | NA | 0.57 |
| ES | 20503 | Male | 11.75 | Young | 32.89 | 12.10 | 1.67 | 7 | NA | 1.45 |
| FB | 20509 | Male | 14.11 | Old | 12.7 | 8.86 | 2.24 | 9 | 80-90th percentile | 0.65 |
| LAB | 20511 | Male | 16.02 | Old | 32.89 | 9.51 | 5.94 | 8 | 90-95th percentile | 1.1 |
| COC | 20512 | Female | 14.47 | Old | 14.06 | 12.88 | 2.72 | 8 | <80th percentile | 0.56 |
| SCOT | 20515 | Female | 12.23 | Old | 9 | 12.19 | 0.04 | 3 | <80th percentile | 0.84 |
| COC | 20516 | Female | 11.72 | Young | 14.06 | 12.88 | -0.03 | 8 | NA | 0.63 |
| ES | 20521 | Female | 7.49 | Young | 32.89 | 12.10 | -2.59 | 7 | NA | 0.95 |
| GR | 20525 | Female | 11.53 | Young | 31.75 | 12.76 | 1.35 | 8 | NA | 1.03 |
| LEO | 20528 | Female | 12.3 | Old | 58 | 6.96 | 4.44 | 2 | 80-90th percentile | 0.99 |
| BOX | 20532 | Female | 8.37 | Young | 32.89 | 8.59 | -1.71 | 2 | NA | 0.76 |
| YORK | 20533 | Female | 9.91 | Young | 3.18 | 10.41 | -2.80 | 3 | NA | 0.61 |
| YORK | 20536 | Female | 12 | Young | 3.18 | 10.41 | -0.71 | 3 | NA | 1.75 |
| FB | 20539 | Male | 13.44 | Old | 12.7 | 8.86 | 1.57 | 9 | 80-90th percentile | 0.79 |
| GD | 20540 | Male | 5.79 | Young | 66 | 5.41 | -1.36 | 2 | NA | 1.15 |
| BRI | 20638 | Male | 15.28 | Old | 15.88 | 10.98 | 3.69 | 7 | 80-90th percentile | 0.86 |
| BMD | 20649 | Male | 13.16 | Old | 44.23 | 6.88 | 4.08 | 2 | 90-95th percentile | 1.13 |
| BMD | 20651 | Female | 8.2 | Young | 44.23 | 6.88 | -0.88 | 2 | NA | 0.84 |
| BMD | 20652 | Female | 7.8 | Young | 44.23 | 6.88 | -1.28 | 2 | NA | 0.8 |
| BMD | 20654 | Female | 7.4 | Young | 44.23 | 6.88 | -1.68 | 2 | NA | 1.03 |
| BMD | 20655 | Female | 7.5 | Young | 44.23 | 6.88 | -1.58 | 2 | NA | 0.69 |
| BMD | 20656 | Male | 7.12 | Young | 44.23 | 6.88 | -1.96 | 2 | NA | 1.22 |
| BMD | 20657 | Female | 6.5 | Young | 44.23 | 6.88 | -2.58 | 2 | NA | 0.54 |
| BMD | 20659 | Female | 13.98 | Old | 44.23 | 6.88 | 4.90 | 2 | 90-95th percentile | 1.43 |
| ROT | 20731 | Male | 13.39 | Old | 47.5 | 8.12 | 4.60 | 2 | 90-95th percentile | 0.84 |
| POD | 20753 | Female | 17.78 | Old | 5.67 | 10.33 | 5.29 | 9 | 90-95th percentile | 1.1 |
| POD | 20754 | Female | 18.15 | Old | 5.67 | 10.33 | 5.66 | 9 | 95-99th percentile | 0.5 |
| COC | 2234 | Male | 15.25 | Old | 14.06 | 12.88 | 3.50 | 8 | 80-90th percentile | 0.94 |
| COC | 2314 | Male | 9.78 | Young | 14.06 | 12.88 | -1.97 | 8 | NA | 0.57 |
| LAB | 232 | Female | 8.83 | Young | 32.89 | 9.51 | -1.25 | 8 | NA | 1.11 |
| DBX | 2574 | Male | 10.76 | Old | 49.9 | 5.41 | 2.18 | 2 | 80-90th percentile | 0.65 |
| GSMD | 259 | Female | 13.41 | Old | 57.83 | 9.34 | 5.53 | 2 | 95-99th percentile | 0.96 |
| BMD | 2804 | Male | 12.64 | Old | 44.23 | 6.88 | 3.56 | 2 | 80-90th percentile | 0.47 |
| GSMD | 2857 | Female | 11.24 | Old | 57.83 | 9.34 | 3.36 | 2 | 90-95th percentile | 0.65 |
| LAB | 3069 | Female | 16.1 | Old | 32.89 | 9.51 | 6.02 | 8 | 90-95th percentile | 0.95 |
| LAB | 3119 | Female | 16 | Old | 32.89 | 9.51 | 5.92 | 8 | 90-95th percentile | 0.98 |
| DBX | 3171 | Male | 10.36 | Old | 49.9 | 5.41 | 1.78 | 2 | 80-90th percentile | 0.78 |
| PS | 3359 | Female | 10.24 | Young | 10.21 | 12.94 | -1.85 | 1 | NA | 0.78 |
| PS | 3367 | Female | 10.79 | Young | 10.21 | 12.94 | -1.30 | 1 | NA | 0.67 |
| PS | 3371 | Male | 10.32 | Young | 10.21 | 12.94 | -1.77 | 1 | NA | 0.47 |
| PS | 3385 | Female | 10.3 | Young | 10.21 | 12.94 | -1.79 | 1 | NA | 0.62 |
| ES | 3562 | Male | 15.65 | Old | 32.89 | 12.10 | 5.57 | 7 | 80-90th percentile | 0.99 |
| PS | 3633 | Male | 16.07 | Old | 10.21 | 12.94 | 3.98 | 1 | 80-90th percentile | 0.87 |
| PS | 3634 | Female | 9.95 | Young | 10.21 | 12.94 | -2.14 | 1 | NA | 0.65 |
| LEO | 3662 | Female | 6.21 | Young | 58 | 6.96 | -1.65 | 2 | NA | 0.76 |
| BMD | 3798 | Male | 7.67 | Young | 44.23 | 6.88 | -1.41 | 2 | NA | 0.68 |
| YORK | 3859 | Female | 17.35 | Old | 3.18 | 10.41 | 4.64 | 3 | 90-95th percentile | 1.38 |
| PS | 4072 | Female | 15.04 | Old | 10.21 | 12.94 | 2.95 | 1 | <80th percentile | 0.99 |
| COC | 4094 | Male | 15.48 | Old | 14.06 | 12.88 | 3.73 | 8 | 90-95th percentile | 0.79 |
| FB | 4126 | Male | 5.49 | Young | 12.7 | 8.86 | -6.38 | 9 | NA | 1.16 |
| GSMD | 4247 | Female | 6.56 | Young | 57.83 | 9.34 | -1.32 | 2 | NA | 1.06 |
| PS | 4540 | Female | 9.9 | Young | 10.21 | 12.94 | -2.19 | 1 | NA | 0.75 |
| LAB | 4639 | Female | 9.26 | Young | 32.89 | 9.51 | -0.82 | 8 | NA | 0.71 |
| AS | 4681 | Male | 9.73 | Young | 26.08 | 10.44 | -0.95 | 1 | NA | 1.11 |
| AS | 4766 | Male | 6.44 | Young | 26.08 | 10.44 | -4.24 | 1 | NA | 1.05 |
| LAB | 4819 | Female | 8.5 | Young | 32.89 | 9.51 | -1.58 | 8 | NA | 1 |
| BOX | 5141 | Female | 15.36 | Old | 32.89 | 8.59 | 5.28 | 2 | 95-99th percentile | 0.76 |
| YORK | 5502 | Female | 9.7 | Young | 3.18 | 10.41 | -3.01 | 3 | NA | 0.97 |
| LEO | 5523 | Female | 14.59 | Old | 58 | 6.96 | 6.73 | 2 | 95-99th percentile | 0.67 |
| PS | 5572 | Male | 16.44 | Old | 10.21 | 12.94 | 4.35 | 1 | 80-90th percentile | 0.74 |
| PS | 5583 | Female | 15.3 | Old | 10.21 | 12.94 | 3.21 | 1 | <80th percentile | 0.94 |
| BE | 5699 | Male | 14.07 | Old | 40.82 | 8.76 | 4.69 | 1 | 80-90th percentile | 0.41 |
| POD | 5989 | Female | 12.42 | Young | 5.67 | 10.33 | -0.07 | 9 | NA | 0.93 |
| BC | 6093 | Male | 15.99 | Old | 16 | 13.15 | 4.41 | 1 | 90-95th percentile | 0.83 |
| LEO | 6112 | Male | 7.59 | Young | 58 | 6.96 | -0.27 | 2 | NA | 0.88 |
| LEO | 6127 | Female | 7.18 | Young | 58 | 6.96 | -0.68 | 2 | NA | 0.91 |
| LEO | 6133 | Male | 7.46 | Young | 58 | 6.96 | -0.40 | 2 | NA | 0.84 |
| LEO | 6136 | Female | 7.78 | Young | 58 | 6.96 | -0.08 | 2 | NA | 0.88 |
| LEO | 6141 | Female | 12.72 | Old | 58 | 6.96 | 4.86 | 2 | 90-95th percentile | 1.13 |
| LEO | 6142 | Female | 6.73 | Young | 58 | 6.96 | -1.13 | 2 | NA | 0.56 |
| LEO | 6149 | Female | 7.5 | Young | 58 | 6.96 | -0.36 | 2 | NA | 1.27 |
| GR | 6180 | Male | 11.06 | Young | 31.75 | 12.76 | 0.88 | 8 | NA | 0.61 |
| AS | 6386 | Male | 15.15 | Old | 26.08 | 10.44 | 4.47 | 1 | 95-99th percentile | 0.35 |
| COC | 651 | Male | 14.9 | Old | 14.06 | 12.88 | 3.15 | 8 | 80-90th percentile | 0.73 |
| AS | 6522 | Female | 5.95 | Young | 26.08 | 10.44 | -4.73 | 1 | NA | 0.3 |
| GD | 6529 | Female | 4.69 | Young | 66 | 5.41 | -2.46 | 2 | NA | 0.92 |
| ROT | 6660 | Female | 7.82 | Young | 47.5 | 8.12 | -0.97 | 2 | NA | 1.28 |
| FCR | 6690 | Male | 15 | Old | 29.48 | 8.90 | 4.62 | 8 | >=99th percentile | 1.31 |
| DBX | 6832 | Female | 12.28 | Old | 49.9 | 5.41 | 3.70 | 2 | 95-99th percentile | 0.81 |
| FCR | 6847 | Female | 8.55 | Young | 29.48 | 8.90 | -1.83 | 8 | NA | 0.38 |
| FCR | 6859 | Female | 15 | Old | 29.48 | 8.90 | 4.62 | 8 | >=99th percentile | 0.46 |
| FCR | 6863 | Male | 14.66 | Old | 29.48 | 8.90 | 4.28 | 8 | >=99th percentile | 0.62 |
| FCR | 6893 | Female | 8.36 | Young | 29.48 | 8.90 | -2.02 | 8 | NA | 0.67 |
| SCOT | 6922 | Female | 10.27 | Young | 9 | 12.19 | -1.92 | 3 | NA | 0.83 |
| BC | 6979 | Female | 15.66 | Old | 16 | 13.15 | 4.08 | 1 | 80-90th percentile | 1.15 |
| FCR | 7012 | Male | 8.49 | Young | 29.48 | 8.90 | -1.89 | 8 | NA | 0.85 |
| FCR | 7018 | Female | 13.51 | Old | 29.48 | 8.90 | 3.13 | 8 | 90-95th percentile | 0.86 |
| FCR | 7019 | Female | 14.24 | Old | 29.48 | 8.90 | 3.86 | 8 | >=99th percentile | 0.99 |
| BOX | 7053 | Male | 12.86 | Old | 32.89 | 8.59 | 2.78 | 2 | <80th percentile | 1.05 |
| SCOT | 7073 | Male | 13.88 | Old | 9 | 12.19 | 1.69 | 3 | <80th percentile | 0.84 |
| BC | 7080 | Male | 7.44 | Young | 16 | 13.15 | -4.14 | 1 | NA | 0.81 |
| FCR | 7118 | Female | 8.92 | Young | 29.48 | 8.90 | -1.46 | 8 | NA | 1.08 |
| FCR | 7132 | Female | 12.62 | Old | 29.48 | 8.90 | 2.24 | 8 | 90-95th percentile | 1.01 |
| DBX | 7178 | Male | 10.33 | Old | 49.9 | 5.41 | 1.75 | 2 | 80-90th percentile | 1.08 |
| BE | 7259 | Female | 15.22 | Old | 40.82 | 8.76 | 5.84 | 1 | 95-99th percentile | 0.97 |
| BMD | 7288 | Male | 7.18 | Young | 44.23 | 6.88 | -1.90 | 2 | NA | 1.07 |
| SCOT | 7400 | Female | 13.43 | Old | 9 | 12.19 | 1.24 | 3 | <80th percentile | 0.6 |
| PS | 7407 | Female | 9.76 | Young | 10.21 | 12.94 | -2.33 | 1 | NA | 0.79 |
| COC | 7495 | Male | 16.56 | Old | 14.06 | 12.88 | 4.81 | 8 | 95-99th percentile | 0.69 |
| GSMD | 7686 | Female | 7.82 | Young | 57.83 | 9.34 | -0.06 | 2 | NA | 1.32 |
| BMD | 7707 | Female | 13.33 | Old | 44.23 | 6.88 | 4.25 | 2 | 90-95th percentile | 0.82 |
| LEO | 7753 | Female | 6.99 | Young | 58 | 6.96 | -0.87 | 2 | NA | 1.12 |
| ROT | 7796 | Female | 8.01 | Young | 47.5 | 8.12 | -0.78 | 2 | NA | 0.79 |
| GSMD | 7807 | Female | 11.23 | Old | 57.83 | 9.34 | 3.35 | 2 | 90-95th percentile | 0.83 |
| BOX | 7824 | Male | 13.59 | Old | 32.89 | 8.59 | 3.51 | 2 | 80-90th percentile | 0.78 |
| POD | 7839 | Male | 16.85 | Old | 5.67 | 10.33 | 4.36 | 9 | 80-90th percentile | 0.47 |
| POD | 7855 | Female | 12.18 | Young | 5.67 | 10.33 | -0.31 | 9 | NA | 0.94 |
| BMD | 7912 | Female | 12.85 | Old | 44.23 | 6.88 | 3.77 | 2 | 80-90th percentile | 0.84 |
| ES | 7917 | Female | 10.24 | Young | 32.89 | 12.10 | 0.16 | 7 | NA | 0.72 |
| DBX | 7986 | Female | 11.1 | Old | 49.9 | 5.41 | 2.52 | 2 | 90-95th percentile | 0.37 |
| BE | 8002 | Male | 13.82 | Old | 40.82 | 8.76 | 4.44 | 1 | 80-90th percentile | 0.85 |
| DBX | 8263 | Female | 6.85 | Young | 49.9 | 5.41 | -1.73 | 2 | NA | 0.66 |
| ES | 8265 | Male | 11.43 | Young | 32.89 | 12.10 | 1.35 | 7 | NA | 0.92 |
| BOX | 8488 | Female | 17.47 | Old | 32.89 | 8.59 | 7.39 | 2 | >=99th percentile | 0.95 |
| ROT | 8584 | Male | 8.34 | Young | 47.5 | 8.12 | -0.45 | 2 | NA | 1.08 |
| BE | 8708 | Female | 8.68 | Young | 40.82 | 8.76 | -0.70 | 1 | NA | 0.68 |
| ES | 8845 | Male | 14.62 | Old | 32.89 | 12.10 | 4.54 | 7 | <80th percentile | 1.04 |
| FB | 8930 | Male | 14.45 | Old | 12.7 | 8.86 | 2.58 | 9 | 90-95th percentile | 0.71 |
| GR | 8967 | Female | 15.35 | Old | 31.75 | 12.76 | 5.17 | 8 | 90-95th percentile | 0.65 |
| BOX | 9040 | Male | 14.53 | Old | 32.89 | 8.59 | 4.45 | 2 | 95-99th percentile | 0.93 |
| ROT | 9056 | Female | 13.83 | Old | 47.5 | 8.12 | 5.04 | 2 | 90-95th percentile | 1.09 |
| YORK | 9087 | Female | 22.16 | Old | 3.18 | 10.41 | 9.45 | 3 | >=99th percentile | 1.06 |
| SCOT | 9136 | Male | 10.54 | Young | 9 | 12.19 | -1.65 | 3 | NA | 1.67 |
| LAB | 9142 | Female | 16.04 | Old | 32.89 | 9.51 | 5.96 | 8 | 90-95th percentile | 0.93 |
| AS | 9144 | Male | 7.12 | Young | 26.08 | 10.44 | -3.56 | 1 | NA | 0.93 |
| ROT | 9177 | Male | 13.3 | Old | 47.5 | 8.12 | 4.51 | 2 | 90-95th percentile | 0.56 |
| ROT | 9189 | Male | 13.28 | Old | 47.5 | 8.12 | 4.49 | 2 | 90-95th percentile | 1.35 |
| YORK | 9201 | Female | 16.96 | Old | 3.18 | 10.41 | 4.25 | 3 | 90-95th percentile | 0.77 |
| SCOT | 9254 | Female | 9.81 | Young | 9 | 12.19 | -2.38 | 3 | NA | 0.81 |
| SCOT | 9284 | Male | 12.27 | Old | 9 | 12.19 | 0.08 | 3 | <80th percentile | 0.88 |
| ES | 9303 | Female | 10.39 | Young | 32.89 | 12.10 | 0.31 | 7 | NA | 0.96 |
| FB | 9325 | Female | 9.79 | Young | 12.7 | 8.86 | -2.08 | 9 | NA | 0.98 |
| GD | 9393 | Female | 5.35 | Young | 66 | 5.41 | -1.80 | 2 | NA | 0.84 |
| SCOT | 9471 | Male | 14.59 | Old | 9 | 12.19 | 2.40 | 3 | 80-90th percentile | 1.2 |
| ES | 9501 | Male | 10.78 | Young | 32.89 | 12.10 | 0.70 | 7 | NA | 0.98 |
| SCOT | 9547 | Female | 10.25 | Young | 9 | 12.19 | -1.94 | 3 | NA | 0.86 |
| FB | 9591 | Male | 6.65 | Young | 12.7 | 8.86 | -5.22 | 9 | NA | 1.19 |
| DAC | 9613 | Female | 15.72 | Old | 10.89 | 11.31 | 3.69 | 4 | 80-90th percentile | 1.11 |
| GR | 9653 | Male | 16.59 | Old | 31.75 | 12.76 | 6.41 | 8 | >=99th percentile | 0.19 |
| YORK | 9707 | Female | 16.42 | Old | 3.18 | 10.41 | 3.71 | 3 | 80-90th percentile | 0.66 |
| GD | 9735 | Female | 5.53 | Young | 66 | 5.41 | -1.62 | 2 | NA | 1.12 |
| ES | 9858 | Male | 14.73 | Old | 32.89 | 12.10 | 4.65 | 7 | <80th percentile | 0.96 |
| GSD | 9860 | Female | 8.97 | Young | 35.15 | 8.26 | -0.91 | 1 | NA | 0.84 |
| GR | 9937 | Male | 16 | Old | 31.75 | 12.76 | 5.82 | 8 | 95-99th percentile | 0.83 |
| FB | 12589 | Male | 14.94 | Old | 12.7 | 8.86 | 3.07 | 9 | 95-99th percentile | 1.25 |
| COC | 10468 | Female | 9.99 | Young | 14.06 | 12.88 | -1.76 | 8 | NA | 1.06 |
| DAC | 11880 | Male | 15.22 | Old | 10.89 | 11.31 | 3.19 | 4 | 80-90th percentile | 0.47 |
| GD | 7385 | Male | 5.35 | Young | 66 | 5.41 | -1.80 | 2 | NA | 0.95 |
| ES | 4635 | Female | 9.19 | Young | 32.89 | 12.10 | -0.89 | 7 | NA | 1.27 |
| GR | 7723 | Female | 11.99 | Young | 31.75 | 12.76 | 1.81 | 8 | NA | 1.19 |
| LAB | 8271 | Male | 8.35 | Young | 32.89 | 9.51 | -1.73 | 8 | NA | 0.78 |
| LEO | X159 | Male | 14 | Old | 58 | 6.96 | 6.14 | 2 | 95-99th percentile | 0.69 |
| LEO | X163 | Female | 14 | Old | 58 | 6.96 | 6.14 | 2 | 95-99th percentile | 0.76 |
| GD | 2800 | Male | 9.62 | Old | 66 | 5.41 | 2.47 | 2 | 90-95th percentile | 0.79 |
| BRI | 4799 | Female | 11.18 | Young | 15.88 | 10.98 | -0.41 | 7 | NA | 0.73 |
| BE | 5825 | Male | 6.19 | Young | 40.82 | 8.76 | -3.19 | 1 | NA | 0.68 |
| COC | 65 | Male | 14.74 | Old | 14.06 | 12.88 | 2.99 | 8 | 80-90th percentile | 1.24 |
| DAC | 103421 | Female | 9.81 | Young | 10.89 | 11.31 | -2.22 | 4 | NA | 0.55 |
| BE | 10594 | Male | 13.62 | Old | 40.82 | 8.76 | 4.24 | 1 | 80-90th percentile | 0.62 |
| AI | 114508 | Female | 8.96 | Young | 45 | 10.78 | -0.05 | 5 | NA | 0.88 |
| WSSD | 11476 | Female | 9.5 | Young | 32.5 | 8.26 | -0.62 | NA | NA | 0.63 |
| WHI | 11758 | Male | 16.42 | Old | 14.74 | 13.67 | 4.73 | 10 | 95-99th percentile | 0.58 |
| WSSD | 132119 | Female | 14.19 | Old | 32.5 | 8.26 | 4.07 | NA | NA | 0.72 |
| DAC | 147611 | Female | 6.34 | Young | 10.89 | 11.31 | -5.69 | 4 | NA | 0.84 |
| DAC | 15594 | Male | 16.93 | Old | 10.89 | 11.31 | 4.90 | 4 | 95-99th percentile | 0.42 |
| WHI | 15750 | Male | 11.31 | Young | 14.74 | 13.67 | -0.38 | 10 | NA | 0.4 |
| WHI | 15752 | Female | 10.4 | Young | 14.74 | 13.67 | -1.29 | 10 | NA | 0.55 |
| WSSD | 16077 | Male | 10.44 | Young | 32.5 | 8.26 | 0.32 | NA | NA | 0.44 |
| WHI | 16818 | Female | 11.78 | Young | 14.74 | 13.67 | 0.09 | 10 | NA | 0.37 |
| WSSD | 17557 | Male | 13.76 | Old | 32.5 | 8.26 | 3.64 | NA | NA | 0.39 |
| WSSD | 17837 | Female | 14.21 | Old | 32.5 | 8.26 | 4.09 | NA | NA | 0.56 |
| WSSD | 21059 | Female | 14.45 | Old | 32.5 | 8.26 | 4.33 | NA | NA | 0.49 |
| BE | 2124 | Male | 16.12 | Old | 40.82 | 8.76 | 6.74 | 1 | 95-99th percentile | 0.68 |
| WHI | 21300 | Female | 16.13 | Old | 14.74 | 13.67 | 4.44 | 10 | 90-95th percentile | 0.46 |
| WSSD | 23363 | Female | 14.19 | Old | 32.5 | 8.26 | 4.07 | NA | NA | 0.49 |
| WSSD | 26122 | Male | 14.34 | Old | 32.5 | 8.26 | 4.22 | NA | NA | 1.34 |
| BOX | 2717 | Female | 16.04 | Old | 32.89 | 8.59 | 5.96 | 2 | >=99th percentile | 0.58 |
| PS | 28586 | Male | 17 | Old | 10.21 | 12.94 | 4.91 | 1 | 90-95th percentile | 0.81 |
| WSSD | 29109 | Female | 14.04 | Old | 32.5 | 8.26 | 3.92 | NA | NA | 0.43 |
| WSSD | 29941 | Female | 9.6 | Young | 32.5 | 8.26 | -0.52 | NA | NA | 0.59 |
| WSSD | 31150 | Female | 9.82 | Young | 32.5 | 8.26 | -0.30 | NA | NA | 0.43 |
| WSSD | 31251 | Male | 9.61 | Young | 32.5 | 8.26 | -0.51 | NA | NA | 0.47 |
| WSSD | 31613 | Male | 9.96 | Young | 32.5 | 8.26 | -0.16 | NA | NA | 0.4 |
| WSSD | 31698 | Female | 10.21 | Young | 32.5 | 8.26 | 0.09 | NA | NA | 0.79 |
| WSSD | 31713 | Male | 13.76 | Old | 32.5 | 8.26 | 3.64 | NA | NA | 0.8 |
| WSSD | 32451 | Female | 10.27 | Young | 32.5 | 8.26 | 0.15 | NA | NA | 0.72 |
| WHI | 34364 | Female | 16.25 | Old | 14.74 | 13.67 | 4.56 | 10 | 90-95th percentile | 0.38 |
| AS | 36494 | Male | 14.48 | Old | 26.08 | 10.44 | 3.80 | 1 | 95-99th percentile | 0.49 |
| SCOT | 4023 | Female | 16.05 | Old | 9 | 12.19 | 3.86 | 3 | 95-99th percentile | 0.28 |
| WSSD | 40766 | Female | 9.77 | Young | 32.5 | 8.26 | -0.35 | NA | NA | 0.71 |
| WHI | 4414 | Male | 16.1 | Old | 14.74 | 13.67 | 4.41 | 10 | 90-95th percentile | 0.35 |
| WHI | 4512 | Male | 11.42 | Young | 14.74 | 13.67 | -0.27 | 10 | NA | 0.52 |
| WHI | 46404 | Female | 16.04 | Old | 14.74 | 13.67 | 4.35 | 10 | 90-95th percentile | 0.51 |
| AI | 47305 | Male | 12.96 | Old | 45 | 10.78 | 3.95 | 5 | <80th percentile | 0.42 |
| DAC | 50577 | Male | 9.1 | Young | 10.89 | 11.31 | -2.93 | 4 | NA | 0.58 |
| ES | 51184 | Female | 15.15 | Old | 32.89 | 12.10 | 5.07 | 7 | 80-90th percentile | 0.38 |
| AI | 53627 | Female | 13.38 | Old | 45 | 10.78 | 4.37 | 5 | <80th percentile | 0.85 |
| WHI | 5464 | Female | 9.83 | Young | 14.74 | 13.67 | -1.86 | 10 | NA | 0.57 |
| WHI | 5465 | Male | 11.28 | Young | 14.74 | 13.67 | -0.41 | 10 | NA | 0.69 |
| AI | 55898 | Male | 13.95 | Old | 45 | 10.78 | 4.94 | 5 | 80-90th percentile | 0.69 |
| WSSD | 56216 | Female | 14.5 | Old | 32.5 | 8.26 | 4.38 | NA | NA | 1.06 |
| WSSD | 57213 | Male | 14.85 | Old | 32.5 | 8.26 | 4.73 | NA | NA | 0.58 |
| ES | 57369 | Male | 15.27 | Old | 32.89 | 12.10 | 5.19 | 7 | 80-90th percentile | 1.15 |
| WSSD | 60187 | Female | 10.12 | Young | 32.5 | 8.26 | 0.00 | NA | NA | 0.68 |
| WHI | 6397 | Female | 16.19 | Old | 14.74 | 13.67 | 4.50 | 10 | 90-95th percentile | 0.41 |
| WHI | 6398 | Female | 16.99 | Old | 14.74 | 13.67 | 5.30 | 10 | 95-99th percentile | 0.57 |
| WHI | 6558 | Female | 11.39 | Young | 14.74 | 13.67 | -0.30 | 10 | NA | 0.3 |
| WHI | 6754 | Female | 16.25 | Old | 14.74 | 13.67 | 4.56 | 10 | 90-95th percentile | 0.34 |
| SCOT | 7981 | Male | 9.5 | Young | 9 | 12.19 | -2.69 | 3 | NA | 0.52 |
| GSD | 8234 | Female | 15.4 | Old | 35.15 | 8.26 | 5.52 | 1 | 90-95th percentile | 0.5 |
| WHI | 8843 | Male | 10 | Young | 14.74 | 13.67 | -1.69 | 10 | NA | 0.62 |
| BE | 9234 | Male | 13.94 | Old | 40.82 | 8.76 | 4.56 | 1 | 80-90th percentile | 0.51 |
| AI | 96436 | Male | 9.04 | Young | 45 | 10.78 | 0.03 | 5 | NA | 0.42 |
| DAC | 96487 | Female | 6.14 | Young | 10.89 | 11.31 | -5.89 | 4 | NA | 0.58 |
| GSD | 6040 | Female | 13.98 | Old | 35.15 | 8.26 | 4.10 | 1 | 80-90th percentile | 0.61 |
| BE | 1763 | Female | 14.22 | Old | 40.82 | 8.76 | 4.84 | 1 | 80-90th percentile | 0.49 |
| BE | 2209 | Female | 5.61 | Young | 40.82 | 8.76 | -3.77 | 1 | NA | 0.34 |
| FB | 10896 | Male | 8.62 | Young | 12.7 | 8.86 | -3.25 | 9 | NA | 0.37 |
| DBX | 2623 | Female | 6.52 | Young | 49.9 | 5.41 | -2.06 | 2 | NA | 0.48 |
| ES | 7549 | Female | 14.96 | Old | 32.89 | 12.10 | 4.88 | 7 | 80-90th percentile | 1.08 |
| POD | 108 | Female | 17.03 | Old | 5.67 | 10.33 | 4.54 | 9 | 80-90th percentile | 0.97 |
| DAC | 5525 | Female | 14.02 | Old | 10.89 | 11.31 | 1.99 | 4 | <80th percentile | 0.73 |
| GR | 775 | Male | 14.85 | Old | 31.75 | 12.76 | 4.67 | 8 | 80-90th percentile | 1 |
| DAC | 9479 | Male | 13.89 | Old | 10.89 | 11.31 | 1.86 | 4 | <80th percentile | 1.22 |
| GD | 7539 | Male | 5.86 | Young | 66 | 5.41 | -1.29 | 2 | NA | 0.67 |
| ES | 8959 | Male | 16.13 | Old | 32.89 | 12.10 | 6.05 | 7 | 90-95th percentile | 0.79 |
| BE | 171 | Male | 6.14 | Young | 40.82 | 8.76 | -3.24 | 1 | NA | 0.41 |
| GD | 6445 | Female | 5.45 | Young | 66 | 5.41 | -1.70 | 2 | NA | 0.44 |
| POD | 40481 | Male | 16 | Old | 5.67 | 10.33 | 3.51 | 9 | 80-90th percentile | 1.55 |
| DAC | 10908 | Female | 14.88 | Old | 10.89 | 11.31 | 2.85 | 4 | <80th percentile | 0.85 |
| AI | 13331 | Male | 13.93 | Old | 45 | 10.78 | 4.92 | 5 | 80-90th percentile | 1.74 |
| WHI | 15986 | Female | 10.64 | Young | 14.74 | 13.67 | -1.05 | 10 | NA | 0.87 |
| DAC | 7353 | Female | 11.43 | Young | 10.89 | 11.31 | -0.60 | 4 | NA | 0.87 |
| BRI | 5993 | Female | 8.4 | Young | 15.88 | 10.98 | -3.19 | 7 | NA | 0.45 |
| SCOT | 6408 | Female | 15.2 | Old | 9 | 12.19 | 3.01 | 3 | 90-95th percentile | 0.71 |
| BE | 4009 | Male | 9.64 | Young | 40.82 | 8.76 | 0.26 | 1 | NA | 0.76 |
| BE | 7057 | Male | 9.21 | Young | 40.82 | 8.76 | -0.17 | 1 | NA | 0.26 |
| DAC | 931 | Male | 4.73 | Young | 10.89 | 11.31 | -7.30 | 4 | NA | 0.77 |
| SCOT | 4211 | Female | 14.4 | Old | 9 | 12.19 | 2.21 | 3 | 80-90th percentile | 1.04 |
| COC | 8311 | Female | 10.11 | Young | 14.06 | 12.88 | -1.64 | 8 | NA | 0.7 |
| FCR | 7058 | Male | 7.64 | Young | 29.48 | 8.90 | -2.74 | 8 | NA | 0.96 |
| BRI | 823 | Female | 5.61 | Young | 15.88 | 10.98 | -5.98 | 7 | NA | 0.26 |
| ES | 3506 | Female | 10.33 | Young | 32.89 | 12.10 | 0.25 | 7 | NA | 0.42 |
| COC | 7687 | Female | 14.9 | Old | 14.06 | 12.88 | 3.15 | 8 | 80-90th percentile | 0.86 |
| YORK | 1051 | Male | 12.27 | Young | 3.18 | 10.41 | -0.44 | 3 | NA | 0.84 |
| GD | 2846 | Male | 10.39 | Old | 66 | 5.41 | 3.24 | 2 | 90-95th percentile | 0.45 |
| COC | 3924 | Female | 15.08 | Old | 14.06 | 12.88 | 3.33 | 8 | 80-90th percentile | 0.54 |
| GR | 2194 | Female | 11.98 | Young | 31.75 | 12.76 | 1.80 | 8 | NA | 0.7 |
| SCOT | 5281 | Female | 10.45 | Young | 9 | 12.19 | -1.74 | 3 | NA | 0.84 |
| GR | 6438 | Male | 11.6 | Young | 31.75 | 12.76 | 1.42 | 8 | NA | 0.48 |
| GD | 7295 | Female | 10.62 | Old | 66 | 5.41 | 3.47 | 2 | 90-95th percentile | 0.48 |
| COC | 3498 | Male | 8.62 | Young | 14.06 | 12.88 | -3.13 | 8 | NA | 0.74 |
| BE | 376 | Female | 14.81 | Old | 40.82 | 8.76 | 5.43 | 1 | 90-95th percentile | 1.27 |
| GD | 627 | Female | 11.89 | Old | 66 | 5.41 | 4.74 | 2 | 95-99th percentile | 0.8 |
| DAC | 9572 | Male | 14.9 | Old | 10.89 | 11.31 | 2.87 | 4 | <80th percentile | 0.67 |
| DAC | 1717 | Female | 15.25 | Old | 10.89 | 11.31 | 3.22 | 4 | 80-90th percentile | 0.6 |
| YORK | 313 | Male | 19.02 | Old | 3.18 | 10.41 | 6.31 | 3 | >=99th percentile | 1.01 |
| BOX | 2107 | Male | 8.51 | Young | 32.89 | 8.59 | -1.57 | 2 | NA | 0.76 |
| AI | 5097 | Female | 14.59 | Old | 45 | 10.78 | 5.58 | 5 | 90-95th percentile | 0.36 |
| BOX | 5931 | Female | 9.32 | Young | 32.89 | 8.59 | -0.76 | 2 | NA | 0.7 |
| GR | 6287 | Female | 17.44 | Old | 31.75 | 12.76 | 7.26 | 8 | >=99th percentile | 0.65 |
| COC | 2517 | Male | 8.11 | Young | 14.06 | 12.88 | -3.64 | 8 | NA | 0.73 |
| ROT | 10184 | Female | 8.898630137 | Young | 47.5 | 8.12 | 0.11 | 2 | NA | 0.973457387 |
| LAB | 12915 | Male | 14.4739726 | Old | 32.89 | 9.51 | 4.39 | 8 | 80-90th percentile | 0.950001495 |
| POD | 134447 | Female | 11.38 | Young | 5.67 | 10.33 | -1.11 | 9 | NA | 0.785844054 |
| PS | 142953 | Female | 15.85479452 | Old | 10.21 | 12.94 | 3.77 | 1 | 80-90th percentile | 0.860069162 |
| GSD | 20224 | Female | 13.30958904 | Old | 35.15 | 8.26 | 3.43 | 1 | <80th percentile | 0.977297149 |
| GSD | 20432 | Female | 13.65205479 | Old | 35.15 | 8.26 | 3.77 | 1 | 80-90th percentile | 0.986084453 |
| GSD | 20752 | Female | 13.7 | Old | 35.15 | 8.26 | 3.82 | 1 | 80-90th percentile | 0.971054272 |
| BOX | 215876 | Female | 13.97534247 | Old | 32.89 | 8.59 | 3.89 | 2 | 90-95th percentile | 0.982223489 |
| BRI | 250269800928940 | Female | 15.12876712 | Old | 15.88 | 10.98 | 3.54 | 7 | 80-90th percentile | 0.990248503 |
| GSD | 2EVR960 | Male | 14.33972603 | Old | 35.15 | 8.26 | 4.46 | 1 | 80-90th percentile | 0.900899227 |
| LAB | 4619 | Male | 15.07123288 | Old | 32.89 | 9.51 | 4.99 | 8 | 80-90th percentile | 0.972489894 |
| BMD | 7533 | Male | 12.15 | Old | 44.23 | 6.88 | 3.07 | 2 | 80-90th percentile | 0.993250919 |
| POD | 8044 | Female | 16.71506849 | Old | 5.67 | 10.33 | 4.23 | 9 | 80-90th percentile | 0.838400203 |
| BC | 8812 | Female | 9.739726027 | Young | 16 | 13.15 | -1.84 | 1 | NA | 0.973119989 |
| GSD | UMB380 | Male | 12.8109589 | Old | 35.15 | 8.26 | 2.93 | 1 | <80th percentile | 1.006901934 |
| GR | 10295 | Female | 14.89589041 | Old | 31.75 | 12.76 | 4.71 | 8 | 80-90th percentile | 1.42946742 |
| BOX | 10774 | Male | 6.82 | Young | 32.89 | 8.59 | -3.26 | 2 | NA | 1.935156613 |
| BOX | 11086 | Female | 13.39452055 | Old | 32.89 | 8.59 | 3.31 | 2 | 80-90th percentile | 1.115096362 |
| BRI | 12592 | Female | 16.2630137 | Old | 15.88 | 10.98 | 4.68 | 7 | 80-90th percentile | 1.436963206 |
| COC | 12779 | Male | 20.30684932 | Old | 14.06 | 12.88 | 8.56 | 8 | >=99th percentile | 1.95957591 |
| GR | 9419 | Female | 10.15 | Young | 31.75 | 12.76 | -0.03 | 8 | NA | 1.325642932 |
| FB | 20055 | Female | 13.50410959 | Old | 12.7 | 8.86 | 1.64 | 9 | 80-90th percentile | 1.426856024 |
| GR | 20099 | Male | 14.54246575 | Old | 31.75 | 12.76 | 4.36 | 8 | <80th percentile | 1.377635151 |
| FCR | 20154 | Female | 13.1 | Old | 29.48 | 8.90 | 2.72 | 8 | 90-95th percentile | 1.544106136 |
| WHI | 20181 | Male | 15.51506849 | Old | 14.74 | 13.67 | 3.83 | 10 | 80-90th percentile | 1.202184274 |
| PS | 20265 | Male | 14.87671233 | Old | 10.21 | 12.94 | 2.79 | 1 | <80th percentile | 1.678385811 |
| LEO | 20428 | Female | 13.3 | Old | 58 | 6.96 | 5.44 | 2 | 95-99th percentile | 0.782211619 |
| LEO | 20455 | Male | 11.7 | Old | 58 | 6.96 | 3.84 | 2 | 80-90th percentile | 1.456727202 |
| BRI | 2155 | Female | 15.12328767 | Old | 15.88 | 10.98 | 3.54 | 7 | 80-90th percentile | 0.899500759 |
| LAB | 3078 | Female | 7.69 | Young | 32.89 | 9.51 | -2.39 | 8 | NA | 1.258110174 |
| GSMD | 3784 | Female | 8.715068493 | Young | 57.83 | 9.34 | 0.84 | 2 | NA | 0.953463142 |
| LAB | 5439 | Male | 9.723287671 | Young | 32.89 | 9.51 | -0.36 | 8 | NA | 1.485053689 |
| ROT | 6559 | Female | 8.279452055 | Young | 47.5 | 8.12 | -0.51 | 2 | NA | 1.001828197 |
| SCOT | 8069 | Male | 8.77260274 | Young | 9 | 12.19 | -3.42 | 3 | NA | 2.051142949 |
| POD | 8264 | Male | 11.52876712 | Young | 5.67 | 10.33 | -0.96 | 9 | NA | 1.27571134 |
| WHI | 18810 | Male | 15.87671233 | Old | 14.74 | 13.67 | 4.19 | 10 | 90-95th percentile | 0.243827992 |
| BMD | 548822 | Male | 7.72 | Young | 44.23 | 6.88 | -1.36 | 2 | NA | 1.371930635 |
| FCR | 56517 | Male | 12.63287671 | Old | 29.48 | 8.90 | 2.25 | 8 | 90-95th percentile | 1.186012024 |
| BE | 1570_15 | Male | 13 | Old | 40.82 | 8.76 | 3.62 | 1 | <80th percentile | 0.953647764 |
| GD | 1058_19 | Male | 6.893150685 | Young | 66 | 5.41 | -0.26 | 2 | NA | 1.414244935 |
| BRI | 1535_19 | Male | 14.73972603 | Old | 15.88 | 10.98 | 3.15 | 7 | <80th percentile | 1.124118572 |
| GR | 1937 | Male | 1.19 | Young | 31.75 | 12.76 | -8.99 | 8 | NA | 0.742662157 |
| AI | 10790 | Female | 13.34794521 | Old | 45 | 10.78 | 4.34 | 5 | <80th percentile | 0.986495481 |
| AS | 11116 | Male | 5.983561644 | Young | 26.08 | 10.44 | -4.70 | 1 | NA | 0.979479364 |
| FCR | 11420 | Male | 7.997260274 | Young | 29.48 | 8.90 | -2.39 | 8 | NA | 0.9937221 |
| AI | 12729 | Male | 4.56 | Young | 45 | 10.78 | -4.45 | 5 | NA | 0.936548569 |
| BB | 13043 | Female | 10.4 | Young | 29.48 | 10.92 | 0.02 | 1 | NA | 1.01204122 |
| BB | 13172 | Male | 8.44 | Young | 29.48 | 10.92 | -1.94 | 1 | NA | 0.943236968 |
| BC | 1451 | Male | 15.58630137 | Old | 16 | 13.15 | 4.01 | 1 | 80-90th percentile | 0.976488916 |
| BB | 17640 | Female | 9.27 | Young | 29.48 | 10.92 | -1.11 | 1 | NA | 0.925733416 |
| BB | 18112 | Female | 14.83 | Old | 29.48 | 10.92 | 4.45 | 1 | 80-90th percentile | 0.862693258 |
| BC | 1862 | Female | 15.58356164 | Old | 16 | 13.15 | 4.01 | 1 | 80-90th percentile | 1.017318331 |
| FCR | 18708 | Male | 8.197260274 | Young | 29.48 | 8.90 | -2.19 | 8 | NA | 0.981321002 |
| BB | 18935 | Female | 16 | Old | 29.48 | 10.92 | 5.62 | 1 | 95-99th percentile | 0.935864672 |
| GSMD | 19268 | Female | 2.345205479 | Young | 57.83 | 9.34 | -5.53 | 2 | NA | 1.005761463 |
| BB | 19519 | Female | 14.5 | Old | 29.48 | 10.92 | 4.12 | 1 | 80-90th percentile | 0.902775477 |
| BB | 19870 | Male | 13.5 | Old | 29.48 | 10.92 | 3.12 | 1 | <80th percentile | 0.949146353 |
| BB | 19871 | Female | 14.5 | Old | 29.48 | 10.92 | 4.12 | 1 | 80-90th percentile | 0.961319909 |
| BB | 19921 | Female | 14.59 | Old | 29.48 | 10.92 | 4.21 | 1 | 80-90th percentile | 0.942468591 |
| BB | 19971 | Female | 14 | Old | 29.48 | 10.92 | 3.62 | 1 | <80th percentile | 0.917960867 |
| BB | 20003 | Male | 15.55 | Old | 29.48 | 10.92 | 5.17 | 1 | 90-95th percentile | 0.848403697 |
| LEO | 20021 | Female | 13.57 | Old | 58 | 6.96 | 5.71 | 2 | 95-99th percentile | 0.850467686 |
| BB | 20240 | Female | 17.9 | Old | 29.48 | 10.92 | 7.52 | 1 | >=99th percentile | 0.919428076 |
| BB | 20469 | Female | 11.35 | Young | 29.48 | 10.92 | 0.97 | 1 | NA | 0.752049308 |
| AS | 21421 | Female | 14.73 | Old | 26.08 | 10.44 | 4.05 | 1 | 95-99th percentile | 0.917014134 |
| BC | 21627 | Female | 18.37 | Old | 16 | 13.15 | 6.79 | 1 | >=99th percentile | 0.920726316 |
| GSD | 21681 | Male | 13.9 | Old | 35.15 | 8.26 | 4.02 | 1 | 80-90th percentile | 0.965679329 |
| BB | 22051 | Female | 10.52 | Young | 29.48 | 10.92 | 0.14 | 1 | NA | 0.827187509 |
| BB | 22052 | Female | 8.66 | Young | 29.48 | 10.92 | -1.72 | 1 | NA | 0.905478987 |
| GSMD | 3270 | Male | 8 | Young | 57.83 | 9.34 | 0.12 | 2 | NA | 0.844674135 |
| AS | 5642 | Female | 8.838356164 | Young | 26.08 | 10.44 | -1.85 | 1 | NA | 0.961046039 |
| POD | 5784 | Female | 5.08 | Young | 5.67 | 10.33 | -7.41 | 9 | NA | 0.915663171 |
| GSD | 5906 | Female | 13.04931507 | Old | 35.15 | 8.26 | 3.17 | 1 | <80th percentile | 0.979801215 |
| LAB | 6229 | Female | 9.660273973 | Young | 32.89 | 9.51 | -0.42 | 8 | NA | 0.970993296 |
| LAB | 6415 | Female | 9.312328767 | Young | 32.89 | 9.51 | -0.77 | 8 | NA | 0.975613951 |
| BB | 6453 | Female | 7.16 | Young | 29.48 | 10.92 | -3.22 | 1 | NA | 0.933402021 |
| AS | 7402 | Female | 14.35616438 | Old | 26.08 | 10.44 | 3.67 | 1 | 95-99th percentile | 0.995431248 |
| GSD | 8711 | Male | 9.339726027 | Young | 35.15 | 8.26 | -0.54 | 1 | NA | 0.990596951 |
| GD | 9479GD | Female | 6.77260274 | Young | 66 | 5.41 | -0.38 | 2 | NA | 0.926265464 |
| WHI | 13055 | Male | 11.64383562 | Young | 14.74 | 13.67 | -0.04 | 10 | NA | 0.848244538 |
| PS | 22961 | Male | 15.70958904 | Old | 10.21 | 12.94 | 3.62 | 1 | 80-90th percentile | 0.972789794 |
| GD | L15003862 | Female | 6.602739726 | Young | 66 | 5.41 | -0.55 | 2 | NA | 0.964049887 |
| BB | N0210502 | Female | 9.5 | Young | 29.48 | 10.92 | -0.88 | 1 | NA | 0.862896449 |
| BB | N025843 | Female | 15.61 | Old | 29.48 | 10.92 | 5.23 | 1 | 90-95th percentile | 0.858036197 |
| BB | N103759 | Male | 9.08 | Young | 29.48 | 10.92 | -1.30 | 1 | NA | 0.932686058 |
| BB | N147729 | Female | 9.42 | Young | 29.48 | 10.92 | -0.96 | 1 | NA | 0.880384285 |
| FB | T142108 | Male | 9.339726027 | Young | 12.7 | 8.86 | -2.53 | 9 | NA | 1.02493045 |

**Table S2.** Novel loci associated with breed-average body weight.

| **Lead variant** | **Locus region** | **Effect allele** | **β (kg)** | **SE (kg)** | **p** | **Significance** | **Nearest gene** | **Distance to nearest gene (kb)** | **Genes in locus region (RefSeq 105)** | **Shared with breed-median lifespan** |
|---|---|---|---|---|---|---|---|---|---|---|
| 1_33149841_A_C | chr1:32,430,694-33,839,127 | C | 5.16 | 1 | 3.7 × 10<sup>−7</sup> | genome-wide | NMBR | 622.8 | NMBR,VTA1 | no |
| 1_35227744_G_C | chr1:35,154,566-35,228,999 | C | 6.67 | 1.14 | 7.4 × 10<sup>−9</sup> | genome-wide | PHACTR2 | 0.0 | PHACTR2 | yes |
| 1_43712686_C_T | chr1:42,712,808-44,389,159 | T | 4.3 | 0.82 | 2.0 × 10<sup>−7</sup> | genome-wide | OPRM1 | 260.2 | SYNE1,MYCT1,LOC111095766,VIP,FBXO5,MTRF1L,RGS17,OPRM1,IPCEF1,CNKSR3 | no |
| 1_56500122_T_A | chr1:56,396,239-56,523,303 | A | 5.93 | 1.27 | 3.9 × 10<sup>−6</sup> | suggestive | THBS2 | 70.1 | THBS2 | no |
| 1_76923687_G_A | chr1:76,897,726-76,923,687 | A | 7.97 | 1.44 | 4.2 × 10<sup>−8</sup> | genome-wide | SPATA31D1 | 406.6 | intergenic: RASEF (429211 bp) \| SPATA31D1 (406602 bp) | yes |
| 2_56263214_G_A | chr2:56,053,882-56,702,966 | A | 3.43 | 0.66 | 2.7 × 10<sup>−7</sup> | genome-wide | BTF3 | 78.7 | TMEM174,BTF3,ANKRA2,UTP15,ARHGEF28 | yes |
| 2_58667278_T_C | chr2:58,667,278-58,667,278 | T | 1.94 | 0.42 | 3.9 × 10<sup>−6</sup> | suggestive | CNGB1 | 25.6 | intergenic: CNGB1 (25647 bp) \| KIFC3 (32917 bp) | no |
| 2_85001921_A_C | chr2:84,998,729-85,034,842 | C | 6.09 | 1.11 | 6.2 × 10<sup>−8</sup> | genome-wide | TARDBP | 3.8 | intergenic: TARDBP (593 bp) \| CASZ1 (159798 bp) | yes |
| 3_5444906_T_C | chr3:5,221,081-5,619,406 | C | 5.96 | 1.19 | 7.8 × 10<sup>−7</sup> | suggestive | EFNA5 | 898.4 | LOC106558441 | yes |
| 3_6953358_A_T | chr3:6,945,698-7,581,965 | T | 7.42 | 1.57 | 2.8 × 10<sup>−6</sup> | suggestive | NUDT12 | 740.5 | intergenic: LOC106558441 (1611155 bp), then EFNA5 (2399171 bp) \| NUDT12 (111938 bp) | no |
| 3_11923502_G_A | chr3:11,923,502-11,923,502 | G | 3.39 | 0.67 | 5.0 × 10<sup>−7</sup> | suggestive | RGMB | 496.2 | intergenic: RGMB (496185 bp) \| RIOK2 (625328 bp) | no |
| 3_33666192_A_AC | chr3:32,882,011-33,980,294 | AC | 9.63 | 1.55 | 9.6 × 10<sup>−10</sup> | genome-wide | GABRA5 | 0.0 | GABRG3,GABRA5,GABRB3 | yes |
| 3_60686332_G_A | chr3:60,686,332-61,350,015 | A | 3.4 | 0.73 | 3.7 × 10<sup>−6</sup> | suggestive | ADRA2C | 38.2 | LRPAP1,DOK7,LOC102155289,HGFAC,RGS12,MSANTD1,HTT,GRK4,NOP14 | no |
| 3_62527724_A_T | chr3:62,514,256-62,533,775 | T | 3.05 | 0.65 | 3.5 × 10<sup>−6</sup> | suggestive | FAM53A | 86.5 | intergenic: FAM53A (73067 bp) \| NKX1-1 (100879 bp) | no |
| 3_66766121_G_A | chr3:66,766,035-66,767,289 | A | 1.75 | 0.37 | 3.0 × 10<sup>−6</sup> | suggestive | RAB28 | 28.1 | intergenic: RAB28 (27980 bp) \| HS3ST1 (1539223 bp) | no |
| 4_64408937_T_G | chr4:64,398,732-64,409,575 | G | 6.42 | 1.22 | 1.9 × 10<sup>−7</sup> | genome-wide | EMB | 0.6 | EMB | no |
| 5_30811044_G_A | chr5:30,811,044-30,811,044 | A | 6.89 | 1.37 | 6.3 × 10<sup>−7</sup> | suggestive | PITPNM3 | 0.0 | PITPNM3 | yes |
| 5_33467613_G_A | chr5:33,467,613-34,145,636 | A | 2.33 | 0.43 | 7.2 × 10<sup>−8</sup> | genome-wide | MFSD6L | 0.0 | MFSD6L,PIK3R6,PIK3R5,NTN1,STX8,CFAP52,USP43 | yes |
| 5_52432695_A_G | chr5:50,638,529-53,230,715 | G | 5.7 | 1.08 | 1.7 × 10<sup>−7</sup> | genome-wide | DAB1 | 0.0 | JUN,MYSM1,TACSTD2,OMA1,DAB1,C8B,C8A,FYB2,PRKAA2,PLPP3 | yes |
| 5_59320181_T_A | chr5:58,432,005-60,709,911 | A | 8.79 | 1.48 | 5.1 × 10<sup>−9</sup> | genome-wide | AJAP1 | 297.1 | AJAP1,NPHP4,KCNAB2,CHD5,RPL22,RNF207,ICMT,HES3,GPR153,ACOT7,HES2,ESPN,TNFRSF25,PLEKHG5,NOL9,TAS1R1,ZBTB48,KLHL21,PHF13,THAP3,DNAJC11,CAMTA1 | yes |
| 5_62669351_CG_C | chr5:61,933,401-62,674,147 | C | 7.09 | 1.39 | 4.4 × 10<sup>−7</sup> | suggestive | SPSB1 | 0.0 | RERE,ENO1,CA6,LOC489640,GPR157,H6PD,SPSB1 | yes |
| 6_34484659_G_A | chr6:34,448,319-34,545,805 | A | 5.11 | 0.97 | 2.1 × 10<sup>−7</sup> | genome-wide | RBFOX1 | 0.0 | RBFOX1 | no |
| 6_42053530_T_A | chr6:42,050,552-42,496,238 | A | 7.07 | 1.19 | 5.1 × 10<sup>−9</sup> | genome-wide | CSF1 | 6.5 | EPS8L3,LOC479911,LOC479912,AMPD2,GNAT2,GNAI3,GPR61,AMIGO1,CYB561D1,ATXN7L2,SYPL2,PSMA5,SORT1 | yes |
| 6_45053965_ATCT_A | chr6:43,653,771-45,692,420 | A | 9.59 | 1.2 | 8.1 × 10<sup>−15</sup> | genome-wide | PRMT6 | 575.8 | VAV3,NTNG1,PRMT6 | yes |
| 6_47544353_C_G | chr6:47,282,478-48,449,813 | G | 9.84 | 1.53 | 2.5 × 10<sup>−10</sup> | genome-wide | COL11A1 | 0.0 | COL11A1,LOC106558952,OLFM3 | yes |
| 7_79775438_C_T | chr7:79,774,542-80,519,856 | T | 3.56 | 0.66 | 1.1 × 10<sup>−7</sup> | genome-wide | CTIF | 5.2 | CTIF,ZBTB7C | no |
| 8_15610914_G_T | chr8:15,594,442-15,625,182 | G | 1.77 | 0.37 | 1.8 × 10<sup>−6</sup> | suggestive | SLC25A21 | 0.0 | SLC25A21 | no |
| 8_26189272_TCTCA_T | chr8:25,925,668-26,197,386 | T | 6.53 | 1.39 | 3.0 × 10<sup>−6</sup> | suggestive | RPS29 | 32.3 | intergenic: MDGA2 (1317665 bp) \| RPS29 (24160 bp) | no |
| 8_47234336_TA_T | chr8:47,234,336-47,234,336 | TA | 5.13 | 1.05 | 1.3 × 10<sup>−6</sup> | suggestive | ENTPD5 | 0.0 | ENTPD5 | no |
| 9_26474422_C_T | chr9:26,456,769-26,667,810 | T | 8.96 | 1.82 | 1.0 × 10<sup>−6</sup> | suggestive | SPATA20 | 0.0 | EPN3,SPATA20,CACNA1G,ABCC3,ANKRD40,LUC7L3,LOC100856485 | yes |
| 9_44584685_AAGCTCTCCAG_A | chr9:44,454,785-44,584,685 | A | 7.06 | 1.32 | 1.1 × 10<sup>−7</sup> | genome-wide | BHLHA9 | 0.0 | CPD,GOSR1,TUSC5,BHLHA9 | yes |
| 9_50389203_G_A | chr9:50,389,203-50,659,824 | A | 5.88 | 1.16 | 5.7 × 10<sup>−7</sup> | suggestive | WDR5 | 18.0 | RXRA,LOC102151988 | no |
| 10_4168444_C_A | chr10:4,098,339-5,417,188 | C | 2.33 | 0.43 | 7.0 × 10<sup>−8</sup> | genome-wide | SLC16A7 | 667.8 | FAM19A2 | no |
| 12_10656937_G_A | chr12:10,580,658-10,656,937 | A | 3.3 | 0.69 | 2.1 × 10<sup>−6</sup> | suggestive | CCND3 | 0.0 | MED20,BYSL,CCND3 | no |
| 12_31990581_A_G | chr12:31,865,370-32,006,785 | A | 3.28 | 0.67 | 1.2 × 10<sup>−6</sup> | suggestive | ADGRB3 | 0.0 | ADGRB3 | no |
| 12_34483222_AGG_A | chr12:34,482,003-34,483,222 | AGG | 2.06 | 0.43 | 2.3 × 10<sup>−6</sup> | suggestive | RIMS1 | 0.0 | RIMS1 | no |
| 13_7358534_G_A | chr13:6,777,044-7,532,022 | G | 3.22 | 0.57 | 2.0 × 10<sup>−8</sup> | genome-wide | OXR1 | 0.0 | ZFPM2,OXR1 | no |
| 13_10014673_G_A | chr13:9,168,016-10,171,337 | G | 3.22 | 0.53 | 2.5 × 10<sup>−9</sup> | genome-wide | PKHD1L1 | 0.0 | EMC2,TMEM74,TRHR,NUDCD1,ENY2,PKHD1L1,LOC111098647,EBAG9,SYBU | no |
| 14_30056204_C_G | chr14:30,056,204-30,063,061 | G | 4.86 | 1.03 | 3.0 × 10<sup>−6</sup> | suggestive | AGMO | 0.0 | AGMO | no |
| 15_29445083_C_T | chr15:29,144,462-30,409,424 | T | 6.15 | 1.12 | 5.7 × 10<sup>−8</sup> | genome-wide | TMTC3 | 94.6 | C15H12orf50,C15H12orf29,CEP290,LOC111090022,TMTC3,KITLG,LOC102155235,DUSP6,POC1B | yes |
| 15_33535412_T_C | chr15:32,571,129-33,618,684 | C | 5.56 | 0.94 | 6.5 × 10<sup>−9</sup> | genome-wide | NUDT4 | 92.0 | BTG1,C15H12orf74,PLEKHG7,EEA1,LOC111090036 | yes |
| 17_21063000_C_T | chr17:20,112,778-21,352,434 | T | 9.31 | 1.57 | 5.0 × 10<sup>−9</sup> | genome-wide | AGBL5 | 0.0 | KIF3C,RAB10,GAREM2,HADHA,HADHB,ADGRF3,SELENOI,DRC1,OTOF,C17H2orf70,CIB4,KCNK3,SLC35F6,CENPA,DPYSL5,MAPRE3,TMEM214,AGBL5,OST4,EMILIN1,KHK,CGREF1,ABHD1,PREB,TCF23,SLC5A6,ATRAID,CAD,SLC30A3,DNAJC5G,TRIM54,UCN,MPV17,GTF3C2,EIF2B4,SNX17,ZNF513,PPM1G | yes |
| 17_25465646_A_G | chr17:23,464,957-25,833,826 | G | 9.21 | 1.62 | 2.0 × 10<sup>−8</sup> | genome-wide | SLC30A6 | 0.0 | ALK,YPEL5,LBH,LCLAT1,CAPN13,GALNT14,CAPN14,EHD3,XDH,SRD5A2,MEMO1,DPY30,SPAST,SLC30A6,NLRC4,YIPF4,BIRC6 | yes |
| 17_32499604_T_C | chr17:32,359,461-33,320,924 | C | 7.29 | 1.36 | 1.1 × 10<sup>−7</sup> | genome-wide | SLC8A1 | 118.2 | SLC8A1 | yes |
| 20_22129914_A_ACT | chr20:22,129,914-22,129,914 | ACT | 1.86 | 0.39 | 2.9 × 10<sup>−6</sup> | suggestive | FRMD4B | 123.7 | intergenic: MITF (256369 bp) \| FRMD4B (123705 bp) | no |
| 21_36964264_ATT_A | chr21:36,845,880-37,962,777 | A | 7.14 | 1.29 | 4.7 × 10<sup>−8</sup> | genome-wide | SPON1 | 0.0 | SPON1,RRAS2,COPB1,LOC111091601,PSMA1,PDE3B,CYP2R1,LOC100683373,CALCB,CRSP-3,CRSP-2,CALCA,INSC | yes |
| 22_38824845_G_A | chr22:33,647,918-39,246,151 | A | 10.15 | 1.37 | 4.8 × 10<sup>−13</sup> | genome-wide | SLITRK5 | 556.4 | C22H16orf87,SLITRK1,LOC111091674,SLITRK6 | yes |
| 22_61116972_C_T | chr22:60,960,241-61,146,556 | T | 4.86 | 0.91 | 1.1 × 10<sup>−7</sup> | genome-wide | RASA3 | 0.0 | TMEM255B,GAS6,LOC106557589,RASA3 | no |
| 26_36728098_C_T | chr26:36,494,840-36,781,345 | T | 8.82 | 1.36 | 1.8 × 10<sup>−10</sup> | genome-wide | PRKG1 | 0.0 | PRKG1 | yes |
| 26_38270013_G_A | chr26:38,267,731-38,271,587 | A | 4.22 | 0.91 | 3.9 × 10<sup>−6</sup> | suggestive | RNLS | 0.0 | RNLS | no |
| 27_37904404_C_A | chr27:37,900,587-37,923,285 | A | 6.97 | 1.28 | 7.2 × 10<sup>−8</sup> | genome-wide | CLSTN3 | 0.0 | CLSTN3,RBP5 | yes |
| 29_4270906_C_T | chr29:2,486,010-4,270,906 | T | 7 | 1.21 | 1.3 × 10<sup>−8</sup> | genome-wide | ST18 | 4.3 | SNTG1,PXDNL,PCMTD1,ST18,LOC106557997 | yes |
| 34_25485833_C_T | chr34:25,485,833-25,496,728 | C | 2.84 | 0.59 | 1.8 × 10<sup>−6</sup> | suggestive | MFSD1 | 391.6 | LOC478678 | no |
| 37_8373862_T_A | chr37:8,326,677-8,377,782 | A | 4.36 | 0.94 | 4.0 × 10<sup>−6</sup> | suggestive | SATB2 | 174.4 | intergenic: PLCL1 (751248 bp) \| SATB2 (170513 bp) | no |
| 37_9401025_G_A | chr37:9,389,323-12,817,585 | A | 5.62 | 1.1 | 4.3 × 10<sup>−7</sup> | suggestive | SPATS2L | 2.5 | SPATS2L,KCTD18,SGO2,AOX4,AOX2,BZW1,LOC100684200,CLK1,PPIL3,NIF3L1,ORC2,FAM126B,NDUFB3,CFLAR,CASP10,CASP8,ALS2CR12,TRAK2,STRADB,C2CD6,TMEM237,MPP4,ALS2,CDK15,FZD7,KIAA2012,SUMO1,NOP58,BMPR2,FAM117B,ICA1L,WDR12,CARF,NBEAL1,LOC100686469,ABI2,RAPH1,CD28,CTLA4,ICOS | yes |
| 37_15881879_C_T | chr37:15,881,879-16,657,797 | T | 7.45 | 1.48 | 6.8 × 10<sup>−7</sup> | suggestive | CREB1 | 39.3 | CREB1,METTL21A,CCNYL1,FZD5,PLEKHM3,LOC609894,CRYGD,CRYGB,CRYGA,C37H2orf80,IDH1,PIKFYVE,PTH2R | yes |
| 38_22632135_G_A | chr38:22,630,733-22,632,135 | A | 5.61 | 1.2 | 3.5 × 10<sup>−6</sup> | suggestive | OR10J5 | 0.0 | OR10J5 | no |
| X_107820317_T_TG | chrX:107,199,224-107,823,367 | T | 1.7 | 0.35 | 1.3 × 10<sup>−6</sup> | suggestive | ZIC3 | 1.7 | LOC611979,GPR101,ZIC3 | no |

**Table S3.** Breed-average body weight loci shared with breed-median lifespan.

| Weight lead | Locus region | Nearest gene | Effect allele | β weight (kg) | p weight | Significance (weight) | β lifespan (years) | p lifespan | Lifespan lead(s) in locus | Status |
|---|---|---|---|---|---|---|---|---|---|---|
| 1_35227744_G_C | chr1:35,154,566-35,228,999 | PHACTR2 (0.0 kb) | C | 6.67 | 7.4 × 10<sup>−9</sup> | genome-wide | -0.73 | 4.3 × 10<sup>−7</sup> | 1_35227744_G_C (r2 = 1.00) | novel |
| 1_76923687_G_A | chr1:76,897,726-76,923,687 | SPATA31D1 (406.6 kb) | A | 7.97 | 4.2 × 10<sup>−8</sup> | genome-wide | -0.9 | 7.4 × 10<sup>−7</sup> | 1_76923687_G_A (r2 = 1.00) | novel |
| 2_56263214_G_A | chr2:56,053,882-56,702,966 | BTF3 (78.7 kb) | A | 3.43 | 2.7 × 10<sup>−7</sup> | genome-wide | -0.36 | 1.8 × 10<sup>−5</sup> | 2_56702966_A_G (r2 = 0.48) | novel |
| 2_85001921_A_C | chr2:84,998,729-85,034,842 | TARDBP (3.8 kb) | C | 6.09 | 6.2 × 10<sup>−8</sup> | genome-wide | -0.7 | 7.1 × 10<sup>−7</sup> | 2_85028085_C_T (r2 = 0.90) | novel |
| 3_5444906_T_C | chr3:5,221,081-5,619,406 | EFNA5 (898.4 kb) | C | 5.96 | 7.8 × 10<sup>−7</sup> | suggestive | -0.71 | 2.4 × 10<sup>−6</sup> | 3_5619406_G_A (r2 = 0.85) | novel |
| 3_33666192_A_AC | chr3:32,882,011-33,980,294 | GABRA5 (0.0 kb) | AC | 9.63 | 9.6 × 10<sup>−10</sup> | genome-wide | -1.15 | 4.9 × 10<sup>−9</sup> | 3_33666192_A_AC (r2 = 1.00) | novel |
| 5_30811044_G_A | chr5:30,811,044-30,811,044 | PITPNM3 (0.0 kb) | A | 6.89 | 6.3 × 10<sup>−7</sup> | suggestive | -0.84 | 1.4 × 10<sup>−6</sup> | 5_30811044_G_A (r2 = 1.00) | novel |
| 5_33467613_G_A | chr5:33,467,613-34,145,636 | MFSD6L (0.0 kb) | A | 2.33 | 7.2 × 10<sup>−8</sup> | genome-wide | -0.23 | 1.7 × 10<sup>−5</sup> | 5_34145636_G_A (r2 = 0.20) | novel |
| 5_52432695_A_G | chr5:50,638,529-53,230,715 | DAB1 (0.0 kb) | G | 5.7 | 1.7 × 10<sup>−7</sup> | genome-wide | -0.64 | 3.1 × 10<sup>−6</sup> | 5_53230715_C_A (r2 = 0.70) | novel |
| 5_59320181_T_A | chr5:58,432,005-60,709,911 | AJAP1 (297.1 kb) | A | 8.79 | 5.1 × 10<sup>−9</sup> | genome-wide | -1.04 | 3.6 × 10<sup>−8</sup> | 5_59320181_T_A (r2 = 1.00); 5_60709904_T_TC (r2 = 0.92) | novel |
| 5_62669351_CG_C | chr5:61,933,401-62,674,147 | SPSB1 (0.0 kb) | C | 7.09 | 4.4 × 10<sup>−7</sup> | suggestive | -0.82 | 3.1 × 10<sup>−6</sup> | 5_62669351_CG_C (r2 = 1.00) | novel |
| 6_42053530_T_A | chr6:42,050,552-42,496,238 | CSF1 (6.5 kb) | A | 7.07 | 5.1 × 10<sup>−9</sup> | genome-wide | -0.83 | 4.2 × 10<sup>−8</sup> | 6_42053530_T_A (r2 = 1.00) | novel |
| 6_45053965_ATCT_A | chr6:43,653,771-45,692,420 | PRMT6 (575.8 kb) | A | 9.59 | 8.1 × 10<sup>−15</sup> | genome-wide | -0.95 | 1.2 × 10<sup>−9</sup> | 6_43912554_G_A (r2 = 0.63) | novel |
| 6_47544353_C_G | chr6:47,282,478-48,449,813 | COL11A1 (0.0 kb) | G | 9.84 | 2.5 × 10<sup>−10</sup> | genome-wide | -1.13 | 7.1 × 10<sup>−9</sup> | 6_47544353_C_G (r2 = 1.00) | novel |
| 9_26474422_C_T | chr9:26,456,769-26,667,810 | SPATA20 (0.0 kb) | T | 8.96 | 1.0 × 10<sup>−6</sup> | suggestive | -1.06 | 3.9 × 10<sup>−6</sup> | 9_26474422_C_T (r2 = 1.00) | novel |
| 9_44584685_AAGCTCTCCAG_A | chr9:44,454,785-44,584,685 | BHLHA9 (0.0 kb) | A | 7.06 | 1.1 × 10<sup>−7</sup> | genome-wide | -0.83 | 7.0 × 10<sup>−7</sup> | 9_44584685_AAGCTCTCCAG_A (r2 = 1.00) | novel |
| 15_29445083_C_T | chr15:29,144,462-30,409,424 | TMTC3 (94.6 kb) | T | 6.15 | 5.7 × 10<sup>−8</sup> | genome-wide | -0.77 | 6.0 × 10<sup>−8</sup> | 15_29445083_C_T (r2 = 1.00) | novel |
| 15_33535412_T_C | chr15:32,571,129-33,618,684 | NUDT4 (92.0 kb) | C | 5.56 | 6.5 × 10<sup>−9</sup> | genome-wide | -0.59 | 8.4 × 10<sup>−7</sup> | 15_33618684_C_T (r2 = 0.97) | novel |
| 17_21063000_C_T | chr17:20,112,778-21,352,434 | AGBL5 (0.0 kb) | T | 9.31 | 5.0 × 10<sup>−9</sup> | genome-wide | -1.1 | 3.9 × 10<sup>−8</sup> | 17_21048401_G_A (r2 = 1.00) | novel |
| 17_25465646_A_G | chr17:23,464,957-25,833,826 | SLC30A6 (0.0 kb) | G | 9.21 | 2.0 × 10<sup>−8</sup> | genome-wide | -1.08 | 1.4 × 10<sup>−7</sup> | 17_25465646_A_G (r2 = 1.00) | novel |
| 17_32499604_T_C | chr17:32,359,461-33,320,924 | SLC8A1 (118.2 kb) | C | 7.29 | 1.1 × 10<sup>−7</sup> | genome-wide | -0.85 | 8.7 × 10<sup>−7</sup> | 17_32499604_T_C (r2 = 1.00) | novel |
| 21_36964264_ATT_A | chr21:36,845,880-37,962,777 | SPON1 (0.0 kb) | A | 7.14 | 4.7 × 10<sup>−8</sup> | genome-wide | -0.83 | 3.4 × 10<sup>−7</sup> | 21_37464889_CA_C (r2 = 0.87) | novel |
| 22_38824845_G_A | chr22:33,647,918-39,246,151 | SLITRK5 (556.4 kb) | A | 10.15 | 4.8 × 10<sup>−13</sup> | genome-wide | -1.2 | 1.0 × 10<sup>−11</sup> | 22_38824845_G_A (r2 = 1.00); 22_33818773_C_T (r2 = 0.71) | novel |
| 26_36728098_C_T | chr26:36,494,840-36,781,345 | PRKG1 (0.0 kb) | T | 8.82 | 1.8 × 10<sup>−10</sup> | genome-wide | -1.05 | 1.2 × 10<sup>−9</sup> | 26_36728098_C_T (r2 = 1.00) | novel |
| 27_37904404_C_A | chr27:37,900,587-37,923,285 | CLSTN3 (0.0 kb) | A | 6.97 | 7.2 × 10<sup>−8</sup> | genome-wide | -0.91 | 1.7 × 10<sup>−8</sup> | 27_37904404_C_A (r2 = 1.00) | novel |
| 29_4270906_C_T | chr29:2,486,010-4,270,906 | ST18 (4.3 kb) | T | 7 | 1.3 × 10<sup>−8</sup> | genome-wide | -0.79 | 3.3 × 10<sup>−7</sup> | 29_2486010_CTCTCTA_C (r2 = 0.76) | novel |
| 37_9401025_G_A | chr37:9,389,323-12,817,585 | SPATS2L (2.5 kb) | A | 5.62 | 4.3 × 10<sup>−7</sup> | suggestive | -0.71 | 3.3 × 10<sup>−7</sup> | 37_9401025_G_A (r2 = 1.00) | novel |
| 37_15881879_C_T | chr37:15,881,879-16,657,797 | CREB1 (39.3 kb) | T | 7.45 | 6.8 × 10<sup>−7</sup> | suggestive | -0.89 | 2.0 × 10<sup>−6</sup> | 37_15881879_C_T (r2 = 1.00) | novel |
| X_82353310_G_A | chrX:82,289,784-86,758,710 | IRS4 (60.9 kb) | A | 2.35 | 2.6 × 10<sup>−10</sup> | genome-wide | -0.22 | 2.6 × 10<sup>−6</sup> | X_82326402_A_G (r2 = 0.89) | known |
| X_101730294_C_A | chrX:101,584,166-104,238,654 | ENOX2 (12.1 kb) | C | 2.69 | 2.3 × 10<sup>−10</sup> | genome-wide | -0.36 | 1.4 × 10<sup>−11</sup> | X_101730294_C_A (r2 = 1.00) | known |

One row per breed-average body weight locus overlapping at least one breed-median lifespan locus. Effect allele, the allele associated with higher breed-average body weight; both β are given for this allele at the breed-weight lead variant (β lifespan, change in breed-median lifespan per copy). Significance refers to the breed-weight association (genome-wide, p < 4 × 10<sup>−7</sup>; suggestive, 4 × 10<sup>−7</sup> ≤ p < 4 × 10<sup>−6</sup>). Lifespan lead(s) in locus, lead variant(s) of the overlapping breed-lifespan locus or loci, with their linkage disequilibrium (r<sup>2</sup>) with the breed-weight lead. Nearest gene, nearest named protein-coding gene to the lead variant (RefSeq 105), with its distance. Status, known or novel as defined in Methods.

**Table S4.** Breed-median lifespan loci not shared with breed-average body weight.

| **Lead variant** | **Locus region** | **Effect allele** | **β (years)** | **SE (years)** | **p** | **Significance** | **Nearest gene** | **Distance to nearest gene (kb)** | **Genes in locus region (RefSeq 105)** |
|---|---|---|---|---|---|---|---|---|---|
| 1_88390765_C_T | chr1:88,386,958-88,392,020 | T | 0.23 | 0.05 | 9.7 × 10<sup>−7</sup> | suggestive | PIP5K1B | 0.0 | PIP5K1B |
| 3_40184160_T_A | chr3:40,184,160-40,184,160 | T | 0.73 | 0.15 | 2.3 × 10<sup>−6</sup> | suggestive | ALDH1A3 | 15.5 | intergenic: ALDH1A3 (15474 bp) \| OTUB1 (41371 bp) |
| 3_79853426_A_C | chr3:79,853,426-80,100,643 | C | 0.23 | 0.05 | 2.2 × 10<sup>−6</sup> | suggestive | PCDH7 | 120.3 | PCDH7 |
| 5_67902007_G_T | chr5:67,902,007-67,907,182 | G | 0.47 | 0.09 | 2.0 × 10<sup>−7</sup> | genome-wide | COTL1 | 0.0 | COTL1 |
| 8_5336411_C_T | chr8:5,328,493-5,358,425 | T | 0.24 | 0.05 | 1.8 × 10<sup>−6</sup> | suggestive | STXBP6 | 458.8 | intergenic: STXBP6 (450904 bp) \| NOVA1 (775183 bp) |
| 12_63316149_C_A | chr12:63,303,797-63,316,149 | A | 0.36 | 0.07 | 2.4 × 10<sup>−7</sup> | genome-wide | PRDM1 | 138.1 | intergenic: PREP (482550 bp) \| PRDM1 (138093 bp) |
| 14_39547351_A_T | chr14:38,892,513-39,560,723 | T | 1.25 | 0.22 | 3.2 × 10<sup>−8</sup> | genome-wide | SNX10 | 20.4 | NFE2L3,HNRNPA2B1,CBX3 |
| 19_39597358_G_A | chr19:39,500,100-39,600,793 | A | 1.13 | 0.23 | 7.3 × 10<sup>−7</sup> | suggestive | THSD7B | 0.0 | THSD7B |
| 22_13741019_A_G | chr22:13,349,122-13,776,342 | A | 0.5 | 0.1 | 2.2 × 10<sup>−6</sup> | suggestive | PCDH17 | 124.1 | intergenic: LOC100855527 (3018482 bp), then OLFM4 (3273363 bp) \| PCDH17 (88796 bp) |
| 22_42852951_AT_A | chr22:42,807,136-43,029,138 | AT | 0.78 | 0.17 | 3.2 × 10<sup>−6</sup> | suggestive | GPC5 | 0.0 | GPC5 |
| 22_55051045_G_A | chr22:55,035,412-55,105,394 | G | 0.46 | 0.1 | 1.7 × 10<sup>−6</sup> | suggestive | EFNB2 | 516.9 | intergenic: SLC10A2 (2493436 bp) \| EFNB2 (462598 bp) |
| 25_26798516_T_G | chr25:26,739,177-26,829,682 | G | 0.24 | 0.05 | 3.0 × 10<sup>−6</sup> | suggestive | MTMR9 | 173.6 | intergenic: MTMR9 (114229 bp) \| XKR6 (154956 bp) |
| 26_24247930_G_A | chr26:24,247,930-24,396,272 | A | 1.1 | 0.21 | 3.1 × 10<sup>−7</sup> | genome-wide | PIK3IP1 | 0.0 | PIK3IP1,PATZ1,LOC106557811,DRG1,EIF4ENIF1 |
| 26_32891121_A_AT | chr26:32,891,121-32,895,026 | A | 0.42 | 0.09 | 1.2 × 10<sup>−6</sup> | suggestive | ZWINT | 431.7 | intergenic: ZWINT (431744 bp) \| LOC106557849 (287202 bp), then PCDH15 (531702 bp) |
| 27_35174492_C_T | chr27:35,171,601-35,174,492 | C | 0.4 | 0.09 | 3.2 × 10<sup>−6</sup> | suggestive | KLRA1 | 20.8 | intergenic: KLRA1 (17912 bp) \| LOC100683252 (11191 bp), then KLRK1 (460446 bp) |
| 28_14690957_C_T | chr28:14,690,957-14,690,957 | C | 0.49 | 0.1 | 2.4 × 10<sup>−6</sup> | suggestive | NOLC1 | 0.0 | NOLC1 |
| 28_16439808_A_G | chr28:16,437,817-16,446,380 | G | 0.42 | 0.09 | 2.5 × 10<sup>−6</sup> | suggestive | SFR1 | 0.0 | SFR1,CFAP43 |
| 30_37330334_C_T | chr30:37,330,334-37,330,334 | C | 0.21 | 0.05 | 4.0 × 10<sup>−6</sup> | suggestive | ISLR | 1.0 | intergenic: ISLR (982 bp) \| STRA6 (1500 bp) |
| 32_14035945_T_C | chr32:14,035,232-14,106,052 | C | 0.3 | 0.06 | 3.0 × 10<sup>−6</sup> | suggestive | CCSER1 | 0.0 | CCSER1 |
| 36_16760391_G_A | chr36:16,756,631-16,763,784 | G | 0.4 | 0.08 | 3.9 × 10<sup>−7</sup> | genome-wide | ITGA6 | 17.0 | intergenic: LOC111094202 (5213 bp), then DLX2 (254266 bp) \| ITGA6 (13579 bp) |
| 36_30408387_C_A | chr36:30,322,191-30,408,387 | C | 0.86 | 0.18 | 2.1 × 10<sup>−6</sup> | suggestive | COL3A1 | 79.9 | intergenic: GULP1 (73386 bp) \| COL3A1 (79863 bp) |

**Table S5.**
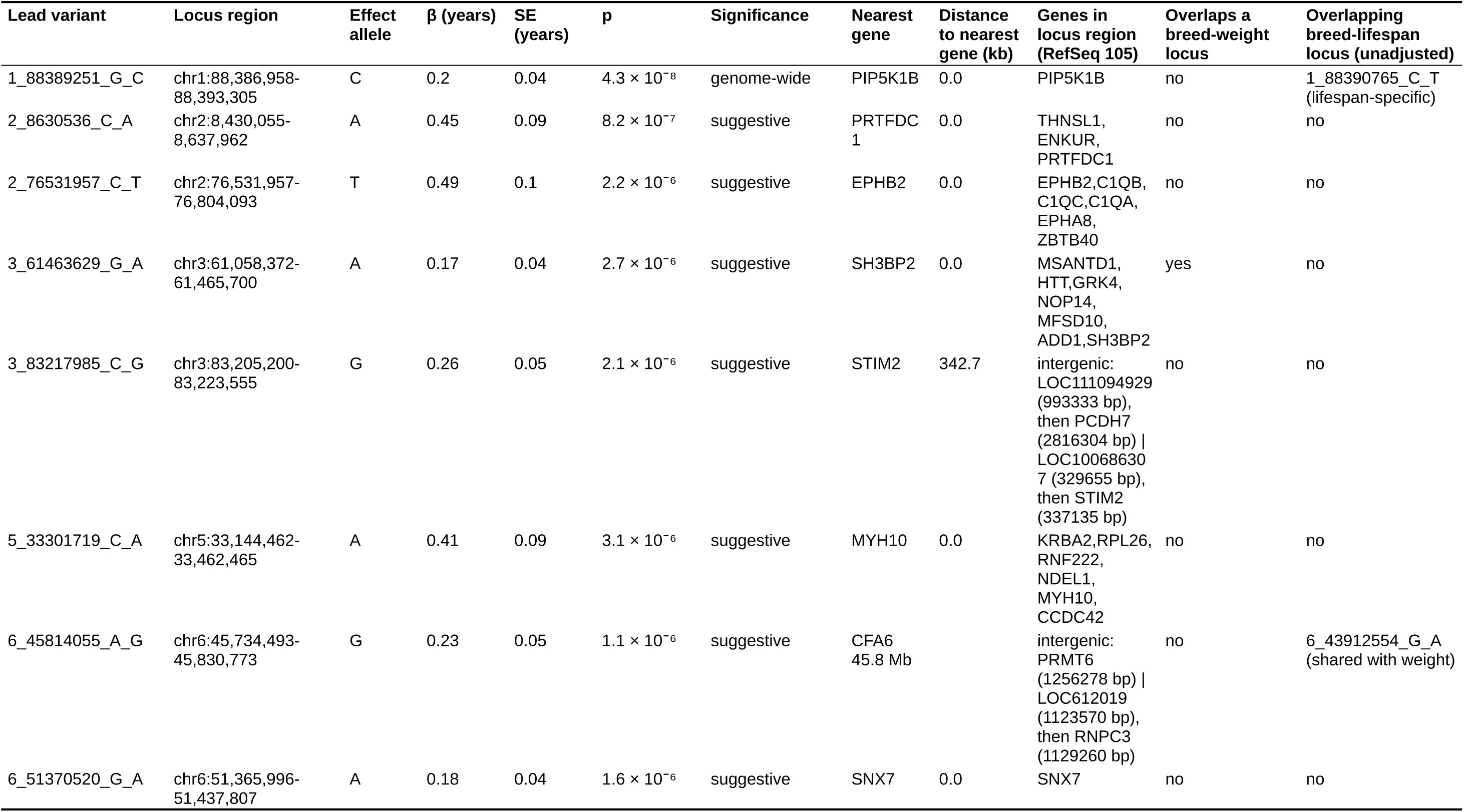
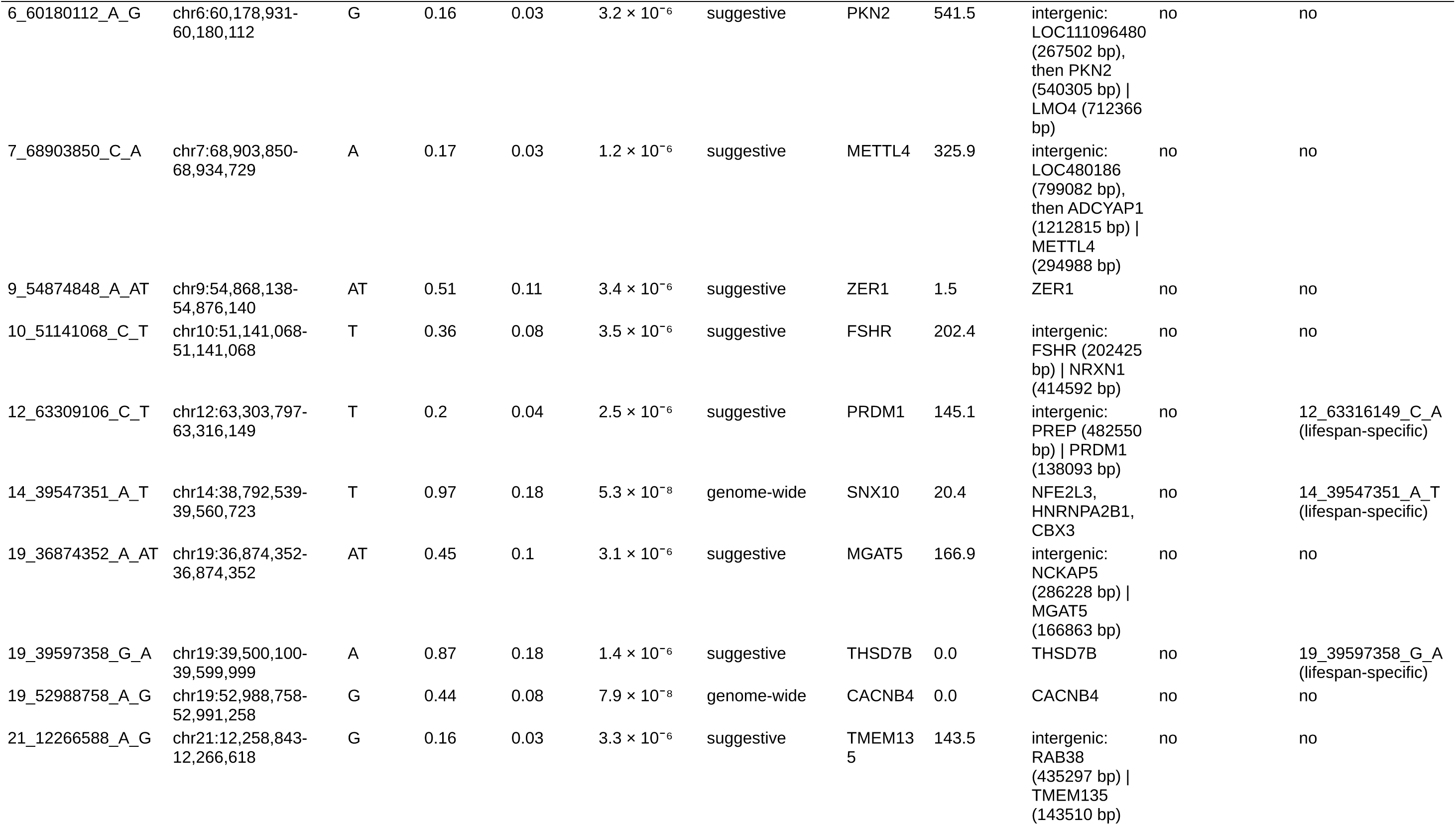
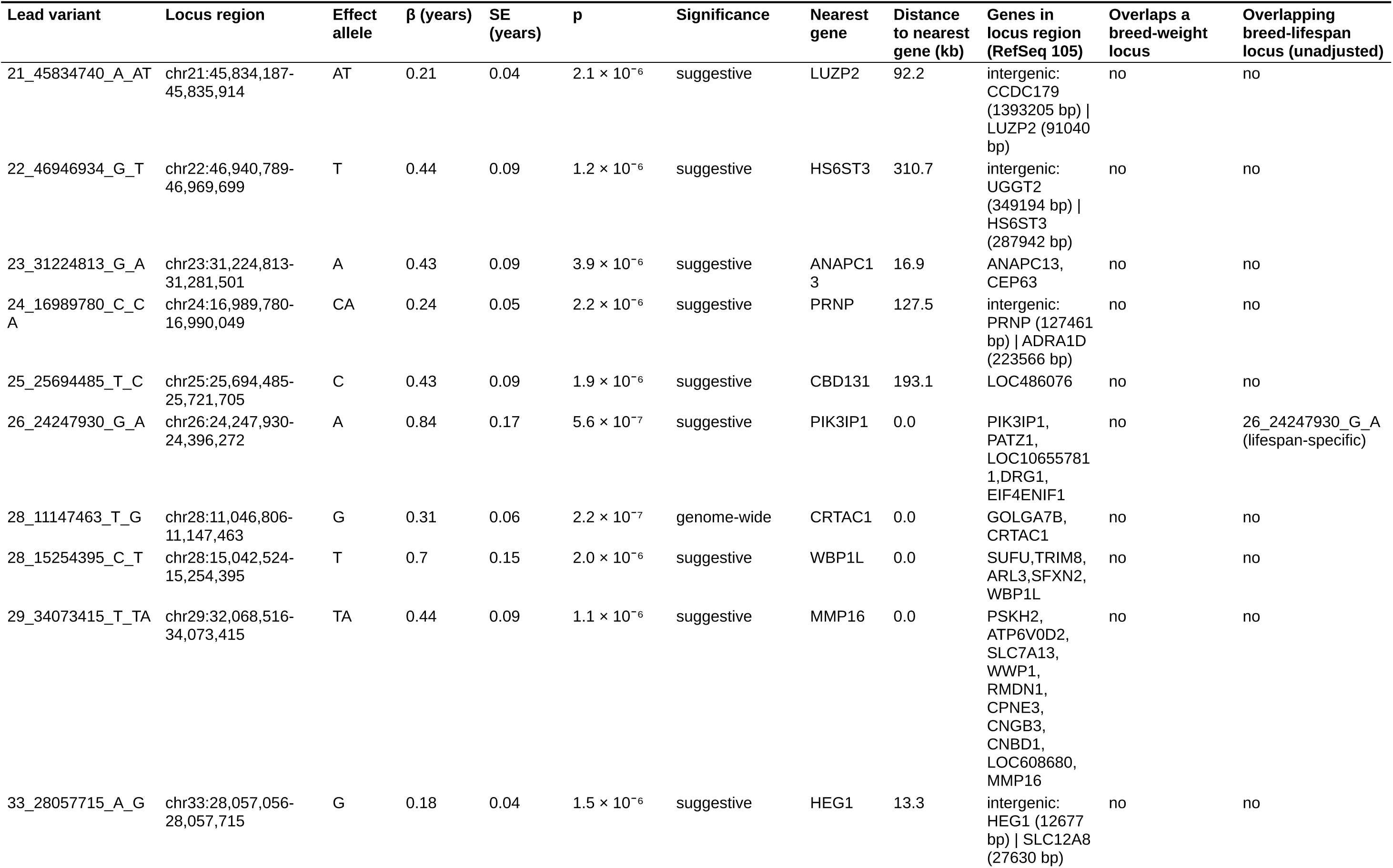
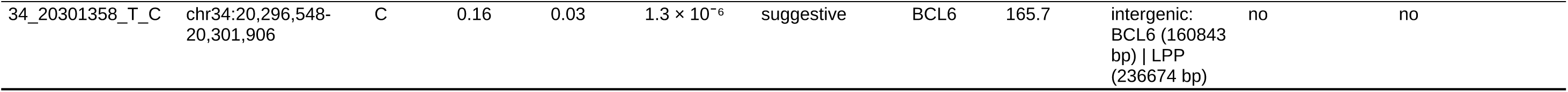
Loci associated with breed-median lifespan adjusted for breed-average body weight.

| **Lead variant** | **Locus region** | **Effect allele** | **β (years)** | **SE (years)** | **p** | **Significance** | **Nearest gene** | **Distance to nearest gene (kb)** | **Genes in locus region (RefSeq 105)** | **Overlaps a breed-weight locus** | **Overlapping breed-lifespan locus (unadjusted)** |
|---|---|---|---|---|---|---|---|---|---|---|---|
| 1_88389251_G_C | chr1:88,386,958-88,393,305 | C | 0.2 | 0.04 | 4.3 × 10<sup>−8</sup> | genome-wide | PIP5K1B | 0.0 | PIP5K1B | no | 1_88390765_C_T (lifespan-specific) |
| 2_8630536_C_A | chr2:8,430,055-8,637,962 | A | 0.45 | 0.09 | 8.2 × 10<sup>−7</sup> | suggestive | PRTFDC1 | 0.0 | THNSL1, ENKUR, PRTFDC1 | no | no |
| 2_76531957_C_T | chr2:76,531,957-76,804,093 | T | 0.49 | 0.1 | 2.2 × 10<sup>−6</sup> | suggestive | EPHB2 | 0.0 | EPHB2,C1QB, C1QC,C1QA, EPHA8, ZBTB40 | no | no |
| 3_61463629_G_A | chr3:61,058,372-61,465,700 | A | 0.17 | 0.04 | 2.7 × 10<sup>−6</sup> | suggestive | SH3BP2 | 0.0 | MSANTD1, HTT,GRK4, NOP14, MFSD10, ADD1,SH3BP2 | yes | no |
| 3_83217985_C_G | chr3:83,205,200-83,223,555 | G | 0.26 | 0.05 | 2.1 × 10<sup>−6</sup> | suggestive | STIM2 | 342.7 | intergenic: LOC111094929 (993333 bp), then PCDH7 (2816304 bp) \| LOC100686307 (329655 bp), then STIM2 (337135 bp) | no | no |
| 5_33301719_C_A | chr5:33,144,462-33,462,465 | A | 0.41 | 0.09 | 3.1 × 10<sup>−6</sup> | suggestive | MYH10 | 0.0 | KRBA2,RPL26, RNF222, NDEL1, MYH10, CCDC42 | no | no |
| 6_45814055_A_G | chr6:45,734,493-45,830,773 | G | 0.23 | 0.05 | 1.1 × 10<sup>−6</sup> | suggestive | CFA6 45.8 Mb |  | intergenic: PRMT6 (1256278 bp) \| LOC612019 (1123570 bp), then RNPC3 (1129260 bp) | no | 6_43912554_G_A (shared with weight) |
| 6_51370520_G_A | chr6:51,365,996-51,437,807 | A | 0.18 | 0.04 | 1.6 × 10<sup>−6</sup> | suggestive | SNX7 | 0.0 | SNX7 | no | no |
| 6_60180112_A_G | chr6:60,178,931-60,180,112 | G | 0.16 | 0.03 | 3.2 × 10<sup>−6</sup> | suggestive | PKN2 | 541.5 | intergenic: LOC111096480 (267502 bp), then PKN2 (540305 bp) \| LMO4 (712366 bp) | no | no |
| 7_68903850_C_A | chr7:68,903,850-68,934,729 | A | 0.17 | 0.03 | 1.2 × 10<sup>−6</sup> | suggestive | METTL4 | 325.9 | intergenic: LOC480186 (799082 bp), then ADCYAP1 (1212815 bp) \| METTL4 (294988 bp) | no | no |
| 9_54874848_A_AT | chr9:54,868,138-54,876,140 | AT | 0.51 | 0.11 | 3.4 × 10<sup>−6</sup> | suggestive | ZER1 | 1.5 | ZER1 | no | no |
| 10_51141068_C_T | chr10:51,141,068-51,141,068 | T | 0.36 | 0.08 | 3.5 × 10<sup>−6</sup> | suggestive | FSHR | 202.4 | intergenic: FSHR (202425 bp) \| NRXN1 (414592 bp) | no | no |
| 12_63309106_C_T | chr12:63,303,797-63,316,149 | T | 0.2 | 0.04 | 2.5 × 10<sup>−6</sup> | suggestive | PRDM1 | 145.1 | intergenic: PREP (482550 bp) \| PRDM1 (138093 bp) | no | 12_63316149_C_A (lifespan-specific) |
| 14_39547351_A_T | chr14:38,792,539-39,560,723 | T | 0.97 | 0.18 | 5.3 × 10<sup>−8</sup> | genome-wide | SNX10 | 20.4 | NFE2L3, HNRNPA2B1, CBX3 | no | 14_39547351_A_T (lifespan-specific) |
| 19_36874352_A_AT | chr19:36,874,352-36,874,352 | AT | 0.45 | 0.1 | 3.1 × 10<sup>−6</sup> | suggestive | MGAT5 | 166.9 | intergenic: NCKAP5 (286228 bp) \| MGAT5 (166863 bp) | no | no |
| 19_39597358_G_A | chr19:39,500,100-39,599,999 | A | 0.87 | 0.18 | 1.4 × 10<sup>−6</sup> | suggestive | THSD7B | 0.0 | THSD7B | no | 19_39597358_G_A (lifespan-specific) |
| 19_52988758_A_G | chr19:52,988,758-52,991,258 | G | 0.44 | 0.08 | 7.9 × 10<sup>−8</sup> | genome-wide | CACNB4 | 0.0 | CACNB4 | no | no |
| 21_12266588_A_G | chr21:12,258,843-12,266,618 | G | 0.16 | 0.03 | 3.3 × 10<sup>−6</sup> | suggestive | TMEM135 | 143.5 | intergenic: RAB38 (435297 bp) \| TMEM135 (143510 bp) | no | no |
| **Lead variant** | **Locus region** | **Effect allele** | **β (years)** | **SE (years)** | **p** | **Significance** | **Nearest gene** | **Distance to nearest gene (kb)** | **Genes in locus region (RefSeq 105)** | **Overlaps a breed-weight locus** | **Overlapping breed-lifespan locus (unadjusted)** |
| 21_45834740_A_AT | chr21:45,834,187-45,835,914 | AT | 0.21 | 0.04 | 2.1 × 10<sup>−6</sup> | suggestive | LUZP2 | 92.2 | intergenic: CCDC179 (1393205 bp) \| LUZP2 (91040 bp) | no | no |
| 22_46946934_G_T | chr22:46,940,789-46,969,699 | T | 0.44 | 0.09 | 1.2 × 10<sup>−6</sup> | suggestive | HS6ST3 | 310.7 | intergenic: UGGT2 (349194 bp) \| HS6ST3 (287942 bp) | no | no |
| 23_31224813_G_A | chr23:31,224,813-31,281,501 | A | 0.43 | 0.09 | 3.9 × 10<sup>−6</sup> | suggestive | ANAPC13 | 16.9 | ANAPC13, CEP63 | no | no |
| 24_16989780_C_CA | chr24:16,989,780-16,990,049 | CA | 0.24 | 0.05 | 2.2 × 10<sup>−6</sup> | suggestive | PRNP | 127.5 | intergenic: PRNP (127461 bp) \| ADRA1D (223566 bp) | no | no |
| 25_25694485_T_C | chr25:25,694,485-25,721,705 | C | 0.43 | 0.09 | 1.9 × 10<sup>−6</sup> | suggestive | CBD131 | 193.1 | LOC486076 | no | no |
| 26_24247930_G_A | chr26:24,247,930-24,396,272 | A | 0.84 | 0.17 | 5.6 × 10<sup>−7</sup> | suggestive | PIK3IP1 | 0.0 | PIK3IP1, PATZ1, LOC106557811,DRG1, EIF4ENIF1 | no | 26_24247930_G_A (lifespan-specific) |
| 28_11147463_T_G | chr28:11,046,806-11,147,463 | G | 0.31 | 0.06 | 2.2 × 10<sup>−7</sup> | genome-wide | CRTAC1 | 0.0 | GOLGA7B, CRTAC1 | no | no |
| 28_15254395_C_T | chr28:15,042,524-15,254,395 | T | 0.7 | 0.15 | 2.0 × 10<sup>−6</sup> | suggestive | WBP1L | 0.0 | SUFU,TRIM8, ARL3,SFXN2, WBP1L | no | no |
| 29_34073415_T_TA | chr29:32,068,516-34,073,415 | TA | 0.44 | 0.09 | 1.1 × 10<sup>−6</sup> | suggestive | MMP16 | 0.0 | PSKH2, ATP6V0D2, SLC7A13, WWP1, RMDN1, CPNE3, CNGB3, CNBD1, LOC608680, MMP16 | no | no |
| 33_28057715_A_G | chr33:28,057,056-28,057,715 | G | 0.18 | 0.04 | 1.5 × 10<sup>−6</sup> | suggestive | HEG1 | 13.3 | intergenic: HEG1 (12677 bp) \| SLC12A8 (27630 bp) | no | no |
| 34_20301358_T_C | chr34:20,296,548-20,301,906 | C | 0.16 | 0.03 | 1.3 × 10<sup>−6</sup> | suggestive | BCL6 | 165.7 | intergenic: BCL6 (160843 bp) \| LPP (236674 bp) | no | no |

**Table S6.**
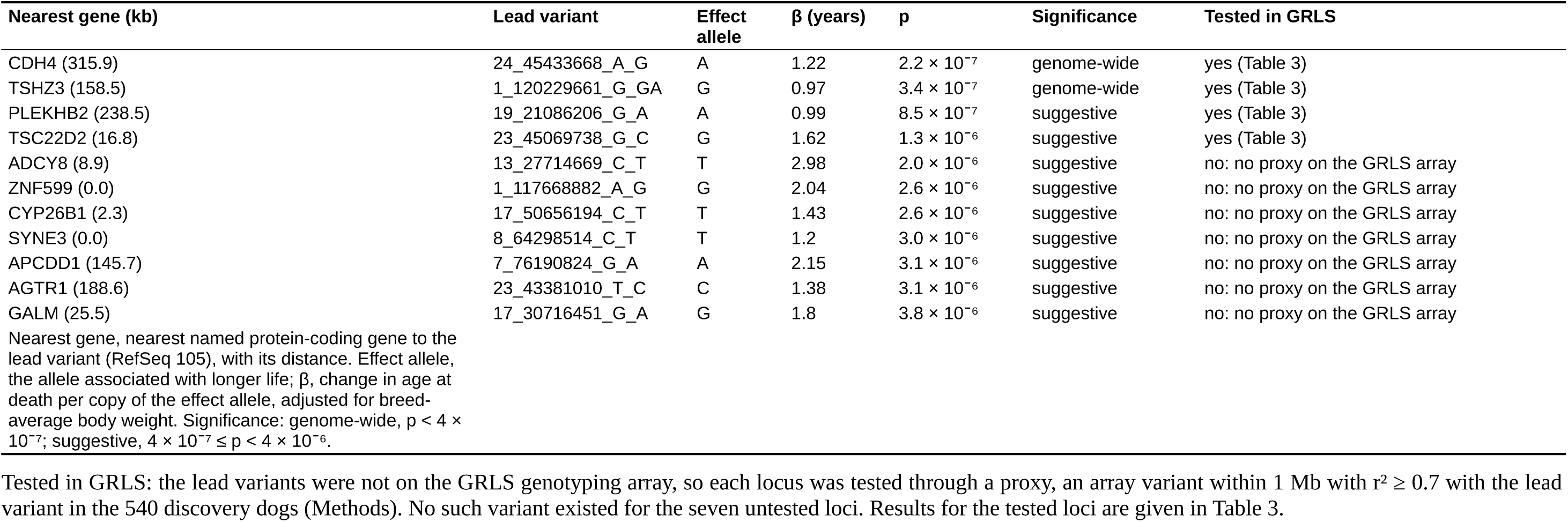
Loci associated with age at death in the Cani-DNA discovery cohort.

| Nearest gene (kb) | Lead variant | Effect allele | β (years) | p | Significance | Tested in GRLS |
|---|---|---|---|---|---|---|
| CDH4 (315.9) | 24_45433668_A_G | A | 1.22 | 2.2 × 10<sup>−7</sup> | genome-wide | yes (Table 3) |
| TSHZ3 (158.5) | 1_120229661_G_GA | G | 0.97 | 3.4 × 10<sup>−7</sup> | genome-wide | yes (Table 3) |
| PLEKHB2 (238.5) | 19_21086206_G_A | A | 0.99 | 8.5 × 10<sup>−7</sup> | suggestive | yes (Table 3) |
| TSC22D2 (16.8) | 23_45069738_G_C | G | 1.62 | 1.3 × 10<sup>−6</sup> | suggestive | yes (Table 3) |
| ADCY8 (8.9) | 13_27714669_C_T | T | 2.98 | 2.0 × 10<sup>−6</sup> | suggestive | no: no proxy on the GRLS array |
| ZNF599 (0.0) | 1_117668882_A_G | G | 2.04 | 2.6 × 10<sup>−6</sup> | suggestive | no: no proxy on the GRLS array |
| CYP26B1 (2.3) | 17_50656194_C_T | T | 1.43 | 2.6 × 10<sup>−6</sup> | suggestive | no: no proxy on the GRLS array |
| SYNE3 (0.0) | 8_64298514_C_T | T | 1.2 | 3.0 × 10<sup>−6</sup> | suggestive | no: no proxy on the GRLS array |
| APCDD1 (145.7) | 7_76190824_G_A | A | 2.15 | 3.1 × 10<sup>−6</sup> | suggestive | no: no proxy on the GRLS array |
| AGTR1 (188.6) | 23_43381010_T_C | C | 1.38 | 3.1 × 10<sup>−6</sup> | suggestive | no: no proxy on the GRLS array |
| GALM (25.5) | 17_30716451_G_A | G | 1.8 | 3.8 × 10<sup>−6</sup> | suggestive | no: no proxy on the GRLS array |

Nearest gene, nearest named protein-coding gene to the lead variant (RefSeq 105), with its distance. Effect allele, the allele associated with longer life; β, change in age at death per copy of the effect allele, adjusted for breed-average body weight. Significance: genome-wide, p < 4 × 10<sup>−7</sup>; suggestive, 4 × 10<sup>−7</sup> ≤ p < 4 × 10<sup>−6</sup>.
Tested in GRLS: the lead variants were not on the GRLS genotyping array, so each locus was tested through a proxy, an array variant within 1 Mb with r<sup>2</sup> ≥ 0.7 with the lead variant in the 540 discovery dogs (Methods). No such variant existed for the seven untested loci. Results for the tested loci are given in Table 3.

**Table S7.** Gene sets over-represented among the genes at the loci of each GWAS.

| GWAS | Library | Term | Overlap | p | Adjusted p | Loci contributing genes | Genes |
|---|---|---|---|---|---|---|---|
| Breed-average body weight | Aging Perturbations from GEO down | Mouse striatum 6 months vs 21 months GDS4153 aging:78 | 17/283 | 4.2 × 10<sup>−5</sup> | 0.012 | 14 | PTH2R, JUN, IPCEF1, TRHR, GPR101, IRS4, OTOF, TMEM74, GABRA5, SPON1, SLITRK1, OLFM3, SATB2, VIP, RRAS2, CACNA1G, DUSP6 |
| Breed-average body weight | Aging Perturbations from GEO down | Mouse liver 6 months vs 26 months GSE20425 aging:365 | 14/222 | 1.2 × 10<sup>−4</sup> | 0.017 | 8 | HMGA2, FAM126B, ABCC3, NDUFB3, GLI1, CENPA, FZD7, VIP, CRYGA, MAPRE3, SHMT2, GSTM2, NOP58, SLC5A6 |
| Breed-average body weight | Aging Perturbations from GEO down | Rat femur 26 weeks vs 52 weeks GDS509 aging:269 | 15/269 | 2.6 × 10<sup>−4</sup> | 0.025 | 11 | JUN, CGREF1, CTBP1, NDUFA4L2, RPL22, ENO1, NDUFB3, PCNA, COL11A1, CAPN6, TMEM214, RPS3A, GSTM2, RPS27A, CDK4 |
| Breed-average body weight | Aging Perturbations from GEO down | Mouse retina 3 months vs 20 months GSE17423 aging:335 | 14/247 | 3.6 × 10<sup>−4</sup> | 0.025 | 8 | ICMT, LUC7L3, CLSTN3, R3HDM2, CTBP1, MPP4, TMEM237, SUMO1, MIP, KIF5A, KPNA2, PDE6B, HMGN2, GAS6 |
| Breed-average body weight | Aging Perturbations from GEO down | Rat diaphysealfemoralcallus 6 weeks vs 26 weeks GDS1995 aging:115 | 16/319 | 5.4 × 10<sup>−4</sup> | 0.031 | 12 | EMB, RPL22, GNAI3, NACA, PCNA, COL11A1, OXR1, IGF1, CAPN6, COPB1, RPS3A, BTG1, GSTM5, GSTM2, RPS27A, CDK4 |
| Breed-average body weight | Aging Perturbations from GEO down | Rat liver 12 months vs 24 months GSE11097 aging:194 | 14/269 | 8.3 × 10<sup>−4</sup> | 0.04 | 8 | XDH, INHBE, AGMO, STAC3, SCP2, GLS2, NDUFB3, CTDSP2, HSD17B6, IGF1, RDH16, TSPAN31, C8B, GSTM2 |
| Breed-average body weight | Aging Perturbations from GEO down | Mouse retinal pigment epithelium / choroid 7 weeks vs 64 weeks GSE38671 aging:213 | 15/316 | 0.0014 | 0.044 | 8 | OST4, CRYGD, FAM126B, TRA2B, ANAPC13, CRYGC, ENO1, LRPAP1, PDE6B, CRYGA, HADHA, CRYGB, KIF5A, ATP5B, HMGN2 |
| Breed-average body weight | Aging Perturbations from GEO down | Rat diaphysealfemoralcallus 6 weeks vs 52 weeks GDS1995 aging:116 | 12/221 | 0.0013 | 0.044 | 9 | EMB, GNAI3, NACA, PCNA, COL11A1, RPS29, CAPN6, RPS3A, BTG1, GSTM2, RPS27A, CDK4 |
| Breed-average body weight | Aging Perturbations from GEO down | Rat liver 6 months vs 18 months GSE11097 aging:197 | 14/280 | 0.0012 | 0.044 | 9 | XDH, AOX1, INHBE, STAC3, SCP2, GLS2, PTGES3, KHK, RPS29, IGF1, SLC26A1, ATP5B, HMGN2, DUSP6 |
| Breed-median lifespan | Aging Perturbations from GEO down | Mouse striatum 6 months vs 21 months GDS4153 aging:78 | 12/283 | 3.7 × 10<sup>−5</sup> | 0.0098 | 11 | JUN, ISLR, IRS4, OTOF, SPON1, GABRA5, SLITRK1, OLFM3, RRAS2, DLX2, CACNA1G, DUSP6 |
| Breed-median lifespan | Aging Perturbations from GEO down | Rat thoracicaorta 6 months vs 15 months GDS3077 aging:122 | 12/302 | 7.0 × 10<sup>−5</sup> | 0.0098 | 9 | XDH, JUN, HNRNPA2B1, ENO1, HADHB, GNAI3, EEF1A1, RPS2, HADHA, COL3A1, GSTM5, RAB10 |
| Breed-median lifespan | WikiPathways 2024 Human | Mitochondrial Fatty Acid Oxidation Disorders WP5123 | 4/19 | 7.0 × 10<sup>−5</sup> | 0.011 | 3 | ACSL4, HADHB, HADHA, SCP2 |
| Breed-median lifespan | WikiPathways 2024 Human | NRF2 Pathway WP2884 | 9/141 | 5.2 × 10<sup>−5</sup> | 0.011 | 4 | SLC2A5, GSTM4, ABCC3, GSTM1, SLC2A7, GSTM3, GSTM5, GSTM2, SLC5A6 |
| Breed-median lifespan | Reactome Pathways 2024 | Acyl Chain Remodeling of CL | 3/6 | 3.4 × 10<sup>−5</sup> | 0.024 | 2 | LCLAT1, HADHB, HADHA |
| Breed-median lifespan | Reactome Pathways 2024 | Glutathione Conjugation | 5/37 | 7.7 × 10<sup>−5</sup> | 0.028 | 1 | GSTM4, GSTM1, GSTM3, GSTM5, GSTM2 |
| Breed-median lifespan adjusted for body weight | Human Phenotype Ontology | Glomerulonephritis (HP:0000099) | 3/12 | 2.9 × 10<sup>−5</sup> | 0.0024 | 1 | C1QA, C1QB, C1QC |
| Breed-median lifespan adjusted for body weight | Human Phenotype Ontology | Systemic lupus erythematosus (HP:0002725) | 3/11 | 2.2 × 10<sup>−5</sup> | 0.0024 | 1 | C1QA, C1QB, C1QC |
| Breed-median lifespan adjusted for body weight | GO Biological Process 2026 | Cell Junction Disassembly (GO:0150146) | 3/9 | 7.5 × 10<sup>−6</sup> | 0.0026 | 1 | C1QA, C1QB, C1QC |
| Breed-median lifespan adjusted for body weight | GO Biological Process 2026 | Complement Activation, Classical Pathway (GO:0006958) | 3/9 | 7.5 × 10<sup>−6</sup> | 0.0026 | 1 | C1QA, C1QB, C1QC |
| Breed-median lifespan adjusted for body weight | Human Phenotype Ontology | Complement deficiency (HP:0004431) | 3/15 | 6.0 × 10<sup>−5</sup> | 0.0033 | 1 | C1QA, C1QB, C1QC |
| Breed-median lifespan adjusted for body weight | GO Biological Process 2026 | Humoral Immune Response Mediated by Circulating Immunoglobulin (GO:0002455) | 3/12 | 1.9 × 10<sup>−5</sup> | 0.0034 | 1 | C1QA, C1QB, C1QC |
| Breed-median lifespan adjusted for body weight | GO Biological Process 2026 | Synapse Pruning (GO:0098883) | 3/12 | 1.9 × 10<sup>−5</sup> | 0.0034 | 1 | C1QA, C1QB, C1QC |
| Breed-median lifespan adjusted for body weight | Human Phenotype Ontology | Abnormality of complement system (HP:0005339) | 3/20 | 1.5 × 10<sup>−4</sup> | 0.0061 | 1 | C1QA, C1QB, C1QC |
| Breed-median lifespan adjusted for body weight | Human Phenotype Ontology | Nephritis (HP:0000123) | 3/24 | 2.6 × 10<sup>−4</sup> | 0.0086 | 1 | C1QA, C1QB, C1QC |
| Breed-median lifespan adjusted for body weight | Aging Perturbations from GEO up | Mouse brain 8 weeks vs 104 weeks GSE20411 aging:311 | 6/192 | 6.9 × 10<sup>−5</sup> | 0.016 | 4 | C1QA, C1QC, RPL26, PRNP, C1QB, RPS10 |
| Breed-median lifespan adjusted for body weight | Aging Perturbations from GEO up | Mouse brain 8 weeks vs 104 weeks GSE20411 aging:310 | 6/241 | 2.4 × 10<sup>−4</sup> | 0.019 | 4 | C1QA, C1QC, RPL26, PRNP, C1QB, RPS10 |
| Breed-median lifespan adjusted for body weight | Aging Perturbations from GEO up | Mouse liver 5 months vs 24 months GSE25325 aging:288 | 7/335 | 2.1 × 10<sup>−4</sup> | 0.019 | 5 | C1QA, C1QC, RPL26, ANAPC13, PRNP, C1QB, RPS10 |
| Breed-median lifespan adjusted for body weight | WikiPathways 2024 Human | Complement Activation WP545 | 3/23 | 2.3 × 10<sup>−4</sup> | 0.024 | 1 | C1QA, C1QB, C1QC |
| Breed-median lifespan adjusted for body weight | Aging Perturbations from GEO up | Human meniscus 40 years vs 55 years GSE45233 aging:209 | 6/280 | 5.3 × 10<sup>−4</sup> | 0.031 | 4 | C1QB, RPSAP9, PRNP, CPNE3, LMO4, RPS10 |
| Breed-median lifespan adjusted for body weight | GO Biological Process 2026 | Imitative Learning (GO:0098596) | 2/6 | 3.0 × 10<sup>−4</sup> | 0.035 | 2 | NRXN1, HTT |
| Breed-median lifespan adjusted for body weight | GO Biological Process 2026 | Vocal Learning (GO:0042297) | 2/6 | 3.0 × 10<sup>−4</sup> | 0.035 | 2 | NRXN1, HTT |
| Breed-median lifespan adjusted for body weight | Aging Perturbations from GEO up | Mouse neocortex 5 months vs 30 months GSE13120 aging:334 | 6/319 | 0.0011 | 0.041 | 4 | C1QA, C1QC, ARL3, CBX3, C1QB, DRG1 |
| Breed-median lifespan adjusted for body weight | Aging Perturbations from GEO up | Mouse tumor 68 days vs 143 days GSE66414 aging:143 | 6/312 | 9.4 × 10<sup>−4</sup> | 0.041 | 3 | C1QA, C1QC, TRIM8, C1QB, WBP1L, RPS10 |
| Breed-median lifespan adjusted for body weight | WikiPathways 2024 Human | Microglia Pathogen Phagocytosis Pathway WP3937 | 3/40 | 0.0012 | 0.042 | 1 | C1QA, C1QB, C1QC |
| Breed-median lifespan adjusted for body weight | WikiPathways 2024 Human | Oxidative Damage Response WP3941 | 3/40 | 0.0012 | 0.042 | 1 | C1QA, C1QB, C1QC |
| Age at death | WikiPathways 2024 Human | Envelope Proteins And Potential Roles In EDMD Physiopathology WP4535 | 3/46 | 4.4 × 10<sup>−5</sup> | 0.0025 | 3 | BANF1, SYNE3, ADCY8 |
| Age at death | MGI Mammalian Phenotype Level 4 2024 | Cerebellum Vermis Hypoplasia MP:0000866 | 2/10 | 7.0 × 10<sup>−5</sup> | 0.015 | 1 | ZIC1, ZIC4 |
| Age at death | MGI Mammalian Phenotype Level 4 2024 | Abnormal Cerebellum Vermis Morphology MP:0000864 | 2/17 | 2.1 × 10<sup>−4</sup> | 0.023 | 1 | ZIC1, ZIC4 |
| Age at death | MGI Mammalian Phenotype Level 4 2024 | Cerebellum Hypoplasia MP:0000851 | 2/23 | 3.9 × 10<sup>−4</sup> | 0.028 | 1 | ZIC1, ZIC4 |
| Age at death | Human Phenotype Ontology | Wide cranial sutures (HP:0010537) | 2/17 | 5.9 × 10<sup>−4</sup> | 0.042 | 2 | BANF1, AGTR1 |
| Age at death | MGI Mammalian Phenotype Level 4 2024 | Abnormal Cerebellar Foliation MP:0000857 | 2/35 | 9.0 × 10<sup>−4</sup> | 0.049 | 1 | ZIC1, ZIC4 |

Terms significant at a Benjamini–Hochberg adjusted p < 0.05 in the over-representation analysis of genes at the loci of each GWAS (Methods).

